# Astrocytes regulate locomotion by orchestrating neuronal rhythmicity in the spinal network via potassium clearance

**DOI:** 10.1101/2022.07.07.498974

**Authors:** Tony Barbay, Emilie Pecchi, Myriam Ducrocq, Nathalie Rouach, Frédéric Brocard, Rémi Bos

**Affiliations:** Institut de Neurosciences de la Timone (UMR7289), Aix-Marseille Université and CNRS, Marseille, France; Center for Interdisciplinary Research in Biology, Collège de France, CNRS, INSERM, Labex Memolife, Université PSL, Paris, France

**Keywords:** Locomotion, Spinal cord, Astrocytes, Neuronal Oscillations, Potassium uptake, Kir4.1

## Abstract

Neuronal rhythmogenesis in the spinal cord is correlated with variations in extracellular K^+^ levels ([K^+^]_e_). Astrocytes play important role in[K^+^]_e_ homeostasis and compute neuronal information. Yet it is unclear how neuronal oscillations are regulated by astrocytic K^+^ homeostasis. Here we identify the astrocytic inward-rectifying K^+^ channel Kir4.1 (a.k.a. *Kcnj10*) as a key molecular player for neuronal rhythmicity in the spinal central pattern generator (CPG). By combining two-photon calcium imaging with electrophysiology, immunohistochemistry and genetic tools, we report that astrocytes display Ca^2+^ transients before and during oscillations of neighbouring neurons. Inhibition of astrocytic Ca^2+^ transients with BAPTA decreases the barium-sensitive Kir4.1 current responsible of K^+^ clearance. Finally, we show in mice that Kir4.1 knockdown in astrocytes progressively prevents neuronal oscillations and alters the locomotor pattern resulting in lower motor performances in challenging tasks. These data identify astroglial Kir4.1 channels as key regulators of neuronal rhythmogenesis in the CPG driving locomotion.

**Significance statement:** Despite decades of research, the cellular mechanisms responsible of the synchronized rhythmic oscillations driving locomotion remain elusive. To gain insight into the function of the spinal locomotor network, numerous studies have characterized diverse classes of locomotor-related neurons to determine their role in generating rhythmic movements during locomotion. In contrast, studies investigating non-neuronal components of the spinal cord are sparse. Our study represents a significant breakthrough by identifying astrocytic K^+^ uptake as a key regulator of neuronal rhythmicity synchronization and locomotor pattern at the cellular, microcircuit and system levels. These data provide mechanistic insights into the neuroglial dialogue at play during rhythmogenesis and point to a novel astroglial target for restoring normal neuronal network excitability in brain disorders and neurodegenerative diseases.

## INTRODUCTION

Mammalian locomotion relies on repeated sequences of muscle contractions triggered by a rhythmic central pattern generator (CPG) network, which is mainly located in the ventromedial part of the lumbar spinal cord (Grillner and El Manira, 2020; Kiehn, 2016). Activation of the spinal CPG network is triggered by descending inputs releasing neuromodulators such as glutamate, serotonin and dopamine. Several populations of CPG interneurons - V0, V2a and Hb9 –display conditional rhythmic oscillations in response to neuromodulators (Grillner and El Manira, 2020; Kiehn, 2016). No single population of ventromedial interneurons appears to act alone or to orchestrate the entire CPG pattern. The spinal rhythmogenesis emerges from the synchronization of individual neurons with intrinsic bursting properties (Brocard et al., 2010). Despite decades of research, the functional organization of the locomotor CPG remains still unclear.

In the spinal locomotor CPG, physiological variation in extracellular K^+^ ([K^+^]_e_) coincides with the emergence of the oscillatory pattern of intrinsic bursting neurons (Brocard et al., 2013), and thus powers up the motor output (Bos et al., 2018; Bracci, 1998). At rest, [K^+^]_e_ is kept close to 3 mM in serum levels (Kofuji and Newman, 2004; Verkhratsky et al., 2018). The maintenance of K^+^ homeostasis is one of the crucial supportive functions mediated by astrocytes (Ben Haim et al., 2015; Sibille et al., 2015). In addition, to display a high resting K^+^ conductance that facilitates the uptake of neuron released K^+^, astrocytes have a hyperpolarized resting membrane potential (RMP). Both parameters rely on the activity of the weakly inwardly rectifying Kir4.1 K^+^ channel, which in the nervous system is expressed exclusively in glial cells, with the highest expression in astrocytes (Kelley et al., 2018). Astrocytic Kir4.1 channels have been described in many CNS regions, including the spinal cord (Olsen et al., 2006; Ransom and Sontheimer, 1995). Studies using genetic, pharmacological and modeling approaches in the brain networks identified Kir4.1 as the main astrocytic inwardly rectifying K^+^ channel regulating [K^+^]_e_ and thus influencing neuronal excitability (Nwaobi et al., 2016; Sibille et al., 2015).

Astrocytes display ramified processes enwrapping synapses, placing them in an ideal position to (i) sense synaptic activity (Panatier et al., 2011) and (ii) adjust network activity by releasing gliotransmitters (Araque et al., 2014; Savtchouk and Volterra, 2018). Rhythmic oscillations are strongly affected in the brain networks when gliotransmission is genetically or pharmacologically modified (Lee et al., 2014; Sheikhbahaei et al., 2018). In the spinal motor network, the locomotor oscillatory rhythm is correlated with enhanced calcium (Ca^2+^) transients in astrocytes (Broadhead and Miles, 2020), which likely triggers ATP release (Witts et al., 2015), modulating excitatory synaptic transmission (Carlsen and Perrier, 2014). Thus, growing evidence point to the contribution of gliotransmission in modulating neuronal rhythmogenesis (Montalant et al., 2021). Astrocytes can also respond to neurotransmitters and neuromodulators (Paukert et al., 2014; Rosa et al., 2015) by modifying [K^+^]_e_, which greatly impact brain oscillations (Bellot-Saez et al., 2017; Ding et al., 2016; Sibille et al., 2015). However, the mechanisms underlying the contribution of astrocytic K^+^ homeostasis to rhythmogenesis remain incompletely understood.

Here we investigated whether astrocyte K^+^ uptake modulates neuronal oscillations patterns in the spinal CPG network. We demonstrate that most of the ventromedial astrocytes are active during neuronal oscillations, and that the inwardly-rectifying Kir4.1 channels are crucial for maintaining neuronal oscillations, which ultimately influence the locomotor activity. This study places astrocytes as central elements of the spinal cord locomotor CPG.

## RESULTS

### Astrocytes respond to changes in extracellular ions underlying neuronal oscillations in the spinal CPG

Astrocytes from the ventromedial part of upper lumbar segments (L1–L2), the main locus of the locomotor CPG (El Manira, 2014; Kiehn, 2016), were recorded from transgenic mice expressing GFP under the control of the astrocytic *Aldh1L1* promoter (Tsai et al., 2012) (Fig 1A). This spinal region contains interneurons endowed with two types of inherent membrane oscillations at a frequency range similar to stepping rhythms: (i) one is TTX-sensitive dependent on the persistent sodium current (INaP) (Tazerart et al., 2008) and triggered by a rise in [K^+^]_e_ with a concomitant decrease in extracellular Ca^2+^ levels ([Ca^2+^]_e_) (Brocard et al., 2013), (ii) the other is TTX-insensitive dependent on the Ca^2+^ currents and triggered by a neuromodulator cocktail composed of 5-HT, dopamine and NMDA, in presence of the voltage-dependent sodium channels blocker, tetrodotoxin (TTX) (Masino et al., 2012; Wilson, 2005; Ziskind-Conhaim et al., 2008). This neuromodulator cocktail mimics the effect of descending inputs from the hindbrain for inducing a locomotor rhythmicity (Grillner and El Manira, 2020). In recording conditions required to generate bursting cells, all astrocytes displayed a reversible depolarisation of their membrane potentials: (i) when [K^+^]_e_ was increased to 6mM and [Ca^2+^]_e_ decreased to 0.9mM (Fig.1 B-C) or (ii) when the neuromodulator cocktail was applied in presence of TTX (Fig. 1 D-E). We observed that in absence of neuronal activity, neuromodulators increased [K^+^]_o_ to ∼6mM (Fig. S1). To explore the temporal dynamics of the astrocytic response compared to neuronal oscillations, we performed dual recordings of a ventromedial Hb9 GFP+ interneuron which is known to display the two types of bursting properties (Tazerart et al., 2008; Ziskind-Conhaim et al., 2008) and an adjacent astrocyte from the upper lumbar segments (L1-L2) (Fig. 1F). In 13 pairs, astrocytic depolarization in response to the TTX-sensitive [K^+^]_e_/[Ca^2+^]_e_ variations (Fig. 1 G) or the TTX-insensitive neuromodulator cocktail (Fig. 1H) either preceded (69%) or followed (31%) the onset of the neuronal rhythmic bursting of the adjacent GFP+ pacemaker interneurons (Fig. 1I), suggesting that astrocytes play an active role in spinal rhythmogenesis.

**Figure 1.**
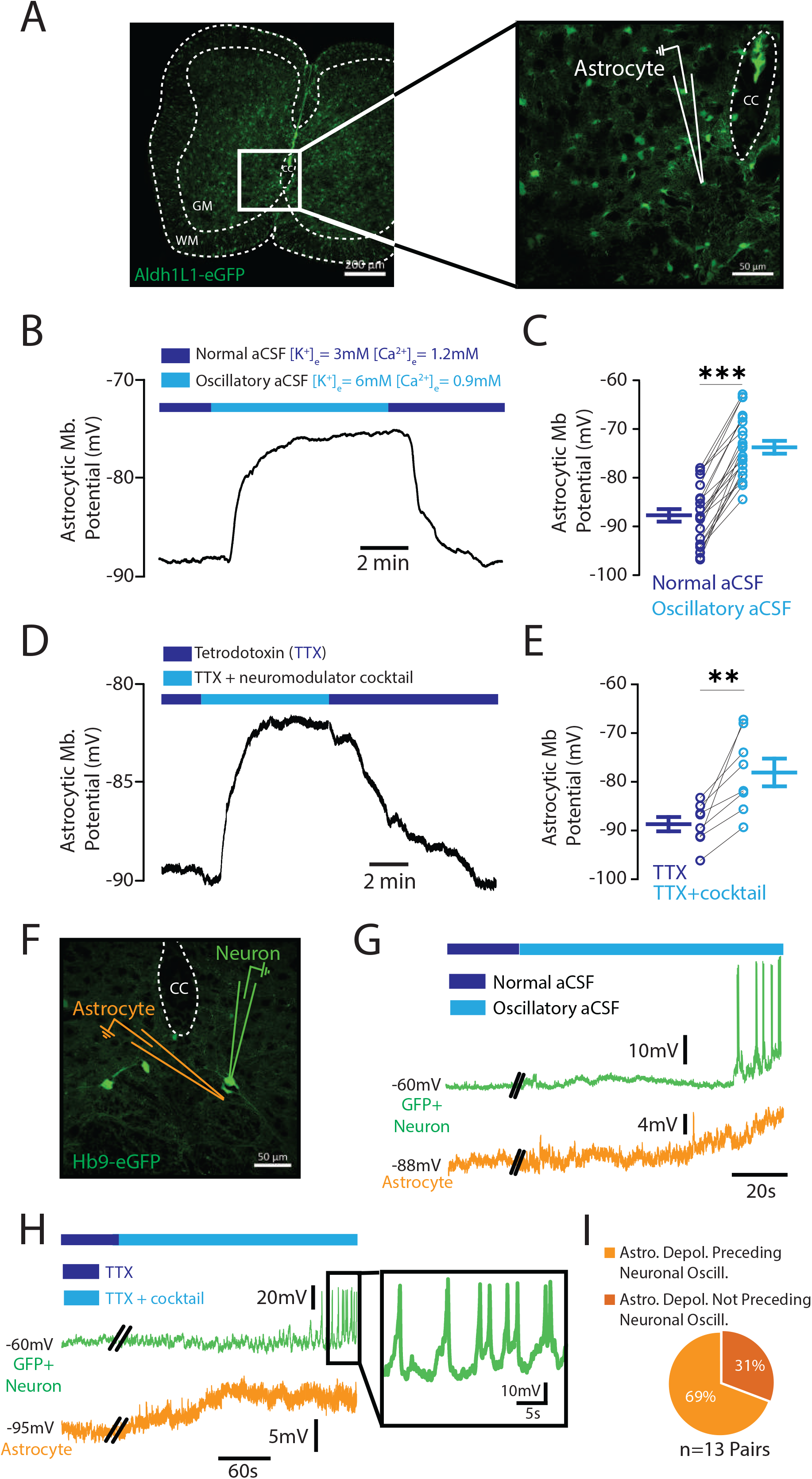
Astrocytes respond to variations in [K^+^]_e_ and [Ca^2+^]_e_ related to neuronal oscillations in the spinal CPG. **A**. Confocal image of endogenous GFP from a lumbar slice of a P8 *Aldh1L1-eGFP* mouse. White inset represents the patch-clamp recording area of the ventromedial GFP+ astrocytes. The recording glass microelectrode is in white. CC stands for central canal, GM for grey matter and WM for white matter. **B**. Representative voltage trace of a GFP+ astrocyte in response to [K^+^]_e_ rise to 6mM and [Ca^2+^]_e_ decrease to 0.9mM. C. Quantification of the effect of the [K^+^]_e_ and [Ca^2+^]_e_ variations on the astrocytic membrane potential (n = 22 astrocytes from 16 mice). **D**. Representative voltage trace of a GFP+ astrocyte in response to bath application of a neuromodulator cocktail (dopamine, serotonin, and NMDA) in presence of TTX. **E**. Quantification of the effect of the neuromodulator cocktail on the astrocytic membrane potential (n = 8 astrocytes from 4 mice). **F**. Confocal image of the endogenous GFP from the ventromedial part of a lumbar slice of a P12 Hb9-eGFP mouse. The green and orange glass microelectrodes represent the dual recording of a GFP+ interneuron and an adjacent astrocyte, respectively. CC for central canal. **G-H**. The [K^+^]_e_ and [Ca^2+^]_e_ variations (G, light blue) or the bath application of the neuromodulator cocktail in presence of TTX (H, light blue) induces bursting in Hb9 GFP+ interneurons (green traces) preceded by a depolarization of the neighbouring astrocyte (orange traces). The black inset in H represents the TTX-insensitive neuronal oscillations. **I**. Proportion of the astrocytic membrane potential depolarization preceding (light orange) or not preceding (dark orange) the onset of neuronal bursting (n=13 pairs from 8 mice). **P < 0.01; ***P < 0.001 (two-tailed Wilcoxon paired test for C and E). Mean ± SEM. For detailed P values, see Source data file.

### Enhanced astrocytic Ca^2+^ transients occur during neuronal oscillations in the ventral spinal cord

To better locate and characterize the dialogue between neurons and astrocytes during spinal oscillations, we performed two-photon Ca^2+^ imaging of neuronal and astrocyte signals in the locomotor CPG area. Mice were injected at birth with adeno-associated virus serotype 9 (AAV9) (Fig. S2A) designed to express GCaMp6f in astrocytic processes under the gfaABC1D promoter and jRGECO1a in neurons under the Syn.NES promoter (Fig. S2B). We simultaneously imaged the chromophores in lumbar slices (L1-L2) of post-natal mice (day 13-16) and first confirmed that neuronal signals were prevented by TTX, in contrast to the signals recorded astrocyte processes that remain intact (Fig. S2C-D).

We next explored the spatio-temporal dynamics of the astrocytic transients and compared it to the neuronal transients. In response to changes in extracellular K^+^ and Ca^2+^ levels or bath application of the neuromodulator cocktail in presence of TTX, neurons displayed Ca^2+^ oscillatory transients (0.16 ± 0.03 Hz and 0.15 ± 0.01 Hz, respectively) (Fig. 2A-B), which correlated with enhanced astrocytic Ca^2+^ transients in processes (Fig. 2A and 2C). Interestingly, we distinguished distinct subsets of astrocytic Ca^2+^ signals. Some astrocytes showed increased Ca^2+^transients before the onset of the neuronal Ca^2+^ oscillations, in contrast to others, which displayed Ca^2+^ transients during neuronal oscillations (Fig. 2A). Quantification showed that the occurrence of Ca^2+^ transients preceded the neuronal oscillations in ∼64% of astrocytes, whereas in ∼36% of them the Ca^2+^ transients occurred during neuronal oscillations (Fig. 2D). These data confirmed our electrophysiological dual recordings (Fig. 1I). We also observed that most of the astrocytes (∼70%) showing Ca^2+^ signal increases in response to ionic variations or perfusion of the neuromodulator cocktail were located in the ventromedial part of the spinal cord (Fig. 2E). Altogether, these data suggest that ventromedial astrocytes are functionally coupled with neuronal rhythmicity in the spinal CPG area.

**Figure 2.**
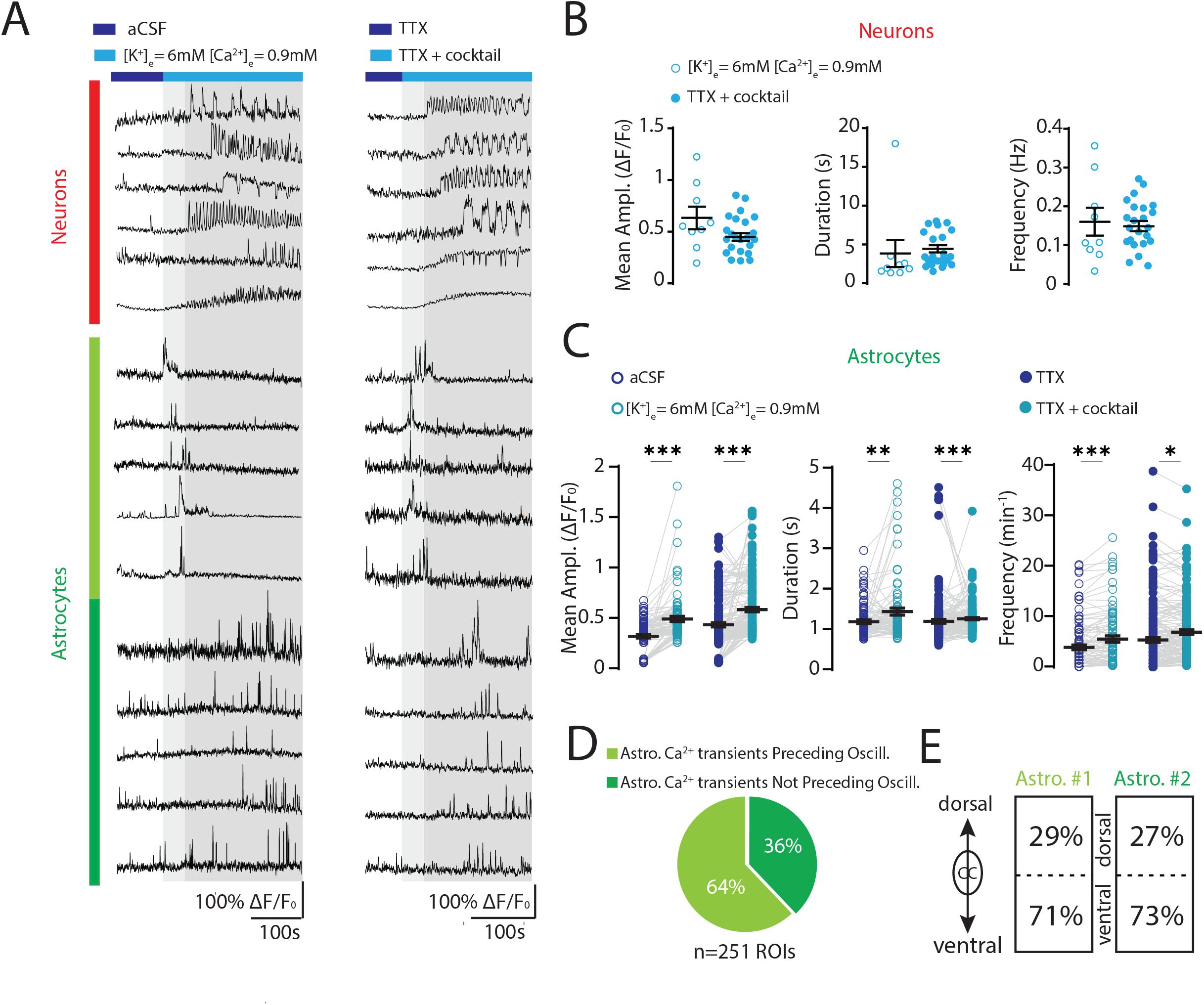
Increase in astrocytic Ca^2+^ transients is correlated to neuronal oscillations. **A. Top**. Examples of neuronal jRGECO1a Ca^2+^ transients (ΔF/Fo) oscillating in response to K^+^ rise and Ca^2+^ decrease (left) or in response to the neuromodulator cocktail in presence of TTX (right). *Middle, bottom*. Examples of the increase of the astrocytic Gcamp6f Ca^2+^ signals (ΔF/F0) before (middle) or during (bottom) the neuronal oscillations (top) from the same field of view. **B**. Quantification of the mean amplitude (left), duration and frequency (right) of the neuronal jRGECO1a Ca^2+^ transients in response to K^+^ rise and Ca^2+^ decrease (empty blue circles) or in response to monoaminergic cocktail in presence of TTX (filled blue circles) (n= 9 and 24 dendritic ROIs from 2 and 3 mice in the ionic variation condition and in the TTX+cocktail condition, respectively). **C**. Quantification of the mean amplitude (left), duration (middle) and frequency (right) of the astrocytic Gcamp6f Ca^2+^ signals in response to K^+^ rise and Ca^2+^ decrease (empty light blue circles) or in response to monoaminergic cocktail in presence of TTX (filled light blue circles) (n= 86 and 165 astrocytic ROIs from 2 and 3 mice in the ionic variation condition and in the TTX+cocktail condition, respectively). **D**. Proportion of astrocytic ROIs which display Ca^2+^ signals increase in amplitude and/or frequency before the onset of neuronal oscillations. **E**. Proportion of astrocytic ROIs which display Ca^2+^ signals increase above (dorsal) or below (ventral) the central canal (CC) in response to oscillatory conditions (ionic variations and neuromodulator cocktail). *P < 0.05, **P < 0.01, ***P < 0.001 (two-tailed Wilcoxon paired test for C). Mean ± SEM. For detailed P values, see Source data file.

### The astrocytic Ca^2+^ transients regulate the Kir4.1-mediated inwardly rectifying K^+^ current

Because astrocytic Ca^2+^ signals modulate [K^+^]_e_ in hippocampus (Wang et al., 2012a) and cerebellum (Wang et al., 2012b), we then investigated whether the increase in astrocytic Ca^2+^ transients related to neuronal oscillations in the spinal CPG area (Fig. 2) stimulates K^+^ uptake. We first recorded the passive membrane properties of the ventromedial GFP+ astrocytes from upper lumbar slices (Fig. 3A-B) and then isolated the Ba^2+^-sensitive Kir4.1 current underlying K^+^ uptake in standard artificial cerebro-spinal fluid (aCSF) (Fig. 3C-D). We then switched to oscillatory aCSF by increasing [K^+^]_e_ to 6mM and reducing [Ca^2+^]_e_ to 0.9mM, which leads to neuronal bursting (Fig. 1G and Fig. 2A-B) and intracellular astrocytic Ca^2+^ transients (Fig. 2A and 2C). In this condition, we observed that preventing the astrocytic Ca^2+^ transients with BAPTA (30mM) inside the patch pipette significantly reduced the barium (Ba^2+^)-sensitive current driven by Kir4.1 channels (Fig. 3 E-F). These data strongly suggest that the increase in intracellular Ca^2+^ from ventromedial astrocytes observed during neuronal oscillations modulates K^+^ uptake via Kir4.1 channels.

**Figure 3.**
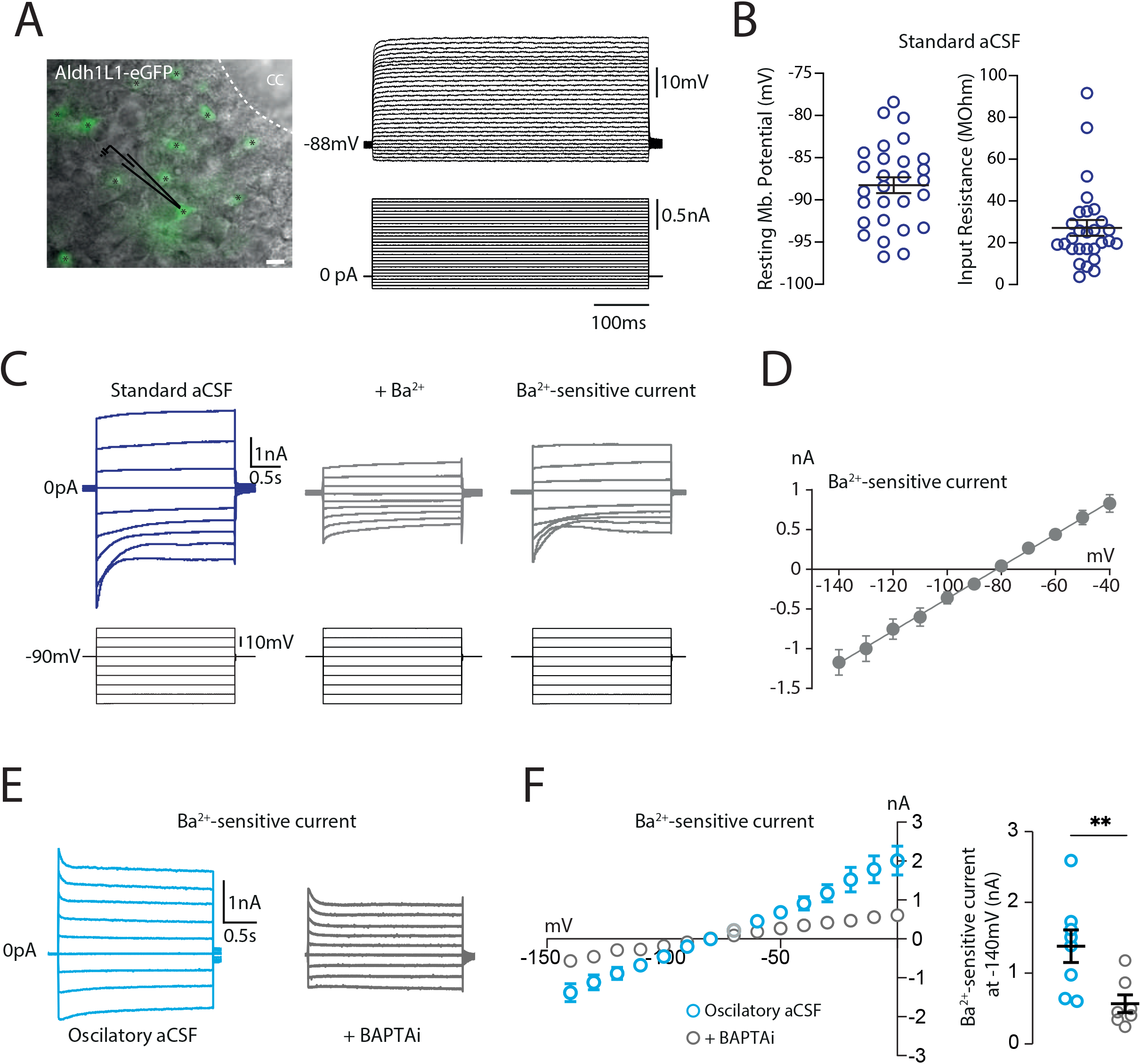
Spinal astrocytes display a Ba^2+^-sensitive Kir4.1 current modulated by intracellular Ca^2+^. **A**. *Left*. DIC image of a lumbar slice from a P8 *Aldh1L1-eGFP* mouse superimposed with the fluorescent GFP signal. The black stars indicate the GFP+ astrocytes. The recording glass pipette is represented in black. *Right*. Representative membrane potentials (top) of the targeted GFP+ astrocyte (left image) in response to incremental current pulses (500ms, Δ = 50pA). **B**. Resting membrane potential (RMP) (left) and input resistance (right) of lumbar GFP+ astrocytes recorded in standard aCSF ([K^+^]_e_ 3mM and [Ca^2+^]_e_ 1.2mM) (n = 28 astrocytes from n = 15 mice) **C**. Representative membrane currents (top) of one GFP+ astrocyte clamped at the RMP in response to incremental voltages pulses (bottom, 2.5s, -140 to -40mV, Δ = 10mV) before (left) or during (middle) application of Ba^2+^ (100 µM) in standard aCSF. **D**. The averaged I/V relationship of the Ba^2+^ -sensitive currents of GFP+ astrocytes (n = 17 astrocytes from 9 mice). **E**. Representative Ba^2+^ -sensitive currents of two astrocytes clamped at the RMP in response to incremental voltages pulses (2.5s, -140 to +0mV, Δ = 10mV) without (blue) or with (black) intracellular perfusion of BAPTA (30 mM) in oscillatory aCSF ([K^+^]_e_ 6mM and [Ca^2+^]_e_ 0.9mM). **F.** *Left*. The averaged I/V curves from lumbar astrocytes without (blue circles) or with (black circles) intracellular BAPTA perfusion in oscillatory aCSF. *Right*. Average effect of intracellular perfusion of BAPTA on the peak amplitude of the astrocytic Kir4.1 current at - 140mV (G). (n = 8 and 7 astrocytes from 3 and 4 mice for control and BAPTAi, respectively). **P<0.01 (two-tailed Mann–Whitney test for F). Mean ± SEM. For detailed P values, see Source data file.

### Kir4.1 channels are expressed in astrocytes from the spinal locomotor CPG region

We next investigated whether Kir4.1 channels are expressed in the spinal locomotor CPG. Using Hb9-eGFP mice which express GFP+ interneurons with intrinsic bursting properties (Tazerart et al., 2008; Ziskind-Conhaim et al., 2008), we found that ventromedial GFP+ interneurons were enwrapped by Kir4.1 labeling, which is diffusely expressed in the grey matter of the upper lumbar segments (Fig. S3A). To confirm that Kir4.1 channels in the locomotor CPG area are exclusively expressed in glial cells with a highest expression in astrocytes, we used the *Aldh1L1-eGFP* astrocyte-specific reporter mice (Cahoy et al., 2008). We examined co-expression of *Aldh1L1-eGFP* with immunolabeling of Kir4.1 channels and neuronal marker NeuN in lumbar slices. We observed that 100% of the GFP+ astrocytes expressed Kir4.1 channels contrary to NeuN+ cells, which lacked Kir4.1 staining (Fig. S3B-C).

### The Ba^2+^-sensitive inwardly rectifying K^+^ (Kir4.1) current is crucial for maintaining neuronal oscillations and locomotor-like activity in the CPG

We next studied the role of Kir4.1 channels in the lumbar rhythmogenesis. We first induced rhythmic oscillations in the GFP+ ventromedial interneurons from Hb9-eGFP mice in response to [K^+^]_e_/[Ca^2+^]_e_ variations (Fig. 4A) or bath perfusion of the neuromodulator cocktail in presence of TTX (Fig. 4B). In both conditions, pharmacological blockade of Kir4.1 channels with Ba^2+^ (100 µM) caused a shift in the spiking pattern from bursting to tonic firing in ∼70% of the TTX-sensitive neuronal oscillations (Fig. 4C) or resulted in a cessation of bursting activity (∼85%) of the TTX-insensitive neuronal oscillations (Fig. 4D). Interestingly, we noticed that before totally preventing the neuronal oscillations, Ba^2+^ progressively decreased the rhythmic bursting frequency accompanied with an increase in burst amplitude and duration in both conditions (Fig. S4). We next confirmed the Ba^2+^ effect on preventing neuronal oscillations at a larger scale by two-photon Ca^2+^ imaging of cell populations. We observed a progressive loss of Ca^2+^ oscillations in the vast majority of neurons in response to Ba^2+^ in presence of the oscillatory aCSF (n=25/28) or the neuromodulator cocktail (n=7/9) (Fig. 4E-F). Bath application of Ba^2+^ also induced a significant rise in amplitude of the astrocytic Ca^2+^ signal in both conditions (Fig. 4G).

**Figure 4.**
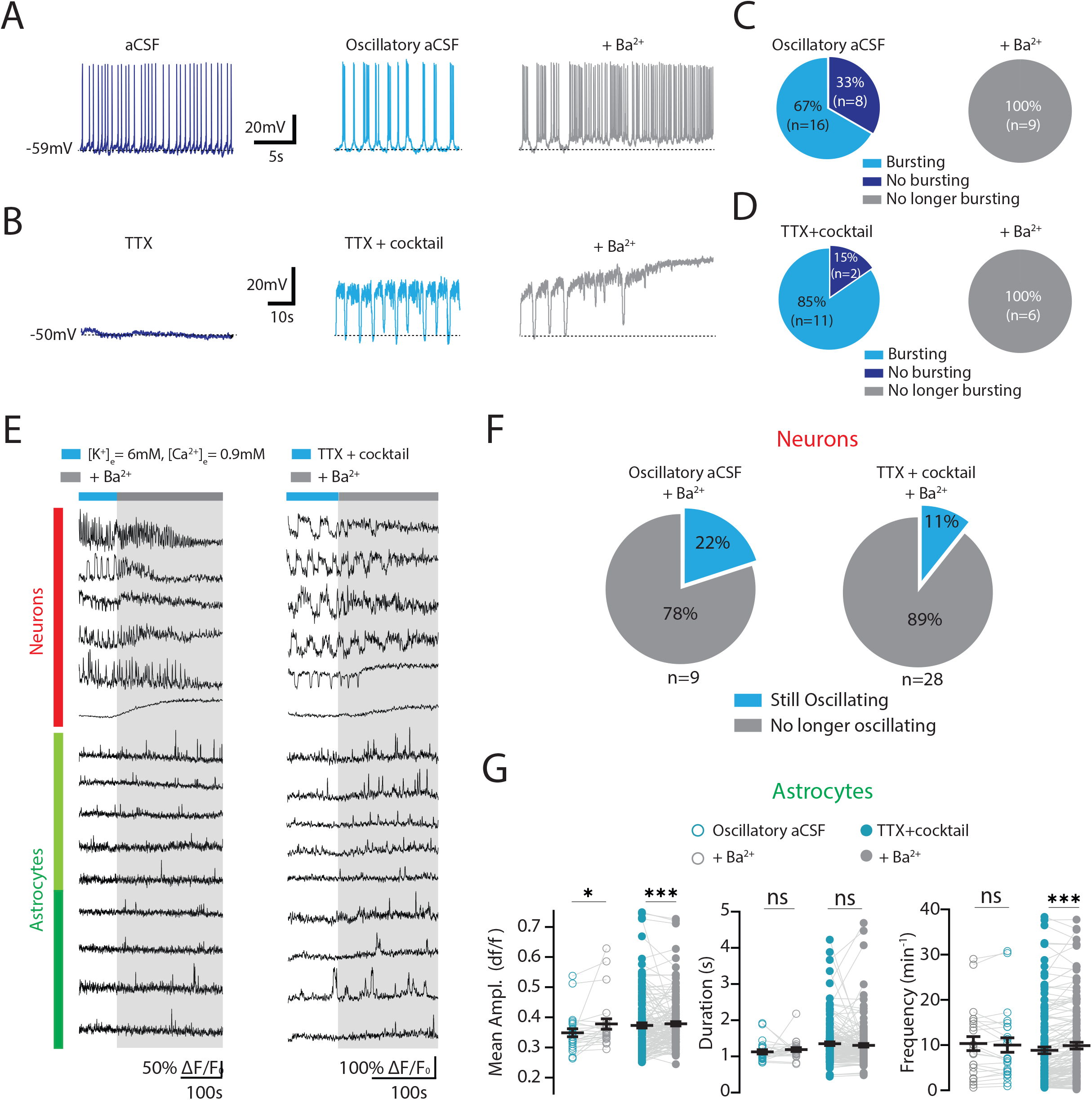
K^+^ homeostasis maintenance through Kir4.1 channels is crucial for neuronal oscillations. **A-B**. Representative voltage traces of a lumbar GFP+ ventromedial interneuron from a P6 Hb9-eGFP mouse before (left) and during (middle) ionic variations ([K^+^]_e_ = 6mM, [Ca^2+^]_e_ = 0.9mM) on which Ba^2+^ (100 µM) is added (right) (A) or in presence of TTX (left) and following application of the neuromodulator cocktail (middle) on which Ba^2+^ is added (right) (B). **C-D**. Quantification of the proportion of GFP+ interneurons oscillating in response to ionic variations (C, left) or to the neuromodulator cocktail (D, left) and following bath application of Ba^2+^ for more than 5 minutes (C-D, right). **E**. *Top*. Examples of neuronal jRGECO1a Ca signals (ΔF/Fo) before (blue) and during bath application of Ba^2+^ (grey) in presence of high [K^+^]_e_ and low [Ca^2+^]_e_ (left) or in presence of the neuromodulator cocktail + TTX (right). *Bottom*. Examples of astrocytic Gcamp6f Ca^2+^ signals (ΔF/F0) before (blue) or during (grey) application of Ba^2+^ in presence of high [K^+^]_e_ and low [Ca^2+^]_e_ (left) or in presence of TTX and the neuromodulator cocktail (right). **F**. Proportion of neuronal ROIs still oscillating or not in response to Ba^2+^ application (>5min) in presence of high [K^+^]_e_ and low [Ca^2+^]_e_ (left) or in presence of TTX and the neuromodulator cocktail (right). **G**. Effect of Ba^2+^ application (empty and filled grey circles) on the mean amplitude (left), duration (middle) and frequency (right) of the astrocytic Gcamp6f Ca^2+^ signals in presence of K^+^ rise and Ca^2+^ decrease (empty light blue circles) or in presence of the neuromodulator cocktail + TTX (filled light blue circles). Each circle represents one astrocytic ROI (n= 24 ROIs from 2 mice in the ionic variation condition and n= 157 ROIs from 3 mice in the TTX+cocktail condition) ns, no significance, *P < 0.05, ***P < 0.001 (two-tailed Wilcoxon test for G). Mean ± SEM. For detailed P values, see Source data file.

To confirm the functional role of the inwardly rectifying K^+^ (Kir4.1) current in the integrated rhythmogenesis, we investigated the contribution of Kir4.1 in the locomotor rhythm-generating network by using *ex vivo* whole-mount spinal cords. Since rostral lumbar segments (L1–L2) have a more powerful rhythmogenic capacity than the caudal ones, we recorded locomotor-like activities from the contralateral L2 ventral roots (Fig. 5A) in response to ionic variations (Fig. 5B) or bath application of a neuromodulator cocktail composed of N-methyl-DL aspartate (NMA) and 5 hydroxytryptamine (5-HT) (Fig. 5C). In both conditions, Ba^2+^ (100 µM) slowed down the locomotor rhythm by increasing the locomotor burst duration and decreasing the burst frequency without apparent effect on burst amplitude (Fig 5B-D). We also observed a decrease in the cross-correlogram coefficient in presence of Ba^2+^ related to a decreased rhythmic alternation (Fig 5E-F) confirming the role of Kir4.1 channels in modulating rhythmogenesis at the network level.

**Figure 5.**
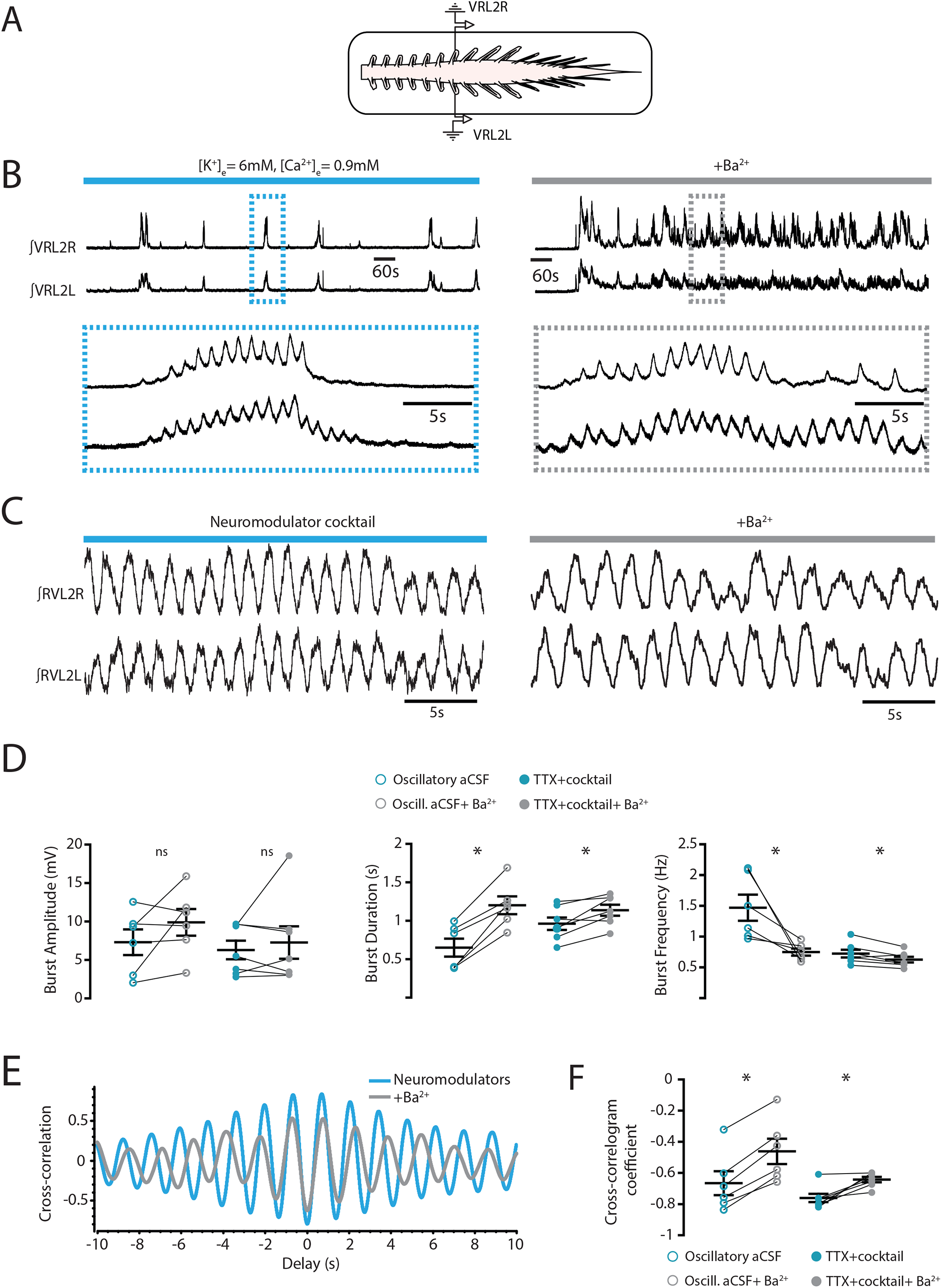
Modulation of astrocytic Kir4.1 channels alters the locomotor-like activity pattern. **A**. Schematic representation of the whole-mount spinal cord (ventral side up) with two contralateral recording glass electrodes. VRL2L stands for ventral root lumbar segment 2 left. VRL2R stands for ventral root lumbar segment 2 right. **B-C**. Extracellular recordings of alternating rhythmic activities of L2 segment ventral roots (integrated signals) in response to [K^+^]_e_ rise and [Ca^2+^]_e_ decrease (B, left) or in response to the neuromodulator cocktail (C, left) and in the presence (B-C, right) of Ba^2+^. **D**. Quantification of the effect of Ba^2+^ (grey circles) on the amplitude (left), duration (middle), and frequency (right) of the rhythmic bursts recorded at the ventral roots (L2 segments) in ionic variation condition (light blue empty circles) or in presence of the neuromodulator cocktail (light blue filled circles). Each circle represents one individual animal (n = 6 mice in high [K^+^]_e_ and low [Ca^2+^]_e_ condition and n= 7 mice in neuromodulator cocktail condition). **E**. Graphical representation of the cross-correlation coefficient between cocktail-induced rhythmic activities of contralateral ventral roots of the L2 segment before (blue trace) or after (red trace) Ba^2+^ application. **F**. Quantification of the mean cross-correlation coefficients between the rhythmic activities of the contralateral ventral roots of the L2 segment before (blue) and after (grey) Ba^2+^ application (n = 6 mice in high [K^+^]_e_ and low [Ca^2+^]_e_ condition and n= 7 mice in neuromodulator cocktail condition). ns, no significance, *P < 0.05 (two-tailed Wilcoxon paired test for D and F). Mean ± SEM. For detailed P values, see Source data file.

### Loss-of-function of Kir4.1 decreases the probability for ventromedial CPG interneurons to oscillate

To further explore the functional role of Kir4.1 channels in spinal rhythmicity, we performed their targeted loss-of function by using a short hairpin RNAs (shRNAs). We injected intrathecally at birth at T13–L1 level an AAV9 virus encoding the Kir4.1-shRNA with a eGFP reporter (Fig. 6A-B). The viral transfection led to a decrease of ∼40% in Kir4.1 membrane protein expression in the lumbar spinal cord (Fig. 6C). Concomitantly, a strong expression of eGFP in the ventromedial part of the spinal cord was observed from T1 to L5-S1 (Fig. 6D, Fig. S5). We first examined the effect of Kir4.1-shRNA on glial and neuronal electrophysiological properties. In the virally transfected GFP+ astrocytes (Fig. S6A-B), we observed in whole-cell configuration a more depolarized RMP (Fig. S6C), a higher input resistance (Fig. S6C), a marked change in the current-voltage (I-V) relationship and a reduced Ba^2+^-sensitive current in mice transduced with Kir4.1-ShRNA compared to Ctrl-ShRNA mice (Fig. S6D-I). We also observed that the RMPs of ventromedial interneurons surrounded by the virally transfected GFP+ astrocytes (Fig. 6E) from mice transduced with Kir4.1-ShRNA were overall more depolarized than the RMPs of interneurons from mice infected with Ctrl-ShRNA (Fig. 6E-F), suggesting that knockdown of Kir4.1 in astrocytes had a widespread impact on the excitability of neighbouring neurons.

**Figure 6.**
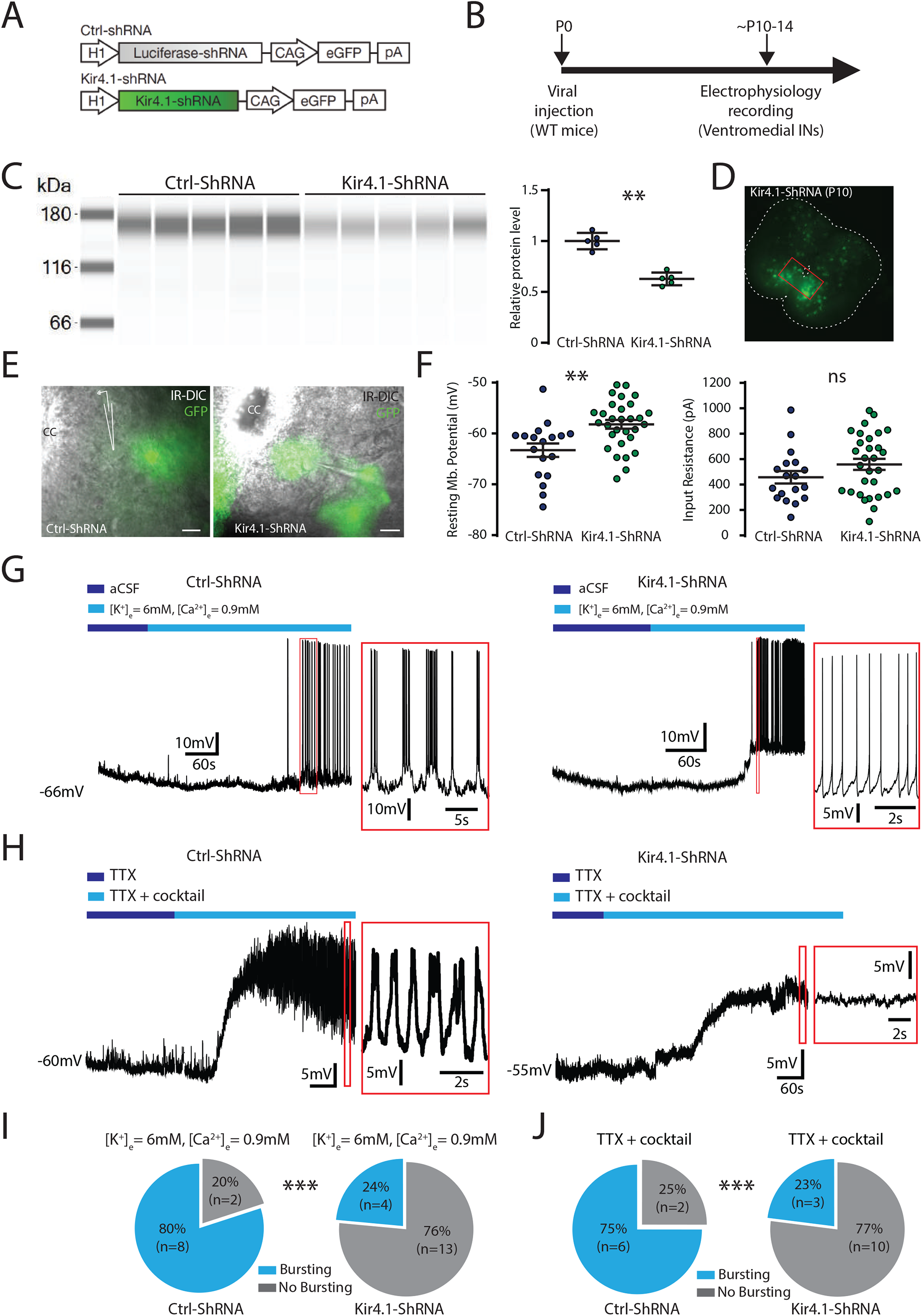
Targeted decrease in Kir4.1 channels reduces the occurrence of neuronal oscillations. **A**. Schematics of the AAV vector engineered to overexpress shRNA or dominant-negative Kir4.1. H1, human H1 promoter; CAG, CMV early enhancer/chicken *Actb* promoter. **B**. Schematic representation of the experimental design. WT for wild-type, INs for interneurons and P for postnatal day. **C**. Left: Kir4.1 pseudo-gel images from Capillary Western Blot of lumbar segments from P14 intrathecally injected at birth with an adeno-associated virus (AAV9) encoding either a scramble shRNA (n = 5 mice) or a Kir4.1-targeting shRNA (n = 5 mice). One mouse per lane. Right: group mean quantification of the ∼168 kDa band normalized to scramble-injected controls. **D**. Representative image of GFP in lumbar slices from P10 mice after intrathecal injection at birth of an adeno-associated virus (AAV9) encoding a Kir4.1-targeting shRNA (Kir4.1-ShRNA). **E**. DIC image of the ventromedial part of a lumbar slice and the corresponding one-photon GFP signal (right) of a P10 mouse injected with AAV9-Ctrl-ShRNA (left) or AAV9-Kir4.1-ShRNA (right). CC means central canal. Scale bars, 20µm. **F**. Quantification of the mean RMP (left), input resistance (middle) and cocktail-evoked membrane depolarisation (right) of ventromedial interneurons from wild-type mice injected with AAV9 encoding for Ctrl-ShRNA (left, n = 18 neurons from 4 mice) or Kir4.1-ShRNA (right, n = 30 neurons from 5 mice). **G-H**. Representative voltage traces of one interneuron of a mouse injected with AAV9-Ctrl-ShRNA (left) or AAV9-Kir4.1-ShRNA (right) in response to ionic changes (E) or in response to the neuromodulator cocktail in presence of TTX (F). **I-J**. Proportion of oscillatory interneurons (blue) or not (grey) from mice injected at birth with an AAV9 encoding either a Luciferase shRNA (Ctrl-ShRNA, n = 4 mice) (left) or a Kir4.1-targeting shRNA (Kir4.1-ShRNA, n = 7 mice) (right) in presence of high [K^+^]_e_ and low [Ca^2+^]_e_ (H) or in presence of the neuromodulator cocktail + TTX (I). ns, no significance, **P < 0.01, ***P < 0.001 (two-tailed Mann-Whitney test for C and F, two-tailed Fisher test for I and J). Mean ± SEM. For detailed P values, see Source data file.

We then tested the ability of the ventromedial interneurons to oscillate after the viral transfection of the spinal cord. We found a significant lower proportion of rhythmic bursting in ventromedial interneurons surrounded by GFP+ astrocytes in mice transduced with Kir4.1-ShRNA compared to Ctrl-ShRNA in response to both ionic variations (∼24% vs 80%) (Fig. 6G and 6I) or bath perfusion of the neuromodulator cocktail in presence of TTX (∼ 23% vs 75 %) (Fig. 6H and 6J). Altogether, these data demonstrate the pivotal role of astrocytic Kir4.1 channels in the oscillatory pattern of ventromedial interneurons from the CPG.

### Loss-of-function of Kir4.1 slows down the *in vivo* locomotor pattern and decreases the locomotor performances

We then tested whether the knockdown of Kir4.1 in the spinal cord has a functional impact on the locomotor behaviors. We thus injected intrathecally at birth an AAV9 encoding either the Kir4.1-shRNA or the Ctrl-shRNA. From 12 to 18 days after the viral injection, we performed a set of behavioural locomotor tests on freely moving mice (Fig. 7A). We first used footprint analysis to evaluate the locomotor pattern (Fig. 7B). Mice transduced with Kir4.1-shRNA displayed an increase in both swing and stand phases compared to Ctrl-ShRNA mice (Fig. 7C-D) related to a slower speed of locomotion and a decrease in regularity index without any modification of the base of support (Fig. 7D). We next tested whether Kir4.1 channels also modulate locomotor behaviours in more challenging tasks. We thus assayed motor coordination using the rotarod and swimming forced test without practice sessions to avoid any compensatory learning mechanisms. Consistently, mice transduced with Kir4.1-shRNA failed to adapt to accelerated speed (Fig. 7E) and displayed a lower swimming performance (Fig. 7F). Importantly, we did not observe any modification of the motoneuronal soma size in mice transduced with Kir4.1-ShRNA compared to Ctrl-ShRNA mice (Fig. S7). In sum, the Kir4.1-mediated inwardly rectifying K^+^ current appears to play a key role for locomotor coordination and the maintenance of the locomotor pattern in freely moving mice.

**Figure 7.**
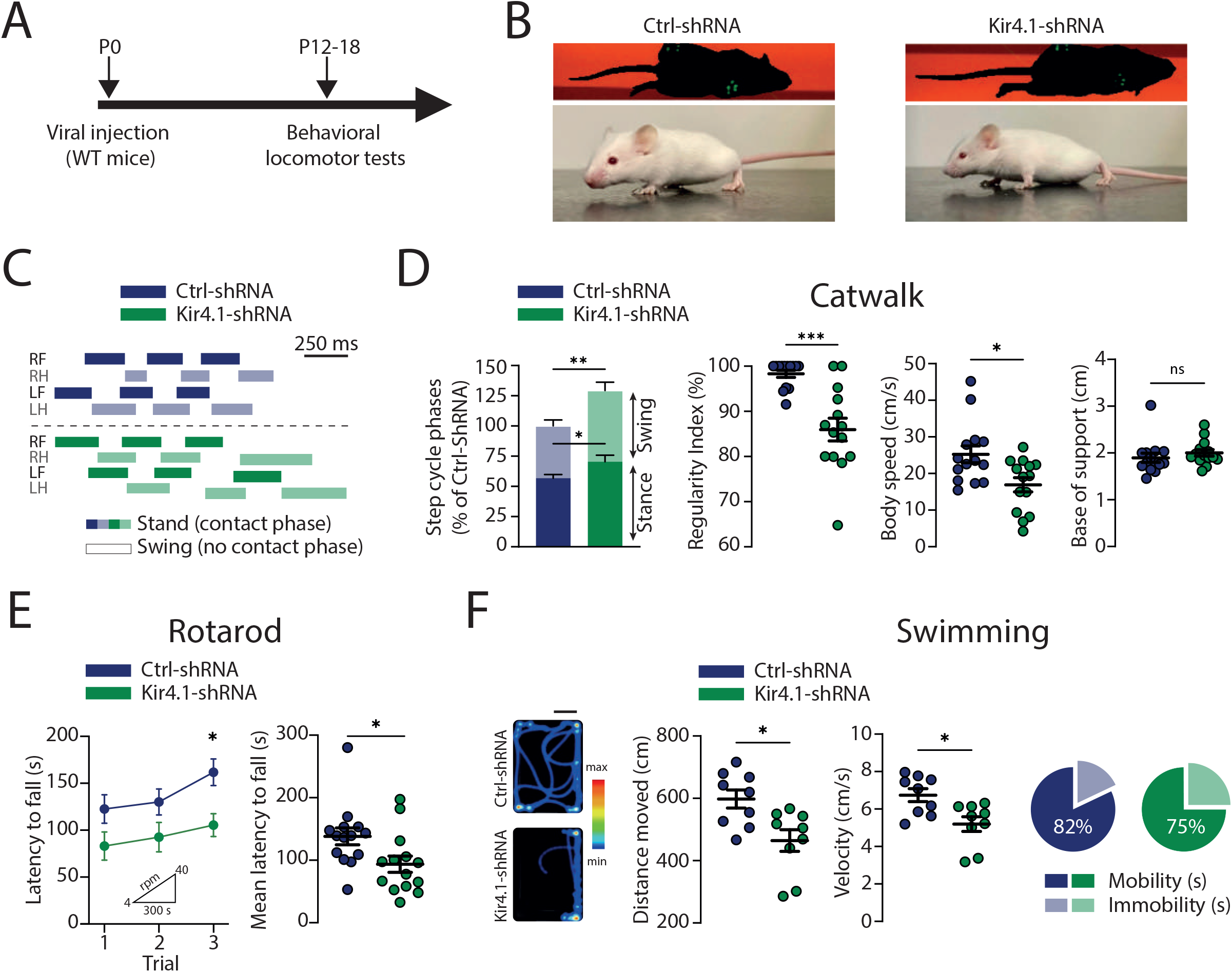
Astrocytic Kir4.1 channels influence the locomotor behaviour. **A**. Schematic representation of the experimental design. WT for wild-type and P for postnatal day. **B**. *Top*. Bottom view of P18 mice injected at birth with AAV9 encoding either for the Ctrl-ShRNA (left) or the Kir4.1-ShRNA (right) freely walking on a glass platform (Catwalk). *Bottom*. Side view of the same P18 mice. **C**. Representative footfall diagrams during CatWalk locomotion of a Ctrl-ShRNA P18 mouse (blue, top) or Kir4.1-ShRNA P18 mice (green, bottom). The stance phase is indicated by horizontal bars and the swing phase by open spaces. RF for Right Forelimb, RH for Right Hindlimb, LF for Left Forelimb, LH for Left Hindlimb. **D**. Comparative quantification of the normalized step cycle phases (stance, swing) (left), regularity index (middle left), body speed (middle right) and hind limb base of support (right) between Ctrl-ShRNA mice (blue, n = 14 mice) and Kir4.1-ShRNA mice (green, n = 14 mice). Each circle represents an individual mouse. **E**. Latency to fall from a rod rotating at accelerated speed (4–40 rpm) during 300s of wild-type mice transduced either with the Ctrl-ShRNA (blue, n = 14 mice) or with Kir4.1-ShRNA (green, n = 14 mice). **F.** *Left:* Heatmaps illustrate the swimming activity of two wild-type mice transduced either with the Ctrl-ShRNA (top) or with the Kir4.1-shRNA (bottom). Scale bar, 10 cm. *Middle:* Comparison of the mean swimming distance (middle left) and the mean velocity (middle right) between Ctrl-shRNA (blue, n = 9 mice) and Kir4.1-shRNA (green, n = 9 mice) mice. Each circle represents one individual mouse. *Right:* Percentage of mobility and immobility for Ctrl-shRNA mice (blue, n = 9 mice) and Kir4.1-shRNA mice (green, n = 9 mice) during the swimming forced test. ns, no significance, *P < 0.05, **P < 0.01, ***P < 0.001 (Mann–Whitney test for D, E right and F, Two-way ANOVA with Sidak’s multiple comparisons test for E left). Mean ± SEM. For detailed P values, see Source data file.

## DISCUSSION

This study demonstrates that astrocytic regulation of extracellular K^+^ plays a critical role in maintenance of neuronal rhythmogenesis in the spinal cord motor circuitry. We report that spinal astrocytes from neonatal mice display Ca^2+^ signals before and during rhythmic neuronal oscillations. Inhibiting astrocytic Ca^2+^ transients with BAPTA decreases the Ba^2+^-sensitive inwardly rectifying K^+^ current underlying K^+^ uptake. The Kir4.1 channels mediating the K^+^ current are expressed in astrocytes enwrapping the ventromedial interneurons from the locomotor CPG. We also demonstrated that knockdown of Kir4.1 decreases the ability of the ventromedial interneurons to oscillate. This perturbation results in an alteration of the locomotor pattern and in lower locomotor performances in challenging tasks.

### Functional coupling of astrocytic Ca^2+^ signals and neuronal oscillations

Recent studies have highlighted unexpected contribution of astrocytic Ca^2+^ signals in regulating rhythmic behaviours including respiration (Sheikhbahaei et al., 2018), mastication (Morquette et al. 2015) and locomotion (Broadhead and Miles, 2020). However, it is still incompletely understood whether the increased astrocytic Ca^2+^ transient is a prerequisite or a consequence of neuronal oscillations. Here, we reveal that ventromedial astrocytes depolarize and display an increase in Ca^2+^ signals both before and in response to ions and neuromodulator changes related to neuronal oscillations. Our two-photon Ca^2+^ imaging data reveal a functional coupling between spinal cord astrocytes and CPG interneurons. The simultaneous imaging of neuronal and astrocytic activity led to the characterization of distinct subsets of active astrocytes in the spinal CPG area. In one subset, the response precedes neuronal oscillation. This active role of astrocytes in rhythmicity has been suggested in the trigeminal sensorimotor circuit (Morquette et al., 2015) or in cortical areas where the increase in astrocytic Ca^2+^ events precedes the switch to the slow-oscillation state (Poskanzer and Yuste, 2016). In another group, the responses are concomitant to neuronal bursting. In line with this, in Drosophila astrocyte Ca^2+^ transients are driven by rhythmic firing of the octopaminergic neurons (Ma et al., 2016). A recent study also demonstrated such a dynamics in fish radial astroglia (analogs of the mammalian astrocyte), where the noradrenergic neuronal activity precedes the Ca^2+^ wave observed in radial astroglia (Mu et al., 2019). However, we cannot totally rule out that one astrocyte may belong to both groups. Altogether, our data highlight a bi-directional coupling between astrocytic Ca^2+^ signal and neuronal oscillations.

### The Ca^2+^-dependent K^+^ uptake in spinal astrocytes

Astrocytic Ca^2+^ transients are often associated to gliotransmission (Araque et al. 2014). Here we propose that an alternative mechanism to the classical gliotransmission operates for spinal rhythmogenesis. Previous works demonstrated that GPCR-mediated Ca^2+^ signalling in astrocytes is linked to an increase of K^+^ uptake in hippocampus (Wang et al., 2012a) and cerebellum (Wang et al., 2012b). This astrocytic Ca^2+^ transients stimulate the Na^+^, K^+^ ATPase pump resulting in a transient decrease in [K^+^]_e_. In our study, we provide for the first time evidence that a blockade of the Ca^2+^ transients from ventromedial lumbar astrocytes significantly reduces the Ba -sensitive current mediated by Kir4.1. This result is in line with observations showing that menthol-induced Ca^2+^ transients in astrocytes increases the Ba^2+^ -sensitive Kir4.1 current from a U251 cell line (Ratto et al., 2020). Because (i) the locomotor-like activity is associated with [K^+^]_e_ homeostasis (Brocard et al., 2013), and (ii) the astrocytic K^+^ uptake is activated by intracellular Ca^2+^ transients (Wang et al., 2012a), we assume that the Ca^2+^ dependent astrocytic K ^+^homeostasis plays a key role in the spinal rhythmicity.

### Astrocytic-dependent K^+^ regulation of the spinal rhythmogenesis

The maintenance of K^+^ homeostasis by astrocytes is crucial in modulating neuronal excitability (Dallérac et al., 2013; Neprasova et al., 2007). However, whether spinal astrocytes may exert a “ionostatic control” of neuronal bursting, as suggested by Ding et al. 2016, is still unclear. Here, we evaluated the consequence of blocking the astrocytic inwardly rectifying K^+^ (Kir4.1) channels on neuronal oscillations in the locomotor CPG. Blocking the Kir4.1-mediated astrocytic K^+^ uptake with Ba^2+^ (100 µM) (Cui et al., 2018; Djukic et al., 2007; Ransom and Sontheimer, 1995) decreases the frequency of neuronal oscillations until progressively preventing them. In line with this effect, injection of Kir4.1-ShRNA resulting in the decrease in Kir4.1 astrocytic membrane expression also significantly reduces the probability of driving neuronal oscillations. These results highlight the key role of astrocytic K^+^ uptake in maintaining the neuronal oscillatory pattern. Although the astrocytic Kir4.1 channels are not homogeneously expressed among and within brain structures (Higashi et al. 2001), we showed a strong Kir4.1 expression in astrocytes enwrapping the ventromedial bursting interneurons and lamina IX motoneurons. This strategic distribution of Kir4.1 channels surrounding the bursting spinal neurons may be viewed as a powerful mechanism to rapidly modulate network rhythmicity. Moreover, since glutamate tunes the CPG network by exerting a speed control of locomotor-like activity (Talpalar et al., 2013) and that [K^+^]e provides the driving force for glutamate uptake (Nwaobi et al., 2016; Verkhratsky and Nedergaard, 2018) the effective K^+^ uptake from the extracellular space through Kir4.1 is thus crucial for tuning the neuron oscillations. In line with this, previous works highlighted the importance of [K^+^]_e_ for rhythmogenesis in hippocampal (Jensen et al. 1994) or cortical (Bellot-Saez et al., 2018) networks. Furthermore, the increase in [K^+^]_e_ induced by neuromodulators may promote the shift between neuronal oscillatory states in cortical slices electrically silenced by TTX (Ding et al., 2016). A computational work also demonstrated that extracellular K^+^ dynamics can cause transition between fast tonic spiking and slow bursting in neocortical networks (Frohlich et al., 2006) in the same range than the spinal locomotor network.

### The ventromedial astrocytes regulate the locomotor pattern in vivo

A dense literature pointed out that during locomotion, mice display enhanced Ca^2+^ signals in astrocytes of the cerebellum (Nimmerjahn et al., 2009; Paukert et al., 2014), visual cortex (Paukert et al., 2014), or somatosensory cortex (Bojarskaite et al., 2020; Dombeck et al., 2007). On the other hand, silencing astrocyte Ca^2+^ activity in brainstem with optogenetics or chemogenetics can modulate the rhythmic respiratory behaviour (Gourine et al., 2010; Sheikhbahaei et al., 2018). In this study, we used genetic tools with ShRNA to knockdowns Kir4.1 in the locomotor CPG and demonstrate that the astrocytic regulation of K^+^ homeostasis plays a key role in controlling the locomotor speed and challenging locomotor tasks *in vivo*. Mice transduced with Kir4.1-ShRNA showed striking locomotor deficits compared to Ctrl-ShRNA mice. Kir4.1 had been reported to contribute to the maintenance of the soma size of large α-motoneurons and peak strength in adult mice (Kelley et al., 2018). We demonstrated that the lowered locomotor performances from Kir4.1-ShRNA mice do not result from changes in motoneuronal morphological features. Indeed, we did not observe any modification of the motoneuronal soma size in mice transduced with Kir4.1-ShRNA compared to Ctrl-ShRNA mice. Tong et al. 2014 observed that the Kir4.1 expression decrease in the striatal network of a mouse model of Huntington disease, results in altered locomotor pattern (Tong et al., 2014). We did not observe any Kir4.1-ShRNA expression in brain or brainstem structures involved in the locomotor control. This result is in line with the restricted diffusion to the thoraco-lumbar segments of the AAV9-ShRNA injected intrathecally at birth in the lumbar segments (Bos et al., 2021). Thus we assume that the behavioural consequence of Kir4.1 knockdown on locomotor performances is due to Kir4.1 decrease in the spinal locomotor networks.

Overall this study defines the contours of a new concept for the spinal locomotor CPG. We propose that the astrocytic Ca^2+^ transients which occurs during the neuronal oscillations likely maintains extracellular K^+^ homeostasis, which in turn adjusts the locomotor pattern (Graphical abstract). Further studies will be needed to decipher the importance for spinal rhythmicity of the astrocytic syncytium mediating K^+^ buffering in the locomotor CPG.

## EXPERIMENTAL PROCEDURES

Further details and an outline of resources used in this work can be found in the Supplemental Procedures.

### Mice

Mice from the 1^st^ to 3^rd^ postnatal week of either sex were used in this study. Animals from different litters were used for each experiment. All animal care and use were conformed to the French regulations (Décret 2010-118) and approved by the local ethics committee (Comité d’Ethique en Neurosciences INT-Marseille, CE71 NbA1301404, authorization Nb 2018110819197361). See the Supplemental Experimental Procedures for more details.

### *Ex vivo* Models

Slice preparations were used for whole-cell recordings and two-photon Ca^2+^ imaging experiments, whereas whole spinal cord preparations were used for fictive locomotion experiments. Preparation procedures are detailed in the Supplemental Experimental Procedures.

### Intracellular Recordings

For *ex vivo* experiments, whole-cell patch-clamp recordings were made from L1-L2 ventromedial GFP+ interneurons (*Hb9:eGFP* mice) or GFP+ astrocytes (*Aldh1L1:eGFP* mice) from lumbar slices. Procedures of intracellular recordings are detailed in the Supplemental Experimental Procedures.

### Extracellular Recordings

Motor outputs were recorded using glass suction electrodes placed in contact with right and left lumbar L2 ventral roots in response to bath application of a monoaminergic cocktail. See the Supplemental Experimental Procedures for more details.

### Two-photon Ca^2+^ imaging

Two-photon excitation of Gcamp6f and jRGECO1a was simultaneously evoked with a laser tuned to 960 nm. Imaging was done at 30.5202 Hz. The microscope was equipped with two detection channels for fluorescence imaging. See the Supplemental Experimental Procedures for more details.

### shRNA constructs

Specific shRNA sequence designed to knockdown *Kir4*.*1* transcript was incorporated into an adeno-associated viral (AAV) vector (serotype 9), which features a H1 promoter to drive shRNA expression and a CAG promoter to drive eGFP expression for identification of transduced cells (Cui et al. 2018). See the Supplemental Experimental Procedures for more details.

### Intrathecal vector delivery

A minimally-invasive technique was used to micro-inject adeno-associated viral (AAV) vectors into the T13-L1 intervertebral space. A total volume of 2 µL /animal was injected. See the Supplemental Experimental Procedures for more details.

### Kir4.1 protein quantification

Kir4.1 expression was analyzed using the 12-230 kDa separation module (SM-W004 ProteinSimple) on an automated capillary western blotting system (‘Jess’ ProteinSimple). See the Supplemental Experimental Procedures for more details.

### Immunohistochemistry

Transverse spinal cord sections at the lumbar L1-L2 level were processed for immunohistochemistry using antibodies against Kir4.1, NeuN. Tissue processing and staining are detailed in the Supplemental Experimental Procedures.

### Behavioral tests

Assessment of motor behaviour were performed on neonatal mice with three different tests: walking, rotarod and swimming. See the Supplemental Experimental Procedures for more details.

### Statistical Analysis

Group measurements were expressed as means ± S.E.M. All statistical analyses are indicated in Figure legends. The level of significance was set at p < 0.05. Statistical analyses were performed using Graphpad Prism 9 software. See the Supplemental Experimental Procedures for more details.

## ACKNOWLEDGMENTS

This work was mainly funded by the CNRS and “Fonds d’Investissement de l’INT jeunes chercheuses, jeunes chercheurs” (FI_INT_JCJC_2019) (to R.B.) and a small part by the Agence National de la Recherche Scientifique (CalpaSCI, ANR-16-CE16-0004) (to F.B.). We thank Ivo Vanzetta, Pascal Weber and Sebastien Roux for their technical advices in imaging sessions, Anne Duhoux for animal care, Cécile Brocard for genotyping and Jérémy Verneuil for providing the Matlab script. We also gratefully acknowledge Eduardo Gascon, Florence Jaouen, Catherine Lepolard and Ana Borges-Correia from Neuro-Vir platform of the Neuro-Bio-Tools facility (Institut de Neurosciences de la Timone, UMR 7289, Marseille, France) for their advices, support and assistance in the design and production of vectors used in this work. We also thank Geneviève Rougon for her critical insights on the manuscript.

## AUTHOR CONTRIBUTIONS

Research design, T.B., E.P. and R.B..; Methodology, T.B., E.P., N.R. and R.B.; Investigation and data analysis, T.B., E.P., M.D., and R.B.; Writing, R.B.; Funding Acquisition, R.B. and F.B..; Supervision, R.B.

---

**Graphical Abstract.**
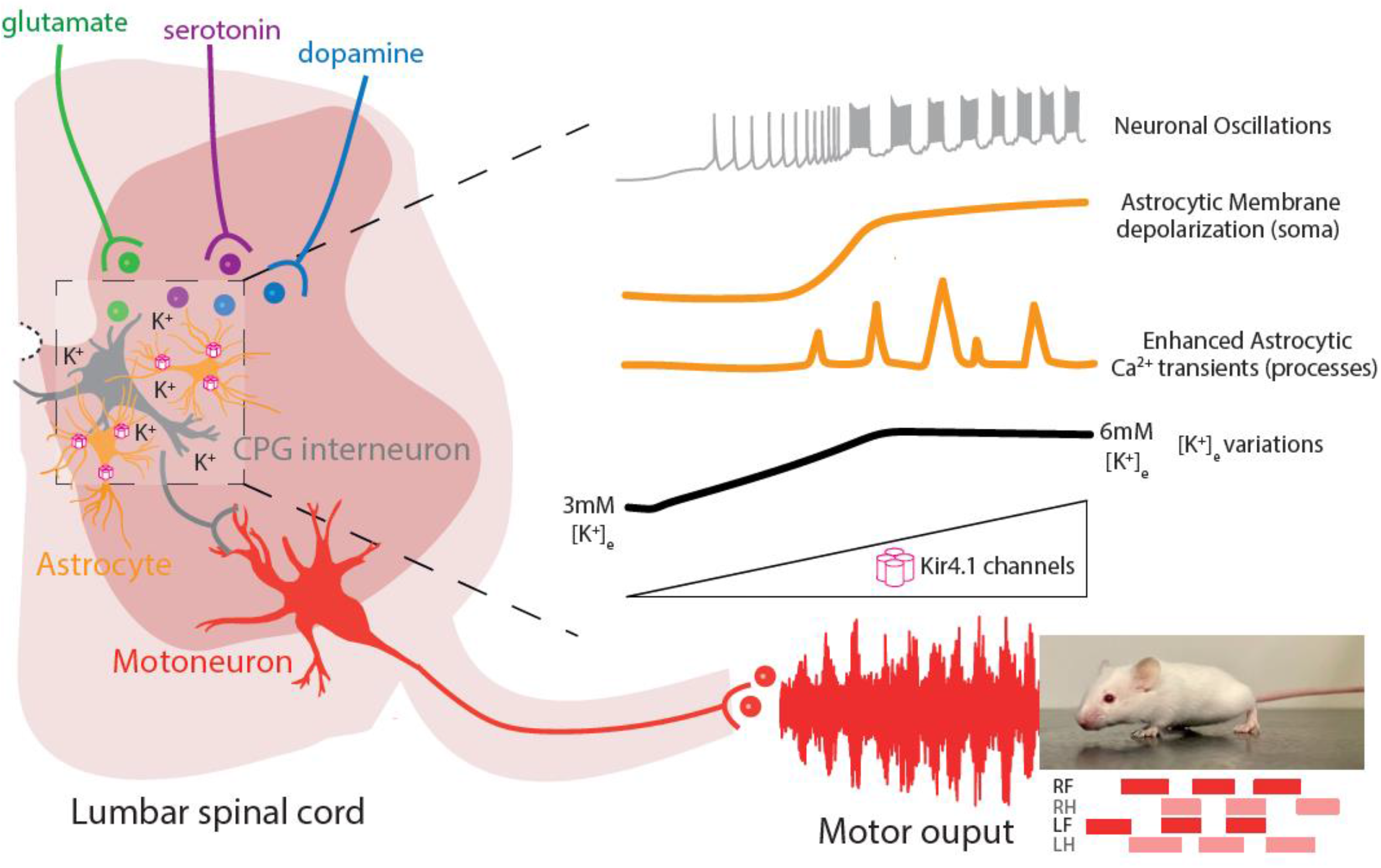
Overview of the astrocytic K^+^ uptake mechanisms influencing the locomotor pattern. Schematic relationship between astrocytes and neurons in the spinal locomotor central pattern generator (CPG) network. K^+^ for potassium, RF for right front limb, LF for left front limb, RH for right hind limb, LH for left hind limb.

## REFERENCES

Araque, A., Carmignoto, G., Haydon Philip G., Oliet Stéphane H.R., Robitaille, R., and Volterra, A. (2014). Gliotransmitters Travel in Time and Space. Neuron 81, 728–739.

Bellot-Saez, A., Cohen, G., van Schaik, A., Ooi, L., J, W.M., and Buskila, Y. (2018). Astrocytic modulation of cortical oscillations. Sci Rep 8, 11565.

Bellot-Saez, A., Kekesi, O., Morley, J.W., and Buskila, Y. (2017). Astrocytic modulation of neuronal excitability through K(+) spatial buffering. Neurosci Biobehav Rev 77, 87–97.

Ben Haim, L., Carrillo-de Sauvage, M.A., Ceyzeriat, K., and Escartin, C. (2015). Elusive roles for reactive astrocytes in neurodegenerative diseases. Front Cell Neurosci 9, 278.

Bojarskaite, L., Bjornstad, D.M., Pettersen, K.H., Cunen, C., Hermansen, G.H., Abjorsbraten, K.S., Chambers, A.R., Sprengel, R., Vervaeke, K., Tang, W., et al. (2020). Astrocytic Ca(2+) signaling is reduced during sleep and is involved in the regulation of slow wave sleep. Nat Commun 11, 3240.

Bos, R., Drouillas, B., Bouhadfane, M., Pecchi, E., Trouplin, V., Korogod, S.M., and Brocard, F. (2021). Trpm5 channels encode bistability of spinal motoneurons and ensure motor control of hindlimbs in mice. Nat Commun 12, 6815.

Bos, R., Harris-Warrick, R.M., Brocard, C., Demianenko, L.E., Manuel, M., Zytnicki, D., Korogod, S.M., and Brocard, F. (2018). Kv1.2 Channels Promote Nonlinear Spiking Motoneurons for Powering Up Locomotion. Cell Reports 22, 3315–3327.

Bracci, E.B. M.; Nistri, A. (1998). Extracellular K+ Induces Locomotor-Like Patterns in the Rat Spinal Cord In Vitro: Comparison With NMDA or 5-HT Induced Activity. J Neurophysiol 79, 2643–2652.

Broadhead, M.J., and Miles, G.B. (2020). Bi-Directional Communication Between Neurons and Astrocytes Modulates Spinal Motor Circuits. Front Cell Neurosci 14, 30.

Brocard, F., Shevtsova Natalia A., Bouhadfane, M., Tazerart, S., Heinemann, U.,, Rybak Ilya A., and Vinay, L. (2013). Activity-Dependent Changes in Extracellular Ca2+ and K+ Reveal Pacemakers in the Spinal Locomotor-Related Network. Neuron 77, 1047–1054.

Brocard, F., Tazerart, S., and Vinay, L. (2010). Do Pacemakers Drive the Central Pattern Generator for Locomotion in Mammals? The Neuroscientist 16, 139–155.

Cahoy, J.D., Emery, B., Kaushal, A., Foo, L.C., Zamanian, J.L., Christopherson, K.S., Xing, Y., Lubischer, J.L., Krieg, P.A., Krupenko, S.A., et al. (2008). A Transcriptome Database for Astrocytes, Neurons, and Oligodendrocytes: A New Resource for Understanding Brain Development and Function. Journal of Neuroscience 28, 264–278.

Carlsen, E.M., and Perrier, J.-F.o. (2014). Purines released from astrocytes inhibit excitatory synaptic transmission in the ventral horn of the spinal cord. Frontiers in Neural Circuits 8.

Cui, Y., Yang, Y., Ni, Z., Dong, Y., Cai, G., Foncelle, A., Ma, S., Sang, K., Tang, S., Li, Y., et al. (2018). Astroglial Kir4.1 in the lateral habenula drives neuronal bursts in depression. Nature 554, 323–327.

Dallérac, G., Chever, O., and Rouach, N. (2013). How do astrocytes shape synaptic transmission? Insights from electrophysiology. Frontiers in Cellular Neuroscience 7.

Ding, F., O’Donnell, J., Xu, Q., Kang, N., Goldman, N., and Nedergaard, M. (2016). Changes in the composition of brain interstitial ions control the sleep-wake cycle. Science 352, 550–555.

Djukic, B., Casper, K.B., Philpot, B.D., Chin, L.-S., and McCarthy, K.D. (2007). Conditional Knock-Out of Kir4.1 Leads to Glial Membrane Depolarization, Inhibition of Potassium and Glutamate Uptake, and Enhanced Short-Term Synaptic Potentiation. Journal of Neuroscience 27, 11354–11365.

Dombeck, D.A., Khabbaz, A.N., Collman, F., Adelman, T.L., and Tank, D.W. (2007). Imaging Large-Scale Neural Activity with Cellular Resolution in Awake, Mobile Mice. Neuron 56, 43–57.

El Manira, A. (2014). Dynamics and plasticity of spinal locomotor circuits. Current Opinion in Neurobiology 29, 133–141.

Frohlich, F., Bazhenov, M., Timofeev, I., Steriade, M., and Sejnowski, T.J. (2006). Slow state transitions of sustained neural oscillations by activity-dependent modulation of intrinsic excitability. J Neurosci 26, 6153–6162.

Gourine, A.V., Kasymov, V., Marina, N., Tang, F., Figueiredo, M.F., Lane, S., Teschemacher, A.G., Spyer, K.M., Deisseroth, K., and Kasparov, S. (2010). Astrocytes Control Breathing Through pH-Dependent Release of ATP. Science 329, 571–575.

Grillner, S., and El Manira, A. (2020). Current Principles of Motor Control, with Special Reference to Vertebrate Locomotion. Physiol Rev 100, 271–320.

Higashi, K., Fujita, A., Inanobe, A., Tanemoto, M., Doi, K., Kubo, T., Kurachi, Y. (2001). An inwardly rectifying K1 channel, Kir4.1, expressed in astrocytes surrounds synapses and blood vessels in brain. Am. J. Physiol. Cell Physiol. 281, 922–931.

Jensen, M.S., Azouz, R., Yaari, Y. (1994). Variant Firing Patterns in Rat Hippocampal Pyramidal Cells Modulated by Extracellular Potassium. J. Neurophysiol. 71, 831–9.

Kelley, K.W., Ben Haim, L., Schirmer, L., Tyzack, G.E., Tolman, M., Miller, J.G., Tsai, H.-H., Chang, S.M., Molofsky, A.V., Yang, Y., et al. (2018). Kir4.1-Dependent Astrocyte-Fast Motor Neuron Interactions Are Required for Peak Strength. Neuron 98, 306-319.e307.

Kiehn, O. (2016). Decoding the organization of spinal circuits that control locomotion. Nature Reviews Neuroscience 17, 224–238.

Kofuji, P., and Newman, E.A. (2004). Potassium buffering in the central nervous system. Neuroscience 129, 1043–1054.

Lee, H.S., Ghetti, A., Pinto-Duarte, A., Wang, X., Dziewczapolski, G., Galimi, F., Huitron-Resendiz, S., Pina-Crespo, J.C., Roberts, A.J., Verma, I.M., et al. (2014). Astrocytes contribute to gamma oscillations and recognition memory. Proceedings of the National Academy of Sciences 111, E3343–E3352.

Ma, Z., Stork, T., Bergles, D.E., and Freeman, M.R. (2016). Neuromodulators signal through astrocytes to alter neural circuit activity and behaviour. Nature 539, 428–432.

Masino, M.A., Abbinanti, M.D., Eian, J., and Harris-Warrick, R.M. (2012). TTX-resistant NMDA receptor-mediated membrane potential oscillations in neonatal mouse Hb9 interneurons. PLoS One 7, e47940.

Montalant, A., Carlsen, E.M.M., and Perrier, J.F. (2021). Role of astrocytes in rhythmic motor activity. Physiol Rep 9, e15029.

Morquette, P., Verdier, D., Kadala, A., Féthière, J., Philippe, A.G., Robitaille, R., and Kolta, A. (2015). An astrocyte-dependent mechanism for neuronal rhythmogenesis. Nature Neuroscience 18, 844–854.

Mu, Y., Bennett, D.V., Rubinov, M., Narayan, S., Yang, C.T., Tanimoto, M., Mensh, B.D., Looger, L.L., and Ahrens, M.B. (2019). Glia Accumulate Evidence that Actions Are Futile and Suppress Unsuccessful Behavior. Cell 178, 27–43 e19.

Neprasova, H., Anderova, M., Petrik, D., Vargova, L., Kubinova, S., Chvatal, A., and Sykova, E. (2007). High extracellular K(+) evokes changes in voltage-dependent K(+) and Na (+) currents and volume regulation in astrocytes. Pflugers Arch 453, 839–849.

Nimmerjahn, A., Mukamel, E.A., and Schnitzer, M.J. (2009). Motor Behavior Activates Bergmann Glial Networks. Neuron 62, 400–412.

Nwaobi, S.E., Cuddapah, V.A., Patterson, K.C., Randolph, A.C., and Olsen, M.L. (2016). The role of glial-specific Kir4.1 in normal and pathological states of the CNS. Acta Neuropathol 132, 1–21.

Olsen, M.L., Higashimori, H., Campbell, S.L., Hablitz, J.J., and Sontheimer, H. (2006). Functional expression of Kir4.1 channels in spinal cord astrocytes. Glia 53, 516–528.

Panatier, A., Vallée, J., Haber, M., Murai Keith K., Lacaille, J.-C., and Robitaille, R. (2011). Astrocytes Are Endogenous Regulators of Basal Transmission at Central Synapses. Cell 146, 785–798.

Paukert, M., Agarwal, A., Cha, J., Doze, Van A., Kang Jin U., and Bergles Dwight E. (2014). Norepinephrine Controls Astroglial Responsiveness to Local Circuit Activity. Neuron 82, 1263–1270.

Poskanzer, K.E., and Yuste, R. (2016). Astrocytes regulate cortical state switching in vivo. Proceedings of the National Academy of Sciences 113, E2675–E2684.

Ransom, C.B., and Sontheimer, H. (1995). Biophysical and pharmacological characterization of inwardly rectifying K+ currents in rat spinal cord astrocytes. Journal of Neurophysiology 73, 333–346.

Ratto, D., Ferrari, B., Roda, E., Brandalise, F., Siciliani, S., De Luca, F., Priori, E.C., Di Iorio, C., Cobelli, F., Veneroni, P., et al. (2020). Squaring the Circle: A New Study of Inward and Outward-Rectifying Potassium Currents in U251 GBM Cells. Cell Mol Neurobiol 40, 813–828.

Rosa, J.M., Bos, R., Sack, G.S., Fortuny, C., Agarwal, A., Bergles, D.E., Flannery, J.G., and Feller, M.B. (2015). Neuron-glia signaling in developing retina mediated by neurotransmitter spillover. eLife 4.

Savtchouk, I., and Volterra, A. (2018). Gliotransmission: Beyond Black-and-White. The Journal of Neuroscience 38, 14–25.

Sheikhbahaei, S., Turovsky, E.A., Hosford, P.S., Hadjihambi, A., Theparambil, S.M., Liu, B., Marina, N., Teschemacher, A.G., Kasparov, S., Smith, J.C., et al. (2018). Astrocytes modulate brainstem respiratory rhythm-generating circuits and determine exercise capacity. Nature Communications 9.

Sibille, J., Dao Duc, K., Holcman, D., and Rouach, N. (2015). The neuroglial potassium cycle during neurotransmission: role of Kir4.1 channels. PLoS Comput Biol 11, e1004137.

Talpalar, A.E., Bouvier, J., Borgius, L., Fortin, G., Pierani, A., and Kiehn, O. (2013). Dual-mode operation of neuronal networks involved in left–right alternation. Nature 500, 85–88.

Tazerart, S., Vinay, L., and Brocard, F. (2008). The Persistent Sodium Current Generates Pacemaker Activities in the Central Pattern Generator for Locomotion and Regulates the Locomotor Rhythm. Journal of Neuroscience 28, 8577–8589.

Tong, X., Ao, Y., Faas, G.C., Nwaobi, S.E., Xu, J., Haustein, M.D., Anderson, M.A., Mody, I., Olsen, M.L., Sofroniew, M.V., et al. (2014). Astrocyte Kir4.1 ion channel deficits contribute to neuronal dysfunction in Huntington’s disease model mice. Nature Neuroscience 17, 694–703.

Tsai, H.-H., Li, H., Fuentealba, L.C., Molofsky, A.V., Taveira-Marques, R., Zhuang, H., Tenney, A., Murnen, A.T., Fancy, S.P.J., Merkle, F., et al. (2012). Regional Astrocyte Allocation Regulates CNS Synaptogenesis and Repair. Science 337, 358–362.

Verkhratsky, A., Bush, N., Nedergaard, M., and Butt, A. (2018). The Special Case of Human Astrocytes. Neuroglia 1, 21–29.

Verkhratsky, A., and Nedergaard, M. (2018). Physiology of Astroglia. Physiol Rev 98, 239–389.

Wang, F., Smith, N.A., Xu, Q., Fujita, T., Baba, A., Matsuda, T., Takano, T., Bekar, L., and Nedergaard, M. (2012a). Astrocytes Modulate Neural Network Activity by Ca2+-Dependent Uptake of Extracellular K+. Science Signaling 5, ra26–ra26.

Wang, F., Xu, Q., Wang, W., Takano, T., and Nedergaard, M. (2012b). Bergmann glia modulate cerebellar Purkinje cell bistability via Ca2+-dependent K+ uptake. Proc Natl Acad Sci U S A 109, 7911–7916.

Wilson, J.M. (2005). Conditional Rhythmicity of Ventral Spinal Interneurons Defined by Expression of the Hb9 Homeodomain Protein. Journal of Neuroscience 25, 5710–5719.

Witts, E.C., Nascimento, F., and Miles, G.B. (2015). Adenosine-mediated modulation of ventral horn interneurons and spinal motoneurons in neonatal mice. Journal of Neurophysiology 114, 2305–2315.

Ziskind-Conhaim, L., Wu, L., and Wiesner, E.P. (2008). Persistent sodium current contributes to induced voltage oscillations in locomotor-related hb9 interneurons in the mouse spinal cord. J Neurophysiol 100, 2254–2264.

