## Supplemental Information for "Astrocytes regulate locomotion by orchestrating neuronal rhythmicity in the spinal network via potassium clearance"

**via potassium clearance**

by

**Tony Barbay^1,3^, Emilie Pecchi^1,3^, Myriam Ducrocq^1^, Nathalie Rouach^2^, Frédéric Brocard^1^, Rémi Bos^1,4,*^**

^1^Institut de Neurosciences de la Timone (UMR7289), Aix-Marseille Université and CNRS, Marseille, France

^2^Center for Interdisciplinary Research in Biology, Collège de France, CNRS, INSERM, Labex Memolife, Université PSL, Paris, France

^3^ These authors contributed equally

^4^ Lead contact

* Correspondence to R.B.:

Number of figures: 7

Supplementary information: 7 figures

Running Title: Astrocytic modulation of spinal rhythmicity

Keywords: Locomotion, Spinal cord, Astrocytes, Neuronal Oscillations, Potassium uptake, Kir4.1.

**supplemental figure legends**

**Figure S1 (Related to Figure 1).** **Indirect measure of K^+^ rise induced by TTX cocktail. A.** Representative voltage trace of one GFP+ astrocyte from a P11 Aldh1L1-eGFP mouse in response to successive exposures of incremental extracellular K+ concentration [K^+^]_e_ (3mM, 6mM, 9mM and 12 mM). **B.** Mean of the astrocytic membrane potentials when recorded in standard aCSF with [K^+^]_e_ = 3mM (black circles), 6mM (dark blue circles), 9mM (light blue circles), 12mM (cyan circles) (n = 8 astrocytes in each condition from n = 3 mice). **C.** Quantification of the astrocytic membrane potential variations as a function of the [K^+^]_e_ rise compared to [K^+^]_e_ = 3mM. Dotted line represents the astrocytic membrane potential variation recorded in the TTX + neuromodulator cocktail condition (n = 8 astrocytes from 4 mice, ΔVm = 10.61 mV compared to 3mM). This data suggests that the neuromodulator cocktail in presence of TTX results in a [K^+^]_e_ rise of ~ 3 mM. ** P < 0.01, ***P < 0.001 (one-way ANOVA with a Dunn’s multiple comparisons test for B). Mean ± SEM. For detailed P values, see Source data file.

**Figure S2 (Related to Figure 2). Simultaneous recording of neuronal and astrocytic signals in response to TTX by using two-photon Ca^2+^ imaging in lumbar slices. A.** Schematic representation of the experimental design. Intrathecal injections of AAV9-gfaABC1D-Gcamp6f and AAV9-Syn.NES-jRGECO1a were made at birth in wild-type mice. Two-photon Ca^2+^ imaging sessions on lumbar slices (L1-L2) were processed from P13-P16 mice. Red inset represents the imaging area. **B.** Two-photon fluorescent images of Gcamp6f (astrocyte, left), jRGECO1a (neurons, middle) and merge signal (right) in the ventromedial part of a lumbar slice from a P16 mouse. CC means central canal. Scale bar is 40 µm. **C.** Examples of astrocytic Gcamp6f Ca^2+^ signals (ΔF/Fo) (left) and neuronal jRGECO1a Ca^2+^ signals (right) in presence or not of TTX. **D.** Quantification of the mean amplitude (left), duration (middle) and frequency (right) of the astrocytic and neuronal Ca^2+^ signals in absence (green circles for astrocytes red circles for neurons) or in presence (blue circles for astrocytes, pink circles for neurons) of TTX (n = 182 astrocytic ROIs from 3 mice and n = 141 neuronal ROIs from 5 mice). Ns, no significance, ***P < 0.001 (two-tailed Wilcoxon test for D). Mean ± SEM. For detailed P values, see Source data file.

**Figure S3 (Related to Figure 3). Spinal astrocytes express Kir4.1 channels enwrapping the pacemaker interneurons. A.** *Top.* Representative confocal images of the endogenous neuronal GFP signal (A1) and the immunofluorescent labelling of the Kir4.1 channels (A2) from a lumbar slice of a P12 Hb9-eGFP mouse. Note the nuclei are labelled in blue with Hoechst (A3) and the merge signal is shown in A4. White circles indicate the targeted bursting GFP+ interneurons. CC stands for central canal. Scale bar, 100µm. *Bottom.* High magnification images of the upper ventromedial part of the lumbar slice from one Hb9-eGFP mouse showing the Kir4.1 channels (A2’) enwrapping the GFP signal of a pacemaker interneuron (A1’) co-stained with Hoetsch (A3’). Merge signal is shown in A4’. Scale bar, 20µm. **B.** *Top.* Representative confocal images of the endogenous astrocytic GFP signal (B1), the immunofluorescent labelling of the Kir4.1 channels (B2) and the neurons immuno-stained with NeuN (B3) from a lumbar slice of a P8 *Aldh1L1-eGFP* mouse. Note the nuclei are labelled in blue with Hoechst (B4) and the merge signal is shown in B5. Scale bar, 50µm. CC for central canal. *Bottom.* High magnification images of the upper white inset (B5) showing the GFP+ astrocytes (B1’) expressing the Kir4.1 channels (white arrows) (B2’). The Kir4.1 signal does not merge with NeuN signal (orange arrows) (B3’). All nuclei cells are stained with Hoechst signal (B4’). Merge signal is shown in B5’. Scale bars, 20µm. **C.** High magnification (63X) images showing one ventromedial GFP+ astrocyte (C1’) expressing Kir4.1 (C2’) surrounding two ventromedial interneurons (NeuN+) which are not labelled for Kir4.1 (C3’). All nuclei cells are stained with Hoechst signal (C4’). Merge signal is shown in C5’. Scale bars, 10µm.

**Figure S4 (Related to Figure 4).** **Ba^2+^ alters the neuronal oscillations pattern before inhibiting it A.** Representative voltage trace of one GFP+ interneuron from a P7 Hb9-eGFP mouse before (blue bar) and during (grey bar) bath application of Ba^2+^ (100µM). Numerated red insets represent the high magnifications of the above voltage trace overtime. The neuron oscillates in presence of ionic variations (higher K^+^ and lower Ca^2+^, inset#1) and its bursting frequency slows down after few minutes of Ba^2+^ (inset#2) before being prevented. After more than 5 minutes of Ba^2+^, the neuron displays a tonic firing (inset #3). **B.** Representative voltage trace of one GFP+ interneuron from a P10 Hb9-eGFP mouse before (blue bars) and during (grey bar) bath application of Ba^2+^ (100µM). Numerated red insets represent the high magnifications of the above voltage trace overtime. The neuron oscillates in presence of TTX and the neuromodulator cocktail (dopamine, NMDA, 5-HT, inset#1) and its bursting frequency slows down after few minutes of Ba^2+^ (inset#2) before being prevented. After more than 5 minutes of Ba^2+^, the neuron does not display any oscillatory activity (inset #3). **C.** Quantification of the Ba^2+^ effect (grey circles) on the amplitude (left), duration (middle), and frequency (right) of the GFP+ interneurons bursts in ionic variation condition (light blue empty circles) or in presence of the neuromodulator cocktail (light blue filled circles) (n = 9 neurons from 7 mice in higher [K^+^]_e_ and lower [Ca^2+^]_e_ condition and n = 6 neurons from 3 mice in neuromodulator cocktail condition) (two-tailed Wilcoxon paired test for C). Mean ± SEM. For detailed P values, see Source data file.

**Figure S5 (Related to Figure 6). Absence of brain transduction following spinal intrathecal AAV9 administration.** Representative confocal images of Hoechst dye (left) and eGFP fluorescence signal (middle) across three different coronal sections of the brain and three coronal sections of the spinal cord from the same P15 Ctrl-ShRNA mouse. The intrathecal administration of AAV9 harboring a self-complementary genome expressing CMV.eGFP and Ctrl-ShRNA or Kir4.1-ShRNA was performed at birth on 4 different mice (n =2 Kir4.1-ShRNA mice and n = 2 Ctrl-ShRNA mice). All 4 injected mice express very weak (or not at all for 3 mice over 4) GFP fluorescence in the brain structures involved in the motor control and locomotion.

**Figure S6 (Related to Figure 6).** **Kir4.1-shRNA decreases the Ba^2+^-sensitive Kir4.1 channels current resulting in the astrocytic membrane potential depolarization and the input resistance increase. A.** Schematic representation of the experimental design. WT for wild-type, GFP for green fluorescent protein. **B.** Superimposition of the DIC image and the GFP signal of a ventromedial astrocyte from a lumbar slice of a P10 mouse injected with AAV9-Ctrl-ShRNA-GFP (left) or AAV9-Kir4.1-ShRNA-GFP (right). **C.** Mean RMP (left) and input resistance (right) of lumbar GFP+ astrocytes from 8 to 13 days-old mice injected at birth with AAV9 encoding either a control shRNA (blue circles, n = 13 astrocytes from n = 3 mice) or a Kir4.1-targeting shRNA (green circles, n = 14 astrocytes from n = 4 mice) recorded in standard aCSF. **D.** Representative membrane currents (top) of two lumbar GFP+ astrocytes of Ctrl-ShRNA (left) and Kir4.1-ShRNA (right) mice. Astrocytes were clamped at the RMP in response to incremental voltages pulses (bottom, 2.5s, -150 to -10mV, Δ = 10mV). **E.** The averaged I/V curves from lumbar GFP+ astrocytes from Ctrl-ShRNA mice (blue circles, n = 12 astrocytes from n = 3 mice) and Kir4.1-ShRNA mice (green circles, n = 14 astrocytes from n = 4 mice). **F***.* Comparative quantification of the peak amplitude Kir4.1 current at -140mV of GFP+ astrocytes from Ctrl-ShRNA mice (n = 12 astrocytes from n = 3 mice) versus Kir4.1-ShRNA mice (n = 14 astrocytes from n = 4 mice). **G***.* Representative membrane currents (top) of one lumbar GFP+ astrocyte of a Ctrl-ShRNA mouse before (left) and during (middle, right) bath application of Ba^2+^ (100 µM). **H***.* Averaged Ba^2+^-sensitive I/V curves of lumbar astrocytes from Ctrl-ShRNA mice (blue circles) or Kir4.1-ShRNA mice (blue circles) (n = 6 astrocytes from 2 Ctrl-ShRNA mice; n = 7 astrocytes from 2 Kir4.1-ShRNA mice) **I***.* Average Ba^2+^-sensitive current amplitude at -140mV in GFP+ astrocytes from Ctl-ShRNA mice (n = 6 from 2 mice) compared to Kir4.1-ShRNA mice (n = 7 from 2 mice). *P < 0.05, ** P<0.01, ***P < 0.001 (two-tailed Mann–Whitney test for C, F and I; comparison of linear regression slopes for E and H). Mean ± SEM. For detailed P values, see Source data file.

**Figure S7 (Related to Figure 7). The soma size from MMP9 (+) motoneurons is not modified in Kir4.1-ShRNA** **mice**. **A**. Representative confocal images of the immunofluorescent labelling of the cells nuclei with Hoechst (A1), the neuronal marker NeuN (A2), the fast-twitch fatigable (IIb) α-motoneuron marker, MMP9 (A3) from a lumbar slice of a P14 Ctrl-ShRNA mouse (top) and a P14 Kir4.1-ShRNA mouse (bottom). Note the merge signal is shown in A4. GM for grey matter, WM for white matter. MMP9 for matrix metalloproteinase-9. Scale bar is 40 µm. **B**. Quantification of the soma area of MMP9 (+) / NeuN (+) cells from Ctrl-ShRNA mice (88 cells from 3 mice) and from Kir4.1-ShRNA mice (105 cells from 3 mice). The error bars represent the SEM. The two-tailed unpaired t-test was used as data followed a normal distribution according to the Shapiro-Wilk test. ns for no significance. Note that all the MMP9 (+) cells in each field of view were measured. For detailed P values, see Source data file.

Ziskind-Conhaim, L., Wu, L., and Wiesner, E.P. (2008). Persistent sodium current contributes to induced voltage oscillations in locomotor-related hb9 interneurons in the mouse spinal cord. J Neurophysiol *100*, 2254-2264.

**supplemental experimental procedures**

### Experimental model

**Mice*.*** Mice of either sex (P3-P4 for fictive locomotion experiments, P5-P14 for Patch-clamp recording, P13-P16 for two-photon Ca^2+^ imaging, P12-P13 for swimming tests, P14-P15 for rotarod tests, P14-P18 for Catwalk tests) were housed under a 12h light/dark cycle in a temperature-controlled area with *ad libitum* access to water and food. Hb9-eGFP mice (C57/Bl6 background) expressing GFP in ventromedial oscillatory interneurons were provided by the Jackson laboratory (Strain #005029) and the *Aldh1L1*-eGFP mice (CD1 background) expressing GFP in astrocytes were kindly provided by Nathalie Rouach (Collège de France, CNRS, INSERM, Labex Memolife, Université PSL, Paris, France). Animals from different litters were used for each experiment. Sample sizes (number of spinal cords and/or cells) for each experiment are indicated in figure legends. All animal care and use were conformed to the French regulations (Décret 2010-118) and approved by the local ethics committee (Comité d’Ethique en Neurosciences INT-Marseille, CE71 Nb A1301404, authorization Nb 2018110819197361).

### Experimental procedures

### shRNA constructs. Specific shRNA sequence designed to knockdown Kir4.1 transcript (Cui et al. 2018) was incorporated into an adeno-associated viral (AAV) vector (serotype 9), which features a H1 promoter to drive shRNA expression and a CAG promoter to drive eGFP expression for identification of transduced cells (Cui et al. 2018). We also used a non-targeting shRNA sequence (Luciferase) which has no homology to any known genes in mouse as a control. AAV were produced following standard protocols (Molecular Cloning, A LABORATORY MANUAL, FOURTH EDITION, 2012) with minor modifications. Briefly, HEK293T cells were transfected with the three plasmids (pRC, pHelper and pAAV) using the MgCl_2_ method. After 60 to 72 hours, AAV particles from the cells and the medium were recovered and concentrated using Takara AAV-Purification All serotypes kit (Takara). We systematically included a digestion step with cryonase to degrade non-encapsidated viral genomes. Viral particles were quantified using quantitative PCR (Takara Titration kit). Vectors titers range was between 2x10^11^ and 3x10^12^ vectors/ml.

**Intrathecal vector delivery.** A minimally-invasive technique was used to micro-inject adeno-associated viral (AAV) vectors into the T13-L1 intervertebral space. Briefly, in pups cryoanesthetized at birth, the intervertebral space was widened by flexing the spine slightly. The tip of the microcapillary preloaded with the AAV particles was lowered into the center of the T13-L1 intervertebral space. A total volume of 2 µL /animal was then slowly injected by hand.

***Ex vivo* preparations and aCSF solutions*.*** *For the slice preparation*, the lumbar spinal cord was isolated in ice-cold (+4°C) artificial cerebro-spinal fluid (aCSF) solution composed of the following (in mM): sucrose (252), KCl (3), NaH_2_PO_4_ (1.25), MgSO_4_ (4), CaCl_2_ (0.2), NaHCO_3_ (26), D-glucose (25), pH 7.4. The lumbar spinal cord was then introduced into a 4% agar solution, quickly cooled, mounted in a vibrating microtome (Leica, VT1000S) and sliced (325 µm) through the L1–2 lumbar segments. Slices were immediately transferred into the holding chamber filled with bubbled (95% O_2_ and 5% CO_2_) **standard aCSF** composed of (in mM): NaCl (120), KCl (3), NaH_2_PO_4_ (1.25), MgSO_4_ (1.3), CaCl_2_ (1.2), NaHCO_3_ (25), D-glucose (20), pH 7.4, 30-32°C. After a 30-60 min resting period, individual slices were transferred to a recording chamber continuously perfused with standard aCSF heated to 32-34°C. The **oscillatory aCSF** characterized by higher extracellular K^+^ and lower Ca^2+^ [Brocard et al. 2013] was composed of the following (in mM): NaCl (120), KCl (6), NaH_2_PO_4_ (1.25), MgSO_4_ (1.3), CaCl_2_ (0.9), NaHCO_3_ (25), D-glucose (20). The rhytmogenic cocktail (Ziskind-Conhaim et al., 2008) that we called **neuromodulator cocktail** in this study, was composed of the following (in µM): tetrodotoxin (TTX, 0.5-1), N-methyl-D-aspartate (NMDA, 20), 5*-*hydroxytryptamine (5-HT, 20), dopamine (DA, 50).

*For the whole-spinal cord preparation,* the spinal cord was transected at T8-9, isolated and transferred with intact dorsal and ventral roots to the recording chamber. The tissue was continuously bubbled (95% O_2_ and 5% CO_2_) and perfused with heated (~27-28°C) **standard aCSF** solution composed of (in mM): 120 NaCl, 4 KCl, 1.25 NaH_2_PO_4_, 1.3 MgSO_4_, 1.2 CaCl_2_, 25 NaHCO_3_, 20 D-glucose, pH 7.4. The **oscillatory aCSF** was composed as described above. The **neuromodulator cocktail** was composed of the following (in µM): N-methyl-DL-aspartate (NMA, 12), 5*-*hydroxytryptamine (5-HT, 5)*.*

***Ex vivo* recordings.** *For the slice preparation,* whole-cell patch-clamp recordings of L1-L2 ventromedial interneurons and astrocytes were performed using a Multiclamp 700B amplifier (Molecular Devices). The apparatus was equipped with two headstages, allowing recordings of pairs of cells. Patch electrodes (4-5 MΩ for neurons) were pulled from borosilicate glass capillaries (1.5 mm OD, 1.12 mm ID; World Precision Instruments) on a Sutter P-97 puller (Sutter Instruments Company) and filled with an intracellular solution (in mM): K^+^-gluconate (140), NaCl (5), MgCl_2_ (2), HEPES (10), EGTA (0.5), ATP (2), GTP (0.4), pH 7.3. For GFP+ astrocyte recordings, pipettes (7-9 MΩ) were filled with an intracellular solution (in mM): K^+^-gluconate (105), NaCl (10), KCl (20), MgCl_2_ (0.15), HEPES (10), EGTA (0.5), ATP (4), GTP (0.3), pH 7.3. In some recordings, 30 mM of BAPTA was added in the pipette solution to chelate intracellular free Ca^2+^ from the intracellular stores. In some experiments, GFP negative astrocytes were identified on the basis of their morphology (i.e., small somata with a diameter of <10 μm surrounded by many processes) and their electrophysiological properties (i.e. a hyperpolarized RMP (<-75 mV), a linear I/V relationship, no spike in response to increased injection currents and a low input resistance (<100 MΩ)). When GFP negative, some astrocytes were filled with Alexa Fluor® 488 or 594 (25µM) diluted in the intracellular solution. Pipette and neuronal capacitive currents were canceled and, after breakthrough, the series resistance was compensated and monitored. Recordings were digitized on-line and filtered at 10 kHz (Digidata 1550B, Molecular Devices). All experiments were designed to gather data within a stable period (i.e., at least 1-2 min after establishing whole-cell access).

*For the whole spinal cord preparation*, motor outputs were recorded from L2 lumbar ventral roots by means of glass suction electrodes connected to an AC-coupled amplifier. Signals from the electrodes placed in contact with the ventral root recordings were amplified (×2,000), high-pass filtered at 70 Hz, low-pass filtered at 3 kHz, and sampled at 10 kHz. Custom-built amplifiers enabled simultaneous online rectification and integration (100 ms time constant) of raw signals. Locomotor-like activity was induced by (i) rising [K^+^]_e_ to 6mM and decreasing [Ca^2+^]_e_ to 0.9mM (oscillatory aCSF, see above) or (ii) bath application of the neuromodulator cocktail composed of N-methyl-DL aspartate (NMA, 12 μM) and 5-hydroxytryptamine (5-HT, 5 μM).

**Two-photon Ca^2+^ imaging.** Two-photon fluorescence measurements were obtained with a dual-scanhead two-photon microscope (FemtoS-Dual, Femtonics Ltd, Budapest, Hungary) and made using an Olympus XLUMPlanFL N 20X, 1.00 (Olympus America, Melville, NY). Two-photon excitation of Gcamp6f and jRGECO1a was simultaneously evoked with a femtosecond pulsed laser (Chameleon Ultra II; Coherent, Santa Clara, CA) tuned to 960 nm. The microscope system was controlled by MESc acquisition software (https://femtonics.eu/femtosmart-software/, Femtonics Ltd, Budapest, Hungary). A single acquisition plane was selected and full-frame imaging was started in a resonant scanning mode at 30.5202 Hz. Scan parameters were [pixels/line × lines/frame (frame rate in Hz)]: [512 × 519 (30.5202)]. Scanning area was 468 µm * 474 µm. This microscope was equipped with two detection channels for fluorescence imaging.

**Immunohistochemistry.** Spinal cords of 10 to 12-days-old mice were dissected out and fixed for 5-6 h in 4% paraformaldehyde (PFA), then rinsed in phosphate buffered saline (PBS) and cryoprotected overnight in 20% sucrose at 4°C. Spinal cords were frozen in OCT medium (Tissue Tek) and 30 μm cryosections were collected from the L1-L2 segments. After having been washed in PBS 3×5 min, the slides were incubated for 1 h in a blocking solution (BSA 1%, Normal Donkey Serum 3% in PBS) with 0.2% triton X-100 and for 48 h at 4 °C in a humidified chamber with the primary antibody anti-Kir4.1 (Alomone, #APC035, 1/1000) or for 12 h at 4 °C with the primary antibody anti-NeuN (Merck Millipore, #MAB377, 1/200), or for 12 h at 4 °C with the primary antibody anti-MMP9 (Merck Sigma-Aldrich, #M9570, 1/500). All antibodies were diluted in the blocking solution with 0.2% triton X-100. Slides were then washed 3×5 min in PBS and incubated for 2 h with an Alexa Fluor® Plus 555- conjugated secondary antibody (Invitrogen #A32794, 1/400), an Cy5- conjugated secondary antibody (Jackson ImmunoResearch, #715-175-151, 1/400) or Alexa Fluor® Plus 555- conjugated secondary antibody (Invitrogen #A32816, 1/400) diluted in the blocking solution. After 3 washes of 5 min in PBS, they were mounted with a gelatinous aqueous medium containing Hoechst 33342 (ThermoFisher, #62249, 1/10000, initial concentration: 20mM). Images were acquired using a confocal microscope (LSM700, Zeiss) equipped with either a 20x air objective, a 40x oil objective or a 63x oil objective and processed with the Zen software (Zeiss).

**Kir4.1 protein analysis by capillary Western Blot.** The lumbar parts of the spinal cords were dissected in aCSF at 4°C and conserved at -80°C until protein extraction. Tissues were homogenized in ice-cold lysis buffer (250 mM sucrose, 3.9 mM Tris pH7.5, 10 mM iodoacetamide) supplemented with protease inhibitors (cOmplete™, Mini, EDTA-free Protease Inhibitor Cocktail, Roche 11836170001). Unsolubilized material was pelleted by centrifugation at 7,000 x g for 5 min at 4°C and discarded. The supernatants were subjected to an additional centrifugation step at 19,000 x g for 70 min at 4°C. The resulting pellets corresponding to the membrane-enriched fraction were resuspended in ice-cold lysis buffer (PBS 1X, 1% IGEPAL® CA-630 1%, SDS 0.1%) supplemented with protease inhibitors (cOmplete™, Mini, EDTA-free Protease Inhibitor Cocktail, Roche 11836170001). Protein concentrations were determined using the Pierce™ BCA Protein Assay Kit (ThermoFisher 23227). Kir4.1 expression was then analyzed using the 12-230 kDa separation module (SM-W004 ProteinSimple) on an automated capillary western blotting system (‘Jess’ ProteinSimple) according to the manufacturer’s protocol with small modifications. Samples were not implemented with DTT and they were not submitted to heat denaturation in order to keep proteins in a more ‘native’ form. A total protein concentration of 0.1 mg/mL was used. Samples were probed with a rabbit polyclonal Kir4.1 antibody (1:60; Alomone APC-035) and revealed with the appropriate detection module (DM-001 ProteinSimple). Loaded samples were normalized to their own total protein content using the ‘total protein detection module’ (DM-TP01 ProteinSimple).

**Assessment of *in vivo* motor behaviors.** *Walking.* The CatWalkXT (Noldus Information Technology, Netherlands) was used to measure walking performance. Each animal walked freely through a corridor on a glass walkway illuminated with beams of light from below. A successful walking trial was defined as having the animal walk at a steady speed (no stopping, rearing, or grooming), and three to five successful trials were collected per animal. The footprints were recorded using a camera positioned below the walkway, and footprint classification was manually corrected to ensure accurate readings. The paw print parameters were then analyzed using the CatWalk software (see data analysis). *Rotarod test.* Mice were placed on a rotarod (Bioseb) rotating at a accelerating from 4 to 40 rpm over a span of 5 min. Mice were given 3 trials with a 60 seconds inter-trial interval. *Swimming.* Mice were gently placed individually in the center of the tank (~19000 cm^3^) filled with heated water (30-33°C). Swimming distance and velocity were quantified during three consecutive 90s periods. Each trial was spaced by a 10 min interval. At the end of the trial, the mouse was immediately removed from the tank, dried off with a paper towel, and returned to its home cage. Swimming parameters tracking and analysis were performed by using an automated video-tracking Ethovision system (Noldus). All behavioral experiments were carried out with the experimenter blind to genotype.

**Drugs.** All solutions were oxygenated with 95% O_2_/5% CO_2_. All salt compounds, N-methyl-D-aspartic acid (NMDA, 20µM; #M3262), dopamine hydrochloride (DA, 50µM; #H8502) N-methyl-DL-aspartic acid (NMA, 12µM; #M2137), 5-hydroxytryptamine creatinine sulfate (5-HT, 5-20 µM; #S2805), Barium chloride dihydrate (Ba^2+^, 100µM; #217565), and 1,2-bis(o-aminophenoxy)ethane-N,N,N′,N′-tetraacetic acid (BAPTA, 30mM; #A9801) were obtained from Sigma-Aldrich. Tetrodotoxin (TTX, 0.5-1µM; #1078) from Tocris Bioscience. All drugs were dissolved in water and added to the aCSF.

### Data analysis

**Electrophysiological data analyses** were analyzed off-line with Clampfit 10.7 software (Molecular Devices). For *intracellular recordings*, several basic criteria were set to ensure optimum quality of intracellular recordings. Only cells exhibiting a stable RMP, access resistance (no > 20% variation) and an action potential amplitude larger than 40 mV were considered. Passive membrane properties of cells (neurons and astrocytes) were measured by determining from the holding potential the largest voltage deflections induced by small current pulses that avoided activation of voltage-sensitive currents. We determined input resistance by the slope of linear fits to voltage responses evoked by small positive and negative current injections. All reported membrane potentials were corrected for liquid junction potentials (+14 mV for neurons and +11.4 mV for astrocytes). To activate Kir-currents in spinal astrocytes, voltages were repeatedly stepped from -140 to 0 mV, in 10 mV increments. Astrocytes were initially clamped at their RMP. 100 µM Ba^2+^ was bath applied to the slice approximately 5 minutes after the start of the recording. The current amplitude was measured as the difference between the baseline level before the initial voltage step, and the mean amplitude over a 10 ms window starting 10 ms after onset of the hyperpolarizing voltage step. The Ba^2+^ sensitive current was obtained by subtracting the current after Ba^2+^ application from the current before application.

For *extracellular recordings*, alternating activity between right/left L2 recordings was taken to be indicative of fictive locomotion. To characterize locomotor burst parameters (during 1-2 mins of steady state), raw extracellular recordings from ventral roots were rectified, integrated and resampled at 50 Hz. Peak amplitude and duration of locomotor burst were measured by using a threshold function and the cycle period was calculated by measuring the time between the first two peaks of the auto-correlogram. The coupling between right/left L2 was estimated by measuring the correlation coefficient of the cross-correlogram at zero phase lag.

**Analysis of firing patterns.** Firing patterns of GFP+ interneurons were characterized as tonic or rhyth­mic bursting discharge in response to ions variations before or after pharmacological or genetic blockade of Kir.1 channels. Tonic discharge typically consisted of low-frequency single spikes (<10 Hz) separated by regular but relatively long intervals (interspike intervals: ISI of 200–600 ms). We defined bursts as depolarizing plateaus over-ridden by at least two actions potentials followed by a silent period. The period separating the beginning of two con­secutive plateaus was used to calculate the inter-burst frequency in intracellular recordings. Bursting frequency was calculated only in neurons that displayed a stable rhythmic discharge for more than five cycles.

**Two-photon Ca^2+^ imaging analyses.** Fluorescent time series depicting the Ca^2+^ changes occurring *ex vivo* in both the neurons and the astrocytes were analyzed with the MESc data acquisition software (Femtonics Ltd, Budapest, Hungary) and the MES curve analyzer tool. Briefly, regions of interest (ROIs) were manually selected based on the appearance of Ca^2+^ transients in the time series images. A signal was declared as a Ca^2+^ transient if it exceeded the baseline by greater than twice the baseline noise (SD). ROIs coincide with (i) active soma or active dendritic components for neurons and (ii) active microdomains or territories from the astrocytic processes in control conditions. Astrocyte territory sizes were estimated by measuring the area of a ROI that surrounded the largest fluorescence projection profile of bushy astrocytes gathered from a series of z-stack images. The astrocytic ROIs were separated at least by a distance of 60µm. The raw fluorescence Ca^2+^ traces were extracted to Excel and analyzed using custom Matlab scripts. They were transformed to show fluorescence change according to Eq. (1):

ΔF/F=(F_1_-F_0_)/F_0_	(1)

where *F* is the fluorescence at any given point in time, and F_0_ is the mean fluorescence value for the 5 to 10 s range, preceding the bath perfusion of the oscillatory aCSF or the neuromodulator cocktail. Following this, the signal-to-noise ratio (SNR) is:

SNR = (ΔF/F)_peak_/ σF

where σF is the standard deviation of the baseline period. We took those events as a Ca^2+^ response that exceeded the threshold of two times the standard deviation of the baseline period. The peak amplitude, duration and frequency of the Ca^2+^ transients were analysed by using a Matlab script. In each FOV where dorsal is up and ventral is down, all astrocytic Ca^2+^ transients above the central canal were considered as dorsal astrocytic activity and distinguished from the ventral astrocytic Ca^2+^ transients.

***In vivo* behavioral analyses.** For walking, the CatWalk XT software (Noldus Information Technology, Netherlands) was used to measure a broad number of spatial and temporal gait parameters in several categories. These include i) dynamic parameters related to individual paw prints, such as duration of the step cycle with the respective duration of the swing and stance phases; ii) parameters related to the position of paw prints with respect to each other, for example the stride length (distance between two consecutive placement of the same paw) and the base of support (the width between the paw pairs); iii) parameters related to time-based relationships between paw pairs, as well as step patterns. These parameters were calculated for each run and for each paw. In the rotarod test the average latency of the subject to fall was recorded (in seconds). A video tracking system (Noldus Information Technology, Wageningen, The Netherlands) was used to measure swimming performance. We ensured there was a smooth tracking curve and that the center point of the animal remained stable before analysis took place. Distance and velocity during the test periods were scored. All behavioral tests were carried out with the experimenter blind to genotype.

### Statistics

No statistical method was used to predetermine sample size. Group measurements were expressed as means ± S.E.M. When two groups were compared, we used the Mann-Whitney test. Fisher test was used to compare the percentages of oscillatory neurons. When two conditions (control vs drugs) were compared, we used the Wilcoxon matched pairs test. We also used a one-way or two-way ANOVA tests for multiple comparisons. For all statistical analyses, the data met the assumptions of the test and the variance between the statistically compared groups was similar. The level of significance was set at p < 0.05. Statistical analyses were performed using Graphpad Prism 9 software. They are all indicated in Figure legends.
