## Supplementary figures and images for "Astrocytes regulate locomotion by orchestrating neuronal rhythmicity in the spinal network via potassium clearance"

### Supplemental Figure 1

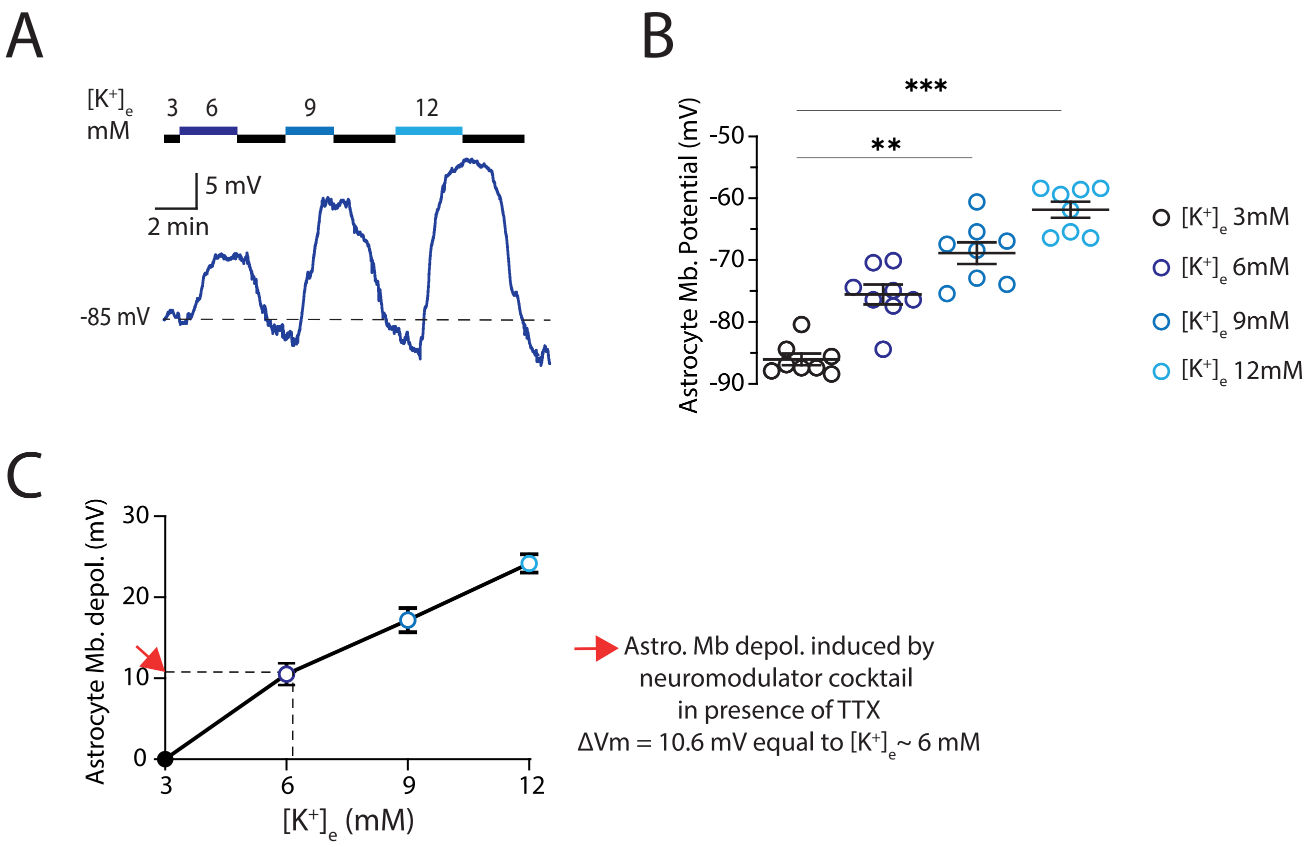

### Supplemental Figure 2

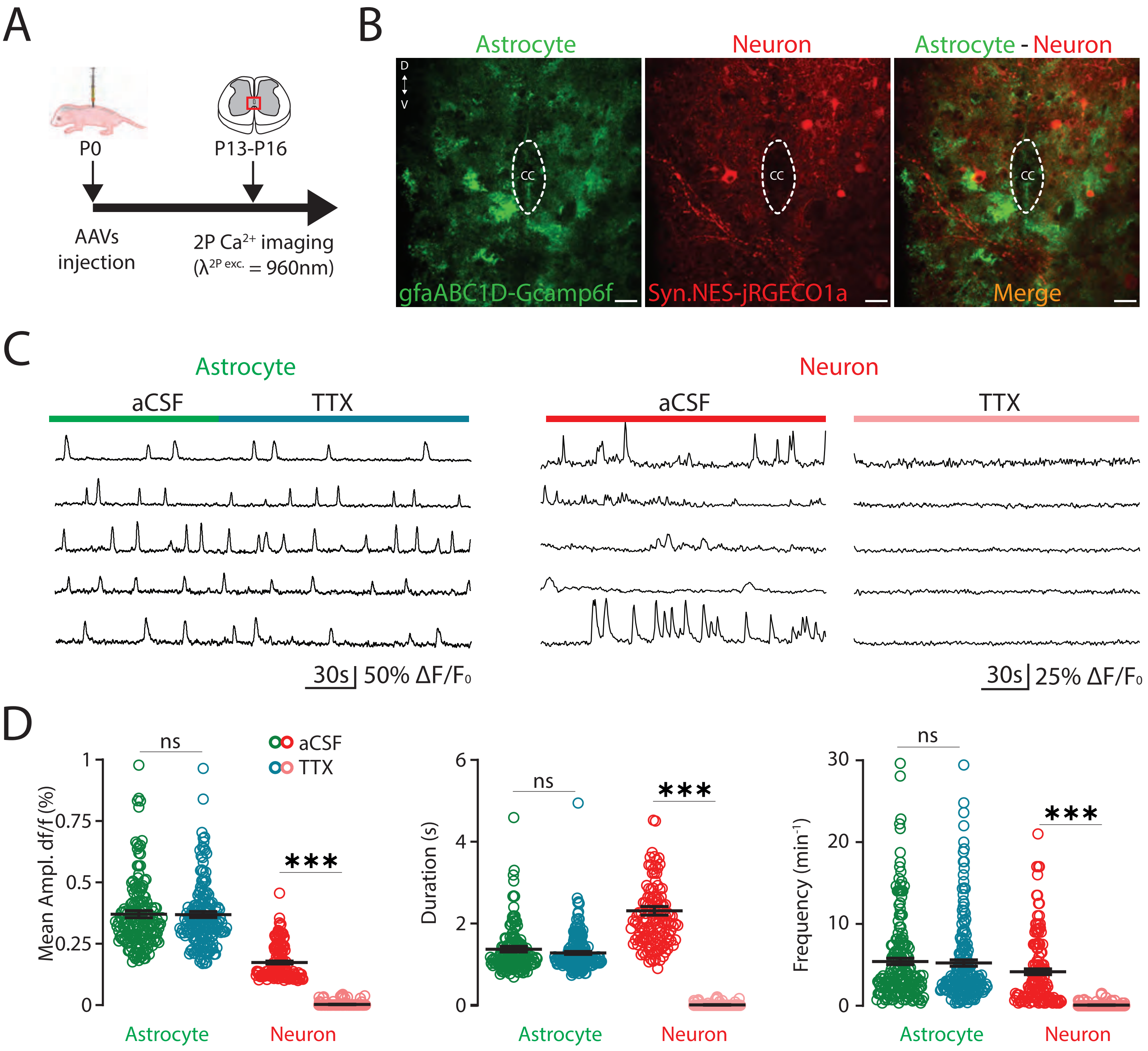

### Supplemental Figure 3

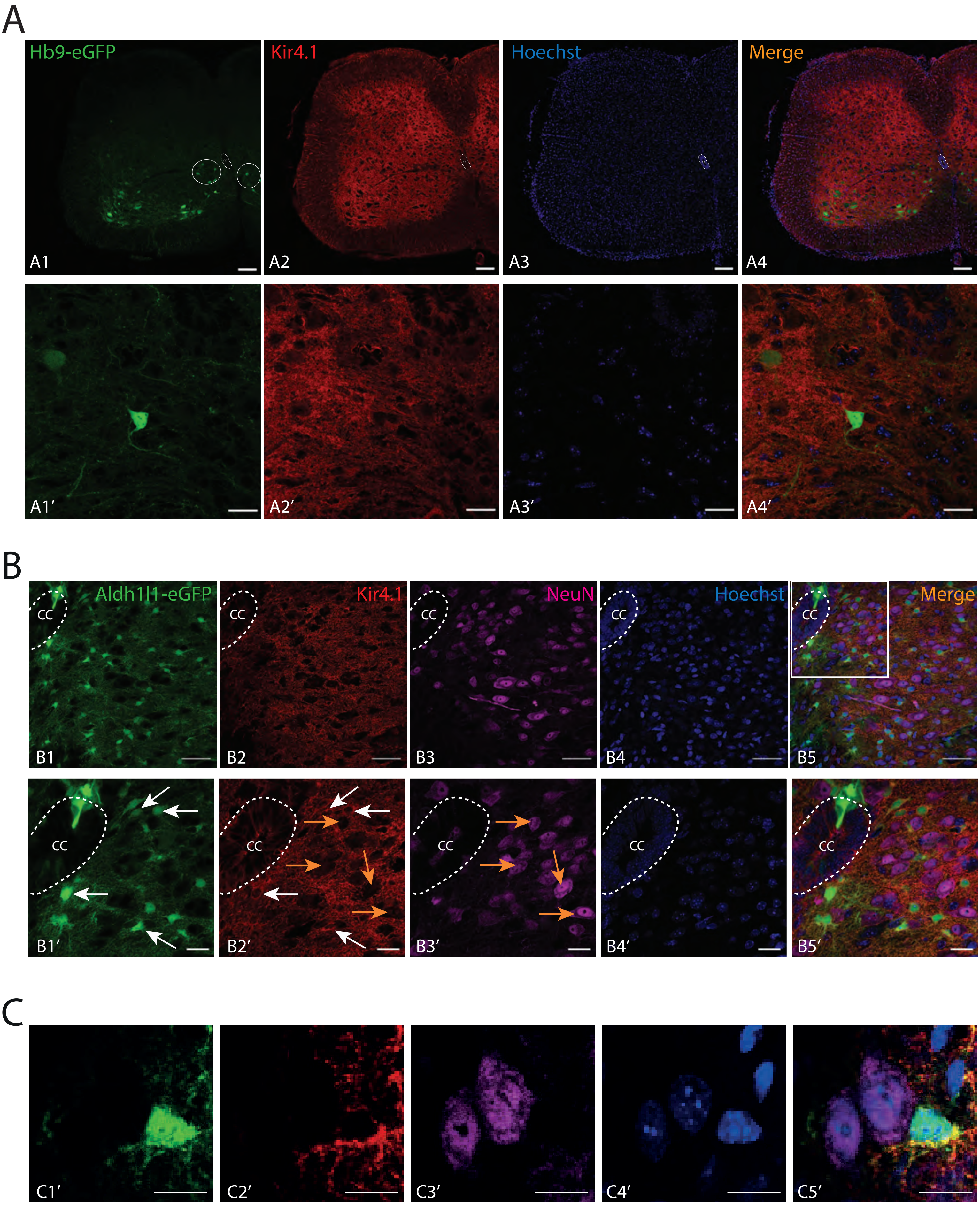

### Supplemental Figure 4

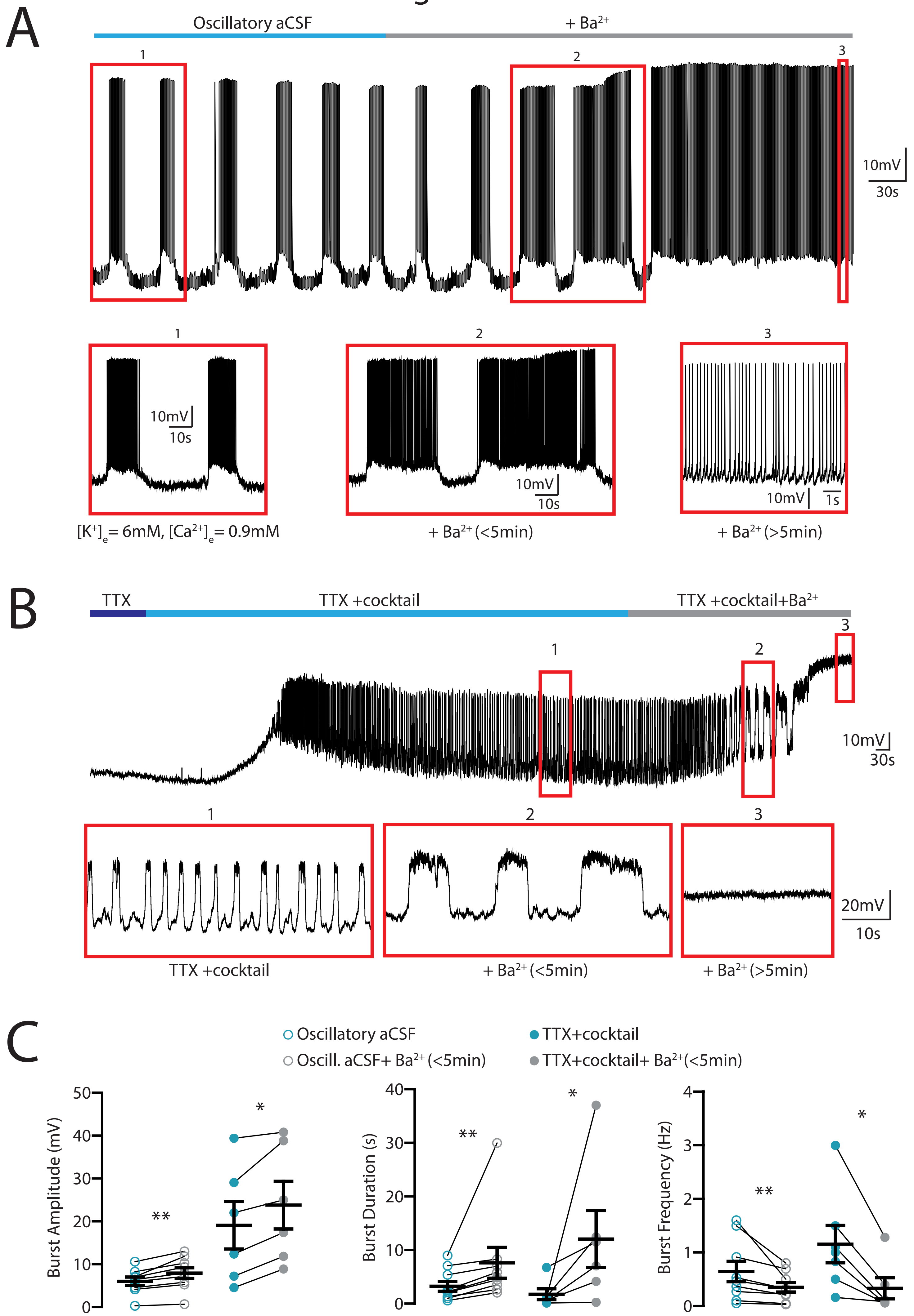

### Supplemental Figure 5

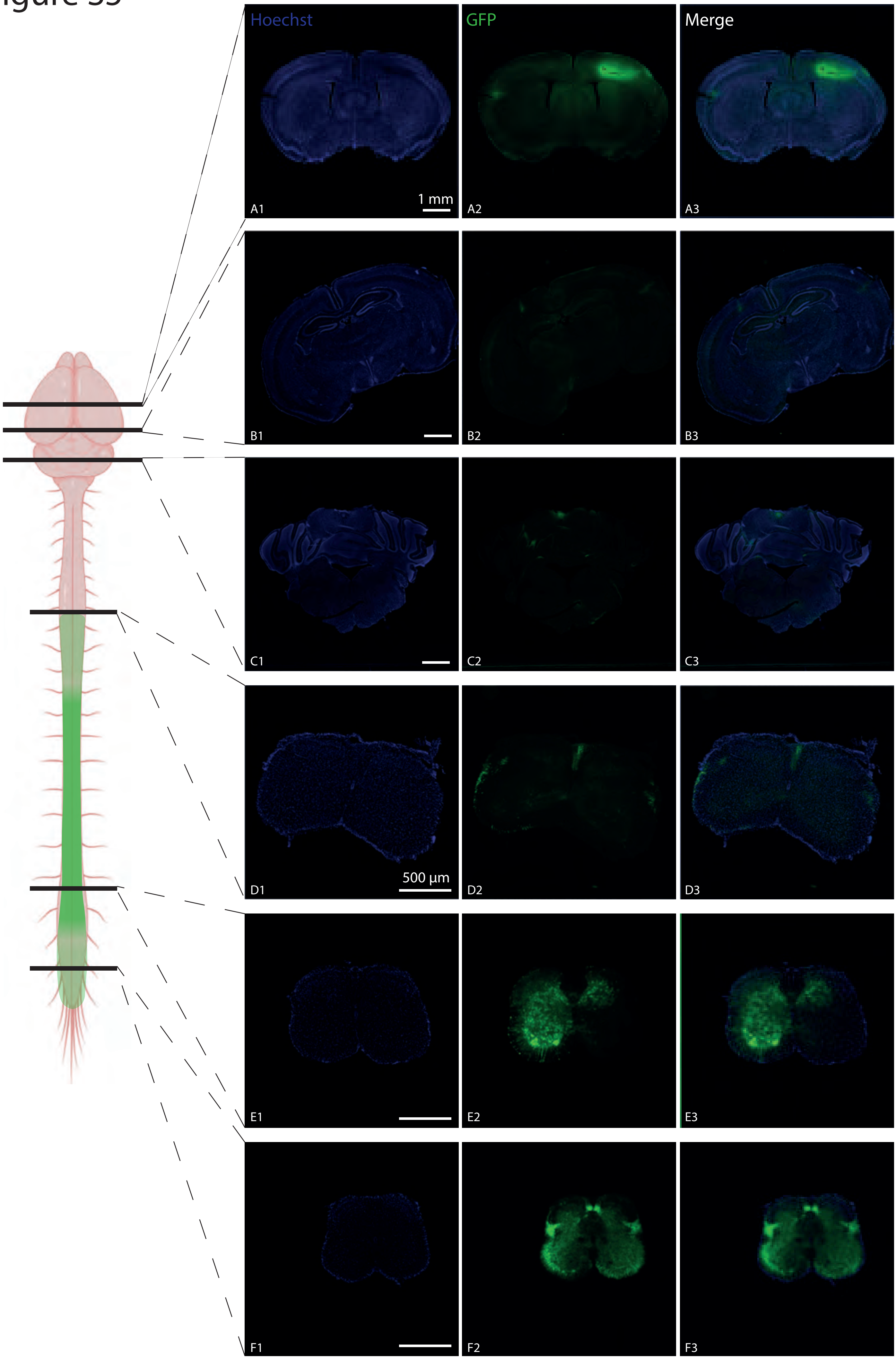

### Supplemental Figure 6

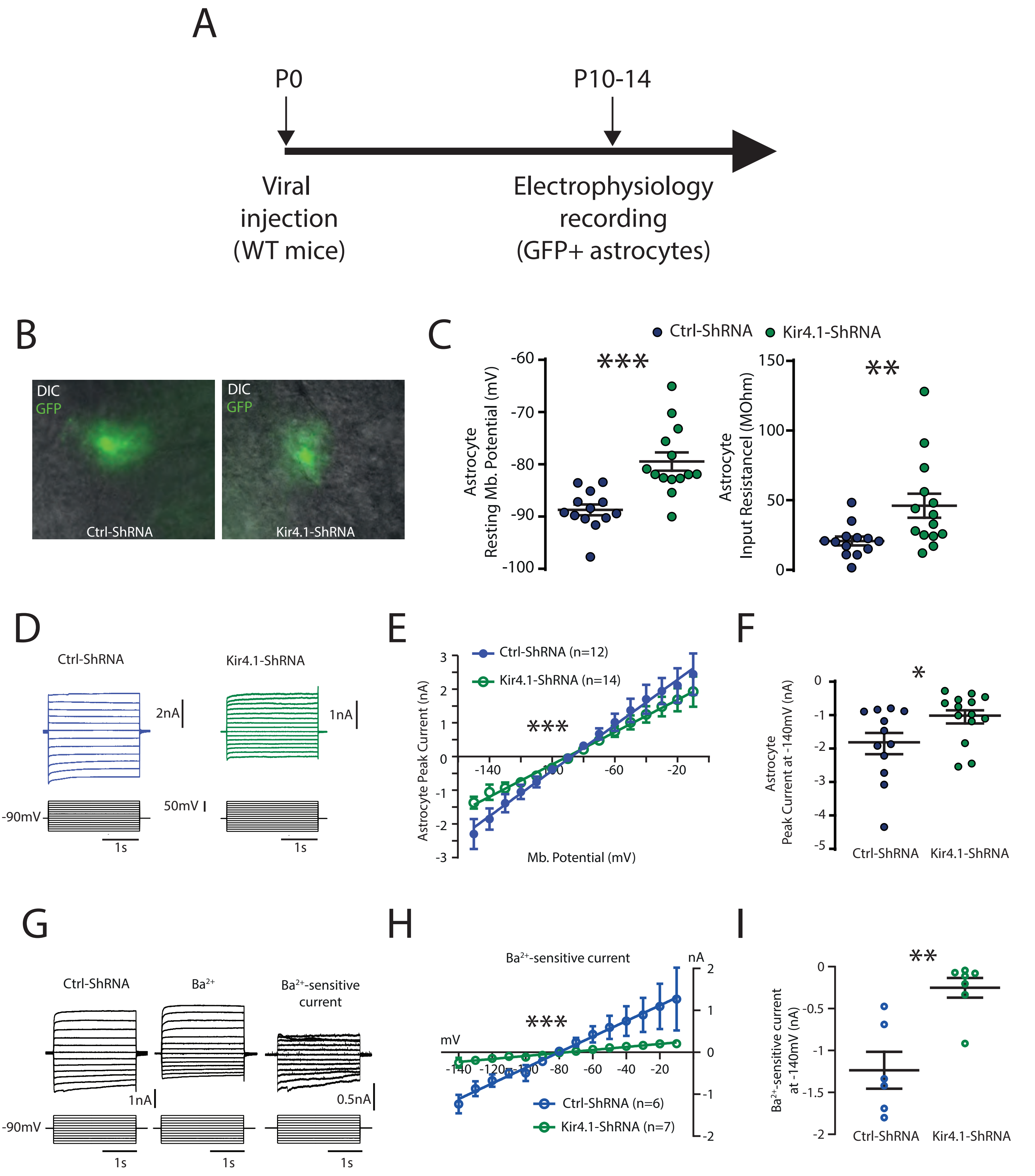

### Supplemental Figure 7

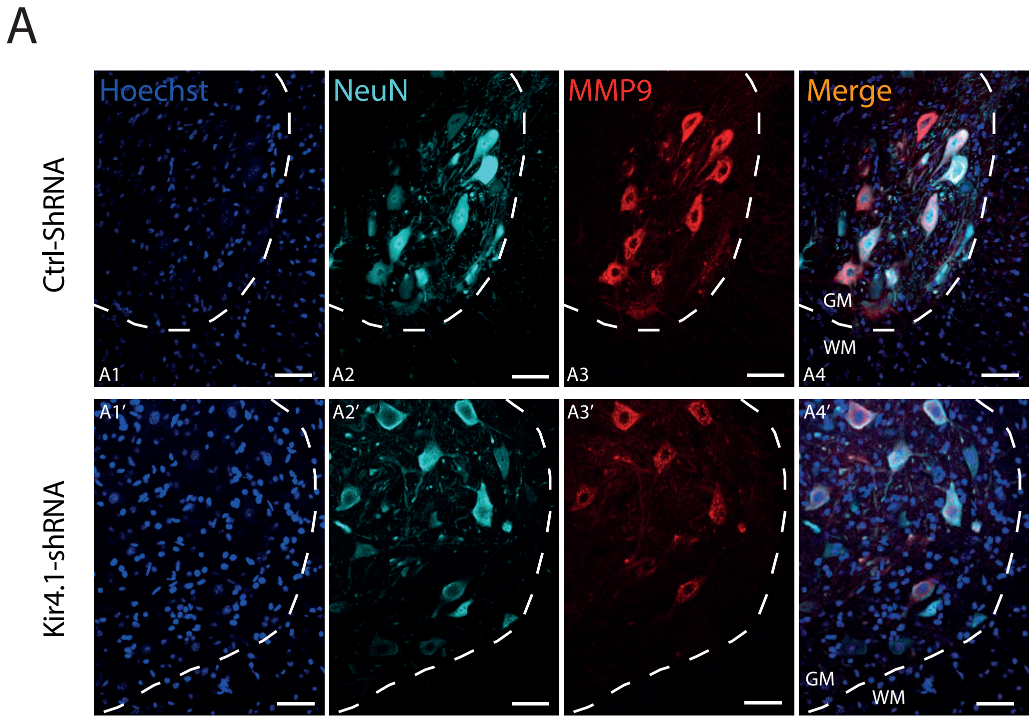

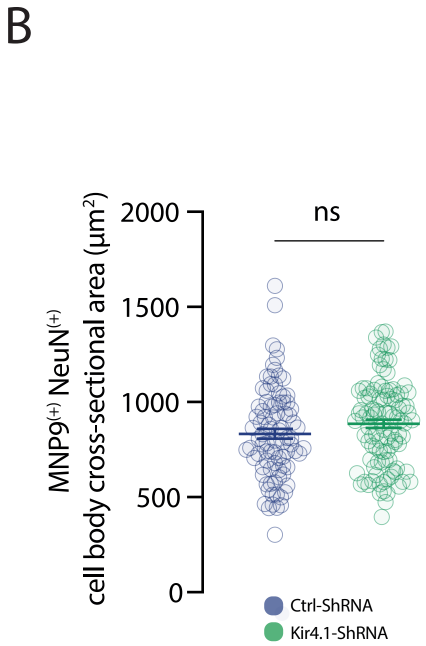
