## Supplementary material for "Astrocytes regulate locomotion by orchestrating neuronal rhythmicity in the spinal network via potassium clearance": Data Source File

from

Available on:

https://amubox.univ-amu.fr/s/bcyGfGwJo5RBeXe

**Fig. 1C** Data Source File

| | age | Vm astro standard aCSF (mV) | Vm astro Oscill. aCSF (mV) | delta Vm (mV) |
|---|---|---|---|---|
| mouse#1 | P11 | -90.4 | -71.31 | 19.09 |
| mouse#2 | P5 | -96.77 | -76.1 | 20.67 |
| mouse#2 | P5 | -93.66 | -79.58 | 14.08 |
| mouse#3 | P6 | -87.89 | -72.89 | 15 |
| mouse#3 | P6 | -88 | -75.55 | 12.45 |
| mouse#4 | P11 | -85.29 | -67.22 | 18.07 |
| mouse#4 | P11 | -84.18 | -67.64 | 16.54 |
| mouse#5 | P12 | -78.53 | -68.47 | 10.06 |
| mouse#6 | P7 | -96.4 | -73.4 | 23 |
| mouse#6 | P7 | -88.4 | -77.4 | 11 |
| mouse#7 | P5 | -84.4 | -76.4 | 8 |
| mouse#8 | P6 | -78.4 | -70.4 | 8 |
| mouse#9 | P9 | -86.4 | -77.4 | 9 |
| mouse#10 | P7 | -94.2 | -84.4 | 9.8 |
| mouse#11 | P8 | -92.4 | -79.4 | 13 |
| mouse#12 | P8 | -88 | -80.98 | 7.02 |
| mouse#13 | P13 | -95.34 | -78.58 | 16.76 |
| mouse#14 | P6 | -93.7 | -81.53 | 12.17 |
| mouse#14 | P6 | -79.25 | -63.5 | 15.75 |
| mouse#15 | P7 | -77.98 | -62.84 | 15.14 |
| mouse#15 | P7 | -88.26 | -73.82 | 14.44 |
| mouse#16 | P8 | -81.5 | -63.4 | 18.1 |
| | MEAN | -87.69772727 | -73.73681818 | 13.9609091 |
| | SEM | 1.288 | 1.328 | 0.92932618 |

**Wilcoxon matched-pairs signed rank test**

| | |
|---|---|
| P value | <0.0001 |
| P value summary | **** |

**Fig. 1E** Data Source File

| | age | Vm astro TTX (mV) | Vm astro TTX+cocktail (mV) | delta Vm (mV) |
|---|---|---|---|---|
| mouse#1 | P10 | -84.92 | -76.42 | 8.5 |
| mouse#1 | P10 | -96.16 | -89.32 | 6.84 |
| mouse#1 | P10 | -86.46 | -81.86 | 4.6 |
| mouse#2 | P12 | -91.17 | -85.62 | 5.55 |
| mouse#3 | P13 | -91.32 | -67.21 | 24.11 |
| mouse#3 | P13 | -83.26 | -73.95 | 9.31 |
| mouse#4 | P7 | -89.47 | -82.2 | 7.27 |
| mouse#4 | P7 | -86.73 | -68 | 18.73 |
| | MEAN | -88.68625 | -78.0725 | 10.61375 |
| | SEM | 1.476 | 2.847 | 2.469 |

**Wilcoxon matched-pairs signed rank test**

| | |
|---|---|
| P value | 0.0078 |
| P value summary | ** |

**Fig. 2B** Data Source File

| Neuron | Oscill. aCSF | Oscillation nb | Ampl. (mV)mV) | Oscillation duration (s)) | Frequency (Hz) (Hz) |
|---|---|---|---|---|---|
| mouse#1 | ROI#1 | 7 | 0.34613043 | 3.51 | 0.07575758 |
| P13 | ROI#2 | 8 | 1.2243481 | 2.58 | 0.08888889 |
| | ROI#3 | 12 | 0.8 | 2.38 | 0.10714286 |
| mouse#2 | ROI#1 | 36 | 0.5887435 | 1.87 | 0.2 |
| P13 | ROI#2 | 11 | 0.55273931 | 1.58 | 0.10576923 |
| | ROI#3 | 5 | 0.5 | 18 | 0.03333333 |
| | ROI#4 | 29 | 0.97432955 | 1.35 | 0.35626536 |
| | ROI#5 | 43 | 0.5206174 | 1.94 | 0.17269076 |
| | ROI#6 | 22 | 0.19712567 | 1.37 | 0.30136986 |
| | MEAN | 19.2222222 | 0.6338 | 3.842 | 0.1601 |
| | SEM | | 0.1057 | 1.784 | 0.03617 |

| Neuron | TTX+Cocktail | Oscillation nb | Ampl. (mV)mV) | Oscillation duration (s)) | Frequency (Hz) (Hz) |
|---|---|---|---|---|---|
| mouse#1 | ROI#1 | 5 | 0.2 | 7.62 | 0.11111111 |
| P16 | ROI#2 | 10 | 0.4 | 6.02 | 0.14705882 |
| mouse#2 | ROI#1 | 7 | 0.8 | 6.97142857 | 0.10294118 |
| P16 | ROI#2 | 9 | 0.6 | 8.1 | 0.1125 |
| mouse#3 | ROI#1 | 4 | 0.4 | 7.9 | 0.04878049 |
| P16 | ROI#2 | 25 | 0.82969904 | 2.49 | 0.18656716 |
| | ROI#3 | 24 | 0.46345903 | 3.53 | 0.14634146 |
| | ROI#4 | 10 | 0.62758087 | 6.71 | 0.0729927 |
| | ROI#5 | 27 | 0.61934994 | 3.09 | 0.16564417 |
| | ROI#6 | 38 | 0.42188235 | 2.16 | 0.2183908 |
| | ROI#7 | 19 | 0.40430752 | 3.9 | 0.11515152 |
| | ROI#8 | 41 | 0.34604624 | 1.64 | 0.25949367 |
| | ROI#9 | 32 | 0.49020699 | 2.52 | 0.2 |
| | ROI#10 | 15 | 0.19568064 | 3.25 | 0.19736842 |
| | ROI#11 | 18 | 0.45299989 | 2.98 | 0.17142857 |
| mouse#4 | ROI#1 | 19 | 0.41856026 | 5.79 | 0.10555556 |
| P13 | ROI#2 | 29 | 0.27169718 | 2.26 | 0.23577236 |
| | ROI#3 | 13 | 0.20200932 | 7.8 | 0.05701754 |
| | ROI#4 | 12 | 0.68017693 | 3.08 | 0.13636364 |
| | ROI#5 | 20 | 0.56751676 | 3.52 | 0.12658228 |
| | ROI#6 | 45 | 0.28696338 | 2.55 | 0.20454545 |
| | ROI#7 | 17 | 0.32632338 | 3.16 | 0.27287319 |
| | ROI#8 | 19 | 0.26380797 | 3.49 | 0.17272727 |
| | ROI#9 | 9 | 0.45227691 | 8.5 | 0.04285714 |
| | MEAN | 19.4583333 | 0.4467 | 4.543 | 0.1504 |
| | SEM | | 0.03648 | 0.4652 | 0.0131 |

**Fig. 2C (a)**

Data Source File

| | standard aCSF Amplitude (df/f) | oscill. aCSF Amplitude (df/f) | standard aCSF Duration (ms) | oscill. aCSF Duration (ms) | standard aCSF Frequency (Events Nb/min) | oscill. aCSF Frequency (Events Nb/min) |
|---|---|---|---|---|---|---|
| 62 / UG / ROI: | 0.30558305 | 0.33951559 | 1128.17373 | 1207.17971 | 7.57894737 | 9.05660377 |
| 54 / UG / ROI: | 0.3194721 | 0.34816121 | 1122.09818 | 1166.55537 | 5.05263158 | 3.16981132 |
| 57 / UG / ROI: | 0.0617747 | 0.34361267 | 935.08182 | 2143.92658 | 0.11232222 | 0.22641509 |
| 29 / UG / ROI: | 0.25799111 | 0.37482399 | 897.18616 | 1157.03682 | 1.26315789 | 0.90566038 |
| 36 / UG / ROI: | 0.48329971 | 0.4208317 | 1580.84078 | 1210.23428 | 2.52631579 | 6.11320755 |
| 10 / UG / ROI: | 0.40103417 | 0.42238674 | 1114.86228 | 1168.05353 | 1.26315789 | 4.98113208 |
| 23 / UG / ROI: | 0.45856937 | 0.73059458 | 958.225641 | 875.108929 | 5.05263158 | 3.16981132 |
| 38 / UG / ROI: | 0.28136669 | 0.43265465 | 1525.09266 | 974.321875 | 1.89473684 | 1.13207547 |
| 59 / UG / ROI: | 0.39910671 | 0.57730222 | 990.256529 | 842.620578 | 3.78947368 | 8.1509434 |
| 1 / UG / ROI:{ | 0.31777791 | 0.44934741 | 2140.32385 | 1028.67896 | 0.63157895 | 1.81132075 |
| 2 / UG / ROI:{ | 0.28002862 | 0.39255407 | 2192.83275 | 900.191795 | 0.63157895 | 1.35849057 |
| 14 / UG / ROI: | 0.07447666 | 0.38150498 | 1827.36063 | 2115.11526 | 0.12325547 | 0.22641509 |
| 19 / UG / ROI: | 0.31695923 | 0.37288578 | 1132.481 | 1785.18543 | 5.05263158 | 4.98113208 |
| 44 / UG / ROI: | 0.33494421 | 0.35487313 | 1082.98892 | 1261.60171 | 3.15789474 | 3.16981132 |
| 30 / UG / ROI: | 0.08921106 | 0.3373051 | 902.490768 | 1561.3413 | 0.23155568 | 0.67924528 |
| 15 / UG / ROI: | 0.61520063 | 1.80767496 | 1845.36551 | 1020.95423 | 0.63157895 | 0.45283019 |
| 48 / UG / ROI: | 0.05561282 | 0.52289633 | 1537.80459 | 1030.10214 | 0.32115648 | 0.67924528 |
| 40 / UG / ROI: | 0.31223667 | 0.37120605 | 1233.02674 | 1237.36187 | 7.57894737 | 7.9245283 |
| 11 / UG / ROI: | 0.55961482 | 0.62116981 | 904.818125 | 911.660405 | 4.42105263 | 5.88679245 |
| 75 / UG / ROI: | 0.07289278 | 0.33938273 | 754.015104 | 982.808277 | 0.51589985 | 1.13207547 |
| 3 / UG / ROI:{ | 0.07778449 | 0.37461131 | 942.518881 | 1479.42109 | 0.21315564 | 0.45283019 |
| 58 / UG / ROI: | 0.27406784 | 0.35634435 | 1890.13838 | 1276.13832 | 1.26315789 | 3.16981132 |
| 46 / UG / ROI: | 0.43977932 | 0.5531526 | 896.756354 | 1073.40934 | 13.2631579 | 13.1320755 |
| 37 / UG / ROI: | 0.41325886 | 0.4711271 | 811.778206 | 794.324869 | 20.2105263 | 25.5849057 |
| 28 / UG / ROI: | 0.05773295 | 0.28298833 | 1159.68315 | 782.097261 | 0.11332554 | 0.22641509 |
| 63 / UG / ROI: | 0.30582343 | 0.3939075 | 1372.49918 | 883.306468 | 3.15789474 | 0.45283019 |
| 78 / UG / ROI: | 0.34697996 | 0.34599411 | 946.697203 | 1302.61599 | 5.05263158 | 10.8679245 |
| 24 / UG / ROI: | 0.5170494 | 0.33127043 | 831.984068 | 1010.6462 | 1.26315789 | 2.03773585 |
| 26 / UG / ROI: | 0.44839891 | 0.5763872 | 783.501446 | 886.670154 | 18.9473684 | 19.245283 |
| 60 / UG / ROI: | 0.30970401 | 0.40184428 | 1008.51208 | 1534.63955 | 6.31578947 | 2.26415094 |
| 50 / UG / ROI: | 0.39443347 | 0.64185723 | 970.577731 | 1005.96406 | 1.89473684 | 0.90566038 |
| 27 / UG / ROI: | 0.46070375 | 0.55250431 | 1130.5297 | 880.44618 | 13.8947368 | 19.9245283 |
| 64 / UG / ROI: | 0.37055487 | 0.42686038 | 1121.21874 | 965.068481 | 16.4210526 | 19.245283 |
| 22 / UG / ROI: | 0.57531906 | 0.38211058 | 885.671554 | 1081.24579 | 9.47368421 | 9.50943396 |
| 41 / UG / ROI: | 0.25530432 | 0.37952326 | 1104.98613 | 1308.48689 | 1.26315789 | 0.67924528 |
| 9 / UG / ROI:{ | 0.36625038 | 0.42342444 | 1202.63433 | 1048.04531 | 11.3684211 | 12.2264151 |
| 25 / UG / ROI: | 0.07439994 | 1.43244302 | 1002.19528 | 1854.14582 | 0.10325648 | 0.22641509 |
| 61 / UG / ROI: | 0.28127726 | 0.40681437 | 1069.90832 | 966.901454 | 0.63157895 | 2.49056604 |
| 10 / UG / ROI: | 0.07299377 | 0.48786092 | 1337.3854 | 835.887271 | 2.13154584 | 4 |
| 16 / UG / ROI: | 0.40569487 | 0.44406513 | 847.636874 | 1006.83537 | 1.33333333 | 2.22222222 |
| 47 / UG / ROI: | 0.45109733 | 0.52597453 | 1084.45273 | 962.29514 | 7.33333333 | 6.66666667 |
| 60 / UG / ROI: | 0.07293994 | 0.37176887 | 1355.56591 | 4146.6217 | 0.11232111 | 0.22222222 |
| 35 / UG / ROI: | 0.07439944 | 0.40700128 | 1129.63826 | 3473.63391 | 0.11232111 | 0.22222222 |
| 49 / UG / ROI: | 0.09399423 | 0.26140679 | 941.365217 | 4395.2817 | 0.11232111 | 0.22222222 |
| 41 / UG / ROI: | 0.26123364 | 0.85961544 | 2948.9338 | 3812.56824 | 0.66666667 | 0.66666667 |
| 26 / UG / ROI: | 0.05556123 | 0.32874028 | 2457.44483 | 1410.88826 | 0.33112468 | 0.66666667 |
| 38 / UG / ROI: | 0.07439942 | 0.25562906 | 2047.8707 | 4606.28786 | 0.13454644 | 0.22222222 |
| 61 / UG / ROI: | 0.07284399 | 0.27788227 | 1706.55891 | 1375.16822 | 0.21131546 | 0.44444444 |

**Fig. 2C (b)** standard aCSF vs oscill. aCSF

Data Source File

| | Amplitude (df/f) | Amplitude (df/f) | Duration (ms) | Duration (ms) | Frequency (Events Nb/min) | Frequency |
|---|---|---|---|---|---|---|
| 34 / UG / ROI: | 0.35441147 | 0.37011567 | 868.248414 | 1158.59475 | 1.33333333 | 2 |
| 54 / UG / ROI: | 0.33415382 | 0.39473875 | 1140.34369 | 1143.26016 | 10 | 11.3333333 |
| 25 / UG / ROI: | 0.31701958 | 0.34848916 | 1251.59409 | 1075.30505 | 6.66666667 | 10.6666667 |
| 3 / UG / ROI:{a | 0.08894494 | 1.24735947 | 1042.99507 | 3107.88689 | 0.3315466 | 0.66666667 |
| 64 / UG / ROI: | 0.33267122 | 0.33914093 | 1091.81854 | 1311.13087 | 2.66666667 | 4.66666667 |
| 36 / UG / ROI: | 0.07449393 | 0.37553753 | 909.848785 | 1450.52539 | 0.2132156 | 0.44444444 |
| 15 / UG / ROI: | 0.52655939 | 0.58681664 | 804.182653 | 939.240015 | 1 | 1.11111111 |
| 11 / UG / ROI: | 0.32063891 | 0.55982306 | 1670.48974 | 759.786445 | 2.66666667 | 3.33333333 |
| 1 / UG / ROI:{8 | 0.46943792 | 0.57545806 | 1477.83934 | 1279.8398 | 4.66666667 | 4.88888889 |
| 4 / UG / ROI:{6 | 0.45818196 | 0.55578972 | 1503.73446 | 1893.07738 | 2 | 2.44444444 |
| 28 / UG / ROI: | 0.29222648 | 0.38883588 | 1385.22927 | 1574.27154 | 4 | 2.66666667 |
| 8 / UG / ROI:{5 | 0.3377066 | 0.44202155 | 751.507774 | 990.339598 | 3.33333333 | 9.11111111 |
| 45 / UG / ROI: | 0.42126274 | 0.46586999 | 1263.01274 | 1345.62322 | 10 | 10.6666667 |
| 21 / UG / ROI: | 0.38765889 | 0.62485369 | 1166.71371 | 1182.40942 | 7.33333333 | 7.33333333 |
| 17 / UG / ROI: | 0.34440248 | 0.39190267 | 1212.76092 | 1270.42547 | 6.66666667 | 6.66666667 |
| 62 / UG / ROI: | 0.07274494 | 0.31334418 | 1010.6341 | 1057.58261 | 0.22311333 | 0.44444444 |
| 31 / UG / ROI: | 0.317438 | 0.43986723 | 2106.73798 | 2463.47452 | 0.66666667 | 0.66666667 |
| 12 / UG / ROI: | 0.37749063 | 0.38817588 | 1095.55919 | 1095.97368 | 3.33333333 | 3.77777778 |
| 40 / UG / ROI: | 0.39239268 | 0.37951275 | 1104.53172 | 1200.68797 | 7.33333333 | 8.44444444 |
| 68 / UG / ROI: | 0.43467191 | 0.43449571 | 928.144723 | 1681.26959 | 6.66666667 | 3.77777778 |
| 24 / UG / ROI: | 0.59472444 | 0.61145174 | 940.402306 | 1015.6105 | 18.6666667 | 21.7777778 |
| 66 / UG / ROI: | 0.45439541 | 0.35933432 | 1050.42466 | 1025.45666 | 13.3333333 | 15.3333333 |
| 44 / UG / ROI: | 0.35206057 | 0.39843372 | 1069.85236 | 1052.55076 | 7.33333333 | 10 |
| 2 / UG / ROI:{1 | 0.27586303 | 0.65336337 | 977.824884 | 1219.05424 | 2 | 3.33333333 |
| 29 / UG / ROI: | 0.35748723 | 0.39050221 | 1443.83654 | 1427.06531 | 6.66666667 | 8.22222222 |
| 56 / UG / ROI: | 0.37907996 | 0.55368613 | 1034.21769 | 1042.74282 | 5.33333333 | 4.88888889 |
| 52 / UG / ROI: | 0.35489 | 0.34737971 | 1329.0544 | 1152.09601 | 9.33333333 | 10.4444444 |
| 46 / UG / ROI: | 0.48735185 | 0.92180911 | 962.093588 | 1279.91722 | 20 | 20.6666667 |
| 9 / UG / ROI:{7 | 0.3449322 | 0.54362841 | 994.076437 | 876.099503 | 14 | 16.6666667 |
| 59 / UG / ROI: | 0.07224388 | 0.35695007 | 828.397031 | 3487.01478 | 0.1131215 | 0.22222222 |
| 30 / UG / ROI: | 0.45195777 | 0.49665649 | 1042.24885 | 1497.16741 | 2.66666667 | 3.33333333 |
| 14 / UG / ROI: | 0.67200371 | 0.9737234 | 942.832254 | 1142.10514 | 8.66666667 | 6 |
| 58 / UG / ROI: | 0.34181607 | 0.36353626 | 916.571282 | 1246.66632 | 4.66666667 | 3.55555556 |
| 7 / UG / ROI:{4 | 0.56894777 | 0.95926298 | 2326.07404 | 2479.64646 | 3.33333333 | 4.88888889 |
| 19 / UG / ROI: | 0.43285759 | 0.5365741 | 948.505919 | 916.826871 | 6.66666667 | 4.66666667 |
| 57 / UG / ROI: | 0.31059399 | 0.36123581 | 969.648702 | 1099.84593 | 13.3333333 | 10.4444444 |
| 55 / UG / ROI: | 0.28163918 | 0.30108655 | 1288.39929 | 1417.91664 | 2 | 2.22222222 |
| 65 / UG / ROI: | 0.35975699 | 0.38566493 | 757.964367 | 926.357812 | 2.66666667 | 4.44444444 |
| MEAN | 0.318 | 0.4887 | 1214 | 1431 | 4.78 | 5.461 |
| SEM | 0.01677 | 0.02639 | 45.69 | 87.96 | 0.5568 | 0.6303 |

**Wilcoxon matched-pairs signed rank test** standard aCSF vs oscill. aCSF

| Astro. transients | Amplitude (df/f) | Duration (ms) | Frequency (Events Nb/min) |
|---|---|---|---|
| P value | <0.0001 | 0.0097 | 0.0001 |

**Fig. 2C (c)** Data Source File

| | TTX Amplitude (df/f) | TTX+Cocktail Amplitude (df/f) | TTX Duration (ms) | TTX+Cocktail Duration (ms) | TTXX Frequency (Events Nb/min) | TTX+Cocktail Frequency (Events Nb/min) |
|---|---|---|---|---|---|---|
| 43 / UG / ROI: | 0.33195078 | 0.44112282 | 1361.81633 | 1781.11189 | 2.52631579 | 3.72413793 |
| 44 / UG / ROI: | 0.49245571 | 0.41354446 | 892.997169 | 1227.41817 | 4.42105263 | 6.62068966 |
| 45 / UG / ROI: | 0.8921256 | 0.50284012 | 1143.14121 | 997.868228 | 5.05263158 | 4.55172414 |
| 46 / UG / ROI: | 0.34835155 | 0.38528172 | 1112.38018 | 1028.45218 | 6.94736842 | 5.37931034 |
| 47 / UG / ROI: | 0.46359073 | 0.54131679 | 1057.86477 | 1559.99689 | 9.47368421 | 4.55172414 |
| 49 / UG / ROI: | 0.31984262 | 0.34127549 | 884.651504 | 1278.26561 | 10.7368421 | 10.3448276 |
| 50 / UG / ROI: | 0.28251339 | 0.33368444 | 932.537404 | 973.647251 | 13.2631579 | 12.4137931 |
| 51 / UG / ROI: | 0.28512765 | 0.36930663 | 1499.03405 | 1365.16004 | 6.31578947 | 4.96551724 |
| 52 / UG / ROI: | 0.31406982 | 0.33043513 | 1161.77362 | 1948.19488 | 4.42105263 | 3.31034483 |
| 53 / UG / ROI: | 0.68194323 | 0.48556817 | 4201.40188 | 1603.96981 | 3.78947368 | 2.48275862 |
| 54 / UG / ROI: | 0.29624728 | 0.33656189 | 3820.04128 | 2048.06484 | 0.63157895 | 0.82758621 |
| 60 / UG / ROI: | 0.35491432 | 0.40390547 | 891.311408 | 1087.99518 | 15.7894737 | 14.8965517 |
| 61 / UG / ROI: | 0.28655976 | 0.31349468 | 1129.86702 | 1550.69565 | 4.42105263 | 5.37931034 |
| 64 / UG / ROI: | 0.27357431 | 0.36241305 | 1260.66979 | 1367.00166 | 3.78947368 | 8.68965517 |
| 65 / UG / ROI: | 0.30039333 | 0.44669145 | 1322.4048 | 1103.16163 | 4.42105263 | 3.31034483 |
| 67 / UG / ROI: | 0.25760414 | 0.34602178 | 1170.62855 | 1189.71107 | 4.42105263 | 8.27586207 |
| 69 / UG / ROI: | 0.25966038 | 0.34212029 | 1041.04278 | 1268.36837 | 0.63157895 | 2.48275862 |
| 72 / UG / ROI: | 0.26565728 | 0.31694742 | 1066.45626 | 1240.17019 | 2.52631579 | 5.37931034 |
| 77 / UG / ROI: | 0.45059155 | 0.45759703 | 915.082157 | 1170.41912 | 11.3684211 | 12.8275862 |
| 78 / UG / ROI: | 0.29889034 | 0.35618373 | 1292.25838 | 1261.22276 | 7.57894737 | 8.27586207 |
| 80 / UG / ROI: | 0.53680934 | 0.43736278 | 1090.15977 | 956.645634 | 15.7894737 | 16.137931 |
| 22 / UG / ROI: | 0.52976258 | 0.59291904 | 1393.86074 | 1213.44705 | 5.88235294 | 6.95652174 |
| 23 / UG / ROI: | 1.25240862 | 0.76927128 | 1066.20025 | 1036.91098 | 10.5882353 | 10.8695652 |
| 24 / UG / ROI: | 0.51110727 | 0.9769758 | 1361.68389 | 1055.23182 | 6.47058824 | 6.52173913 |
| 25 / UG / ROI: | 0.48396182 | 0.61511475 | 1591.82424 | 1213.13268 | 5.29411765 | 8.26086957 |
| 29 / UG / ROI: | 0.48333494 | 0.63787856 | 1461.68981 | 1246.48134 | 4.11764706 | 3.47826087 |
| 35 / UG / ROI: | 1.0187258 | 0.83650269 | 767.574006 | 821.367055 | 31.7647059 | 28.6956522 |
| 37 / UG / ROI: | 0.57793191 | 0.84099646 | 1202.11406 | 1028.42501 | 11.7647059 | 11.7391304 |
| 55 / UG / ROI: | 0.3110748 | 0.35144677 | 1691.66853 | 1275.69719 | 3.78947368 | 7.44827586 |
| 56 / UG / ROI: | 0.28900913 | 0.40604185 | 783.837056 | 1024.038 | 0.63157895 | 2.48275862 |
| 58 / UG / ROI: | 0.24019796 | 0.29115117 | 2231.45019 | 1244.90508 | 1.26315789 | 4.55172414 |
| 62 / UG / ROI: | 0.06177278 | 0.4101451 | 1239.69455 | 1467.92758 | 2.02155564 | 4.13793103 |
| 63 / UG / ROI: | 0.06627329 | 0.32032301 | 1033.07879 | 2013.14429 | 1.21234555 | 2.48275862 |
| 68 / UG / ROI: | 0.06327449 | 0.4704343 | 860.898992 | 981.006675 | 1.02315556 | 1.24137931 |
| 71 / UG / ROI: | 0.07328947 | 0.39253892 | 717.415827 | 937.625914 | 1.23545888 | 2.06896552 |
| 73 / UG / ROI: | 0.08427324 | 0.32281645 | 597.846522 | 946.925016 | 1.02123333 | 2.06896552 |
| 74 / UG / ROI: | 0.3815393 | 0.55941821 | 1435.81397 | 1275.32291 | 3.15789474 | 4.55172414 |
| 76 / UG / ROI: | 0.27516943 | 0.3529703 | 923.330501 | 1180.50307 | 1.26315789 | 5.37931034 |
| 26 / UG / ROI: | 0.62107306 | 0.67972084 | 1045.2191 | 952.611945 | 12.3529412 | 15.6521739 |
| 27 / UG / ROI: | 0.52237231 | 0.5295141 | 974.534371 | 1056.87622 | 8.82352941 | 11.7391304 |
| 31 / UG / ROI: | 0.44028652 | 0.55162941 | 1179.0521 | 908.050797 | 0.58823529 | 2.17391304 |
| 33 / UG / ROI: | 0.57711195 | 0.65915124 | 1372.76558 | 1437.42422 | 7.05882353 | 6.95652174 |
| 34 / UG / ROI: | 0.06628229 | 1.13131117 | 980.546845 | 1473.73963 | 0.21315488 | 0.43478261 |
| 50 / UG / ROI: | 0.92741843 | 0.90615343 | 2066.60842 | 1423.02484 | 2.94117647 | 1.12149533 |
| 52 / UG / ROI: | 1.19082683 | 0.81420955 | 1170.67793 | 1431.34957 | 3.52941176 | 2.80373832 |
| 55 / UG / ROI: | 0.83052665 | 1.17512662 | 1205.91279 | 1217.02447 | 2.94117647 | 2.52336449 |
| 56 / UG / ROI: | 0.89713027 | 0.95073291 | 1188.99969 | 1127.20228 | 7.05882353 | 4.76635514 |

**Fig. 2C (d)**

Data Source File

| | TTX Amplitude (df/f) | TTX+Cocktail Amplitude (df/f) | TTX Duration (ms) | TTX+Cocktail Duration (ms) | TTXX Frequency (Events Nb/min) | TTX+Cocktail Frequency (Events Nb/min) |
|---|---|---|---|---|---|---|
| 57 / UG / ROI: | 0.06627778 | 0.78627116 | 849.285493 | 944.584315 | 1.21354888 | 3.36448598 |
| 59 / UG / ROI: | 0.66905372 | 0.78482951 | 1081.1669 | 1059.20677 | 12.3529412 | 9.53271028 |
| 62 / UG / ROI: | 0.68115007 | 0.89064148 | 1211.24105 | 1324.16863 | 6.47058824 | 5.60747664 |
| 63 / UG / ROI: | 0.08329445 | 1.5601863 | 757.025657 | 803.105332 | 0.78987888 | 1.40186916 |
| 64 / UG / ROI: | 0.68294158 | 0.76337118 | 1030.35039 | 1052.01343 | 11.1764706 | 9.25233645 |
| 68 / UG / ROI: | 0.9481923 | 0.95735514 | 922.658136 | 990.472897 | 17.0588235 | 14.8598131 |
| 69 / UG / ROI: | 0.91530188 | 1.1922164 | 1122.71187 | 1388.67388 | 9.41176471 | 5.60747664 |
| 71 / UG / ROI: | 0.75749144 | 0.77544501 | 1148.13587 | 1088.70356 | 14.7058824 | 10.0934579 |
| 76 / UG / ROI: | 0.77962686 | 0.73752206 | 1132.59696 | 1266.42943 | 10.5882353 | 8.97196262 |
| 81 / UG / ROI: | 0.90472685 | 0.96466479 | 869.823411 | 961.58819 | 22.3529412 | 19.3457944 |
| 85 / UG / ROI: | 0.71830049 | 0.75492852 | 926.094776 | 1209.42114 | 10.5882353 | 11.7757009 |
| 89 / UG / ROI: | 0.80044463 | 0.89470693 | 1010.47335 | 1032.28296 | 15.8823529 | 8.97196262 |
| 90 / UG / ROI: | 0.75198861 | 0.75870607 | 967.532126 | 970.774239 | 16.4705882 | 13.4579439 |
| 93 / UG / ROI: | 0.96448827 | 0.9228517 | 1185.32346 | 1134.11927 | 4.11764706 | 4.48598131 |
| 95 / UG / ROI: | 0.80351025 | 0.68435811 | 1654.63998 | 2012.21678 | 1.17647059 | 1.12149533 |
| 96 / UG / ROI: | 0.61049728 | 0.86585168 | 1449.19492 | 1275.92483 | 5.88235294 | 3.92523364 |
| 31 / UG / ROI: | 1.02742258 | 1.03081948 | 967.317352 | 941.929888 | 21.7647059 | 23.5514019 |
| 65 / UG / ROI: | 1.3032919 | 0.83935712 | 969.881612 | 1162.63353 | 8.23529412 | 8.69158879 |
| 66 / UG / ROI: | 0.94985966 | 1.01840868 | 816.141315 | 968.985489 | 21.1764706 | 19.6261682 |
| 72 / UG / ROI: | 0.60862141 | 1.08199888 | 1142.40423 | 1221.08192 | 2.35294118 | 2.52336449 |
| 73 / UG / ROI: | 0.8546686 | 0.84267212 | 943.222194 | 1093.7329 | 2.94117647 | 6.1682243 |
| 74 / UG / ROI: | 0.96144565 | 0.91034203 | 1421.66051 | 1160.34056 | 8.23529412 | 11.2149533 |
| 77 / UG / ROI: | 1.26435219 | 1.5125615 | 683.977504 | 798.099918 | 38.8235294 | 35.3271028 |
| 78 / UG / ROI: | 0.06610617 | 1.39178654 | 854.97188 | 1173.57709 | 0.12322222 | 0.28037383 |
| 79 / UG / ROI: | 0.09232707 | 1.55921878 | 712.476567 | 844.350706 | 0.12322222 | 0.28037383 |
| 80 / UG / ROI: | 0.67343103 | 1.02538887 | 1091.30868 | 1171.49781 | 2.35294118 | 2.80373832 |
| 82 / UG / ROI: | 0.7560219 | 1.22586411 | 1365.7873 | 1383.68006 | 1.17647059 | 0.8411215 |
| 83 / UG / ROI: | 0.81625807 | 0.86988761 | 844.323852 | 902.558426 | 22.9411765 | 18.5046729 |
| 84 / UG / ROI: | 0.86165457 | 0.95548484 | 900.814505 | 862.392377 | 24.1176471 | 19.6261682 |
| 94 / UG / ROI: | 0.54993496 | 1.066791 | 1121.93544 | 1012.89352 | 1.17647059 | 4.20560748 |
| 97 / UG / ROI: | 0.06606078 | 1.53249996 | 934.946203 | 1421.73292 | 0.12322222 | 0.28037383 |
| 21 / UG / ROI: | 0.75234785 | 1.31910862 | 1031.49779 | 1969.87171 | 1.76470588 | 0.8411215 |
| 22 / UG / ROI: | 0.06777944 | 0.683121 | 859.581488 | 936.372064 | 0.12322222 | 0.28037383 |
| 23 / UG / ROI: | 0.73330353 | 0.73738612 | 1394.83859 | 1342.26242 | 2.94117647 | 3.64485981 |
| 24 / UG / ROI: | 0.06627822 | 0.5314244 | 871.774118 | 769.05205 | 0.12322222 | 0.28037383 |
| 80 / UG / ROI: | 0.39260421 | 0.4502487 | 873.224674 | 1095.10023 | 18.5185185 | 15.483871 |
| 32 / UG / ROI: | 0.05662783 | 0.3864912 | 623.73191 | 1191.4486 | 1.11223254 | 2.15053763 |
| 48 / UG / ROI: | 0.31737577 | 0.34269523 | 1138.75715 | 1352.22791 | 4.44444444 | 5.16129032 |
| 101 / UG / RO | 0.3249023 | 0.33136182 | 1167.69666 | 1320.94394 | 9.62962963 | 7.95698925 |
| 63 / UG / ROI: | 0.2919169 | 0.40724567 | 1750.89673 | 1360.81674 | 2.22222222 | 1.29032258 |
| 73 / UG / ROI: | 0.05662822 | 0.52704759 | 1250.64052 | 3921.60944 | 0.21325564 | 0.43010753 |
| 50 / UG / ROI: | 0.09926667 | 0.38556855 | 893.314658 | 1184.61832 | 1.12123215 | 2.79569892 |
| 42 / UG / ROI: | 0.32730501 | 0.41075678 | 1108.41368 | 1202.9966 | 13.3333333 | 10.5376344 |
| 81 / UG / ROI: | 0.38533803 | 0.44127235 | 1079.69139 | 1297.82907 | 10.3703704 | 12.4731183 |
| 86 / UG / ROI: | 0.38922012 | 0.4192227 | 1056.29863 | 1156.81598 | 5.92592593 | 6.88172043 |
| 12 / UG / ROI: | 0.26119389 | 0.49487327 | 1140.45204 | 1576.34696 | 2.22222222 | 1.50537634 |
| 56 / UG / ROI: | 0.45730251 | 0.38935894 | 4296.34718 | 938.650036 | 0.74074074 | 1.50537634 |
| 49 / UG / ROI: | 0.37634535 | 0.69900976 | 1077.9628 | 889.118952 | 1.48148148 | 5.16129032 |
| 57 / UG / ROI: | 0.06732439 | 0.49784475 | 769.973426 | 886.871676 | 0.78988845 | 1.50537634 |
| 4 / UG / ROI:{( | 0.34144445 | 0.54177155 | 1172.34451 | 1500.69368 | 9.62962963 | 2.79569892 |

| Fig. 2C (e) | TTX Amplitude (df/f) | TTX+Cocktail Amplitude (df/f) | TTX Duration (ms) | TTX+Cocktail Duration (ms) | TTXX Frequency (Events Nb/min) | TTX+Cocktail Frequency (Events Nb/min) | Data Source File |
|---|---|---|---|---|---|---|---|
| 69 / UG / ROI: | 0.28975838 | 0.47927299 | 1983.09229 | 892.175683 | 3.7037037 | 5.59139785 | |
| 72 / UG / ROI: | 0.29862736 | 0.40622063 | 881.732514 | 1287.44371 | 2.96296296 | 3.65591398 | |
| 21 / UG / ROI: | 0.33778628 | 0.47743494 | 1236.64339 | 1131.83995 | 8.88888889 | 13.1182796 | |
| 71 / UG / ROI: | 0.33531735 | 0.40472388 | 1196.77169 | 1295.76603 | 7.40740741 | 6.66666667 | |
| 19 / UG / ROI: | 0.06723278 | 0.43646927 | 854.836921 | 1002.29919 | 0.45658799 | 1.50537634 | |
| 44 / UG / ROI: | 0.39764613 | 0.54785484 | 929.828539 | 967.78741 | 18.5185185 | 22.5806452 | |
| 52 / UG / ROI: | 0.07329277 | 0.38325787 | 664.163242 | 1412.15438 | 0.52213564 | 1.07526882 | |
| 46 / UG / ROI: | 0.27480909 | 0.5705107 | 795.268183 | 1014.69276 | 0.74074074 | 6.88172043 | |
| 64 / UG / ROI: | 0.45152666 | 0.4526828 | 895.870335 | 1015.66971 | 21.4814815 | 17.4193548 | |
| 15 / UG / ROI: | 0.25635802 | 0.68267071 | 813.280298 | 948.80174 | 0.74074074 | 4.08602151 | |
| 95 / UG / ROI: | 0.34554486 | 0.3865769 | 1328.72859 | 1296.79844 | 5.18518519 | 6.23655914 | |
| 87 / UG / ROI: | 0.31057408 | 0.4238685 | 1008.07981 | 1112.7032 | 7.40740741 | 6.66666667 | |
| 83 / UG / ROI: | 0.73904696 | 0.47429575 | 1325.12021 | 983.547563 | 1.48148148 | 4.08602151 | |
| 30 / UG / ROI: | 0.263644 | 0.38089303 | 1458.59912 | 1062.08243 | 2.22222222 | 4.7311828 | |
| 18 / UG / ROI: | 0.31953126 | 0.42048096 | 4511.47605 | 1392.33949 | 2.22222222 | 6.4516129 | |
| 23 / UG / ROI: | 0.44516121 | 0.36860978 | 1427.19145 | 1217.88032 | 6.66666667 | 7.09677419 | |
| 53 / UG / ROI: | 0.28619617 | 0.42846089 | 1443.44454 | 1374.71745 | 1.48148148 | 3.65591398 | |
| 54 / UG / ROI: | 0.49601205 | 0.35918121 | 4218.82916 | 1184.9887 | 0.74074074 | 1.50537634 | |
| 60 / UG / ROI: | 0.35904105 | 0.45931612 | 1246.30591 | 1089.31443 | 12.5925926 | 11.6129032 | |
| 84 / UG / ROI: | 0.31824437 | 0.42304342 | 1108.44193 | 1162.25328 | 4.44444444 | 6.88172043 | |
| 28 / UG / ROI: | 0.4484901 | 0.38322947 | 1386.18135 | 1407.99386 | 5.18518519 | 5.37634409 | |
| 9 / UG / ROI:{( | 0.35549373 | 0.44549306 | 1436.47919 | 1292.71053 | 2.96296296 | 2.3655914 | |
| 3 / UG / ROI:{( | 0.32954332 | 0.52972442 | 1526.4242 | 1988.03226 | 1.48148148 | 1.72043011 | |
| 25 / UG / ROI: | 0.31379218 | 0.36406233 | 1835.36043 | 1653.656 | 0.74074074 | 1.50537634 | |
| 14 / UG / ROI: | 0.34376896 | 0.37220274 | 766.1965 | 1153.68091 | 1.48148148 | 4.08602151 | |
| 29 / UG / ROI: | 0.31292404 | 0.42208626 | 1090.2582 | 1241.08807 | 5.92592593 | 6.66666667 | |
| 24 / UG / ROI: | 0.35491169 | 0.35436119 | 870.636594 | 1434.30589 | 0.74074074 | 1.72043011 | |
| 43 / UG / ROI: | 0.32794825 | 0.50633578 | 1332.19494 | 970.573056 | 8.88888889 | 9.46236559 | |
| 67 / UG / ROI: | 0.39066172 | 0.3826881 | 901.817678 | 1232.28284 | 2.96296296 | 7.52688172 | |
| 51 / UG / ROI: | 0.26689147 | 0.39235218 | 1021.79212 | 1224.54363 | 2.22222222 | 1.29032258 | |
| 55 / UG / ROI: | 0.54056167 | 0.52097676 | 813.730417 | 958.731007 | 25.9259259 | 21.5053763 | |
| 5 / UG / ROI:{5 | 0.38690053 | 0.72248127 | 1560.29684 | 1605.86769 | 4.44444444 | 3.01075269 | |
| 88 / UG / ROI: | 0.32779374 | 0.38004601 | 1281.58055 | 972.290656 | 8.88888889 | 11.6129032 | |
| 74 / UG / ROI: | 0.32423605 | 0.50370481 | 1342.74077 | 1112.07916 | 3.7037037 | 9.67741935 | |
| 85 / UG / ROI: | 0.30718424 | 0.35983844 | 1297.19043 | 1206.63417 | 4.44444444 | 5.59139785 | |
| 13 / UG / ROI: | 0.53511689 | 0.37553868 | 2157.7685 | 1136.70987 | 3.7037037 | 5.16129032 | |
| 100 / UG / RO | 0.34323684 | 0.38192617 | 1036.1331 | 1095.36231 | 13.3333333 | 13.1182796 | |
| 92 / UG / ROI: | 0.45682843 | 0.41963155 | 959.862259 | 960.648422 | 17.7777778 | 20 | |
| 31 / UG / ROI: | 0.38312837 | 0.41816466 | 1008.3659 | 1087.39487 | 1.48148148 | 1.29032258 | |
| 59 / UG / ROI: | 0.32347077 | 0.36642678 | 1128.53793 | 1413.84774 | 5.18518519 | 6.88172043 | |
| 62 / UG / ROI: | 0.25396784 | 0.36997479 | 936.634037 | 1207.60501 | 0.74074074 | 4.08602151 | |
| 33 / UG / ROI: | 0.37500224 | 0.45370418 | 1325.13696 | 925.834281 | 6.66666667 | 7.95698925 | |
| 6 / UG / ROI:{( | 0.32024458 | 0.39186394 | 1127.69421 | 1045.36329 | 3.7037037 | 2.15053763 | |
| 97 / UG / ROI: | 0.43040415 | 0.50735469 | 950.351575 | 1085.31556 | 22.2222222 | 22.5806452 | |
| 16 / UG / ROI: | 0.26443078 | 0.43135832 | 793.33462 | 1494.13655 | 1.48148148 | 6.4516129 | |
| 17 / UG / ROI: | 0.32676446 | 0.52613435 | 1242.25153 | 1263.78755 | 8.14814815 | 13.9784946 | |
| 61 / UG / ROI: | 0.32753986 | 0.31526956 | 810.636192 | 1217.19583 | 0.74074074 | 1.29032258 | |
| 37 / UG / ROI: | 0.33468432 | 0.54560383 | 892.648185 | 2009.79062 | 0.74074074 | 2.3655914 | |
| 79 / UG / ROI: | 0.35897396 | 0.4702292 | 1110.69969 | 1165.42276 | 13.3333333 | 15.9139785 | |
| 105 / UG / RO | 0.29274745 | 0.37966559 | 975.727849 | 1095.36048 | 2.96296296 | 3.65591398 | |

**Fig. 2C (f)**

| | TTX Amplitude (df/f) | TTX+Cocktail Amplitude (df/f) | TTX Duration (ms) | TTX+Cocktail Duration (ms) | TTXX Frequency (Events Nb/min) | TTX+Cocktail Frequency (Events Nb/min) | Data Source File |
|---|---|---|---|---|---|---|---|
| 106 / UG / RO | 0.44547608 | 0.38765471 | 1001.09642 | 992.746564 | 19.2592593 | 20 | |
| 36 / UG / ROI: | 0.36277321 | 0.53371978 | 1006.0078 | 1092.65659 | 5.92592593 | 3.65591398 | |
| 68 / UG / ROI: | 0.3234275 | 0.43476436 | 1180.87033 | 1357.55118 | 11.8518519 | 8.60215054 | |
| 45 / UG / ROI: | 0.33070527 | 0.55551553 | 1249.56375 | 1951.11607 | 5.18518519 | 3.01075269 | |
| 78 / UG / ROI: | 0.33366118 | 0.40784231 | 1071.91742 | 1130.63897 | 9.62962963 | 5.80645161 | |
| 70 / UG / ROI: | 0.06622283 | 0.38869302 | 1339.89678 | 1714.1634 | 0.12324556 | 1.93548387 | |
| 40 / UG / ROI: | 0.07944978 | 0.63563995 | 1116.58065 | 2340.07055 | 0.23156545 | 0.43010753 | |
| 41 / UG / ROI: | 0.30275294 | 0.36015511 | 941.627131 | 1472.05352 | 9.62962963 | 5.59139785 | |
| 77 / UG / ROI: | 0.37466971 | 0.3789662 | 812.594778 | 1307.67548 | 2.96296296 | 3.65591398 | |
| 65 / UG / ROI: | 0.06111556 | 0.69452176 | 677.162315 | 2411.01116 | 0.31564898 | 0.64516129 | |
| 39 / UG / ROI: | 0.39398152 | 0.42375885 | 1264.66347 | 1346.03993 | 7.40740741 | 4.30107527 | |
| 75 / UG / ROI: | 0.26886966 | 0.37674835 | 1836.80758 | 2274.87289 | 1.48148148 | 1.07526882 | |
| 76 / UG / ROI: | 0.38641039 | 0.40525619 | 1467.98762 | 1188.91699 | 8.14814815 | 6.23655914 | |
| 35 / UG / ROI: | 0.32123103 | 0.60812318 | 1385.42072 | 1757.83599 | 2.22222222 | 2.15053763 | |
| 82 / UG / ROI: | 0.31212492 | 0.34025225 | 1354.58029 | 1112.13521 | 4.44444444 | 9.46236559 | |
| 94 / UG / ROI: | 0.39260327 | 0.32192244 | 1266.49257 | 1223.42357 | 8.88888889 | 4.51612903 | |
| 58 / UG / ROI: | 0.36785572 | 0.43801024 | 1019.10761 | 1037.87347 | 10.3703704 | 12.9032258 | |
| 66 / UG / ROI: | 0.28994148 | 0.50833321 | 1204.11086 | 1195.227 | 4.44444444 | 6.88172043 | |
| MEAN | 0.4319 | 0.5825 | 1224 | 1254 | 6.567 | 6.889 | |
| SEM | 0.02098 | 0.02188 | 46.95 | 28.71 | 0.527 | 0.4729 | |

**Wilcoxon matched-pairs signed rank test** TTX vs TTX+Cocktail

| Astro. transients | Amplitude (df/f) | Duration (ms) | Frequency (Events Nb/min) |
|---|---|---|---|
| P value | <0.0001 | 0.0009 | 0.032 |

**Fig. 2D** Data Source File

| Oscill. aCSF | mouse#1 P16 | mouse#2 P16 | TOTAL |
|---|---|---|---|
| | ROIs nb | ROIs nb | ROIs nb |
| Preceding Neuronal Oscill. | 20 | 35 | 55 |
| During Neuronal Oscill. | 18 | 13 | 31 |

| Oscill. aCSF | mouse#1 P16 | mouse#2 P16 | MEAN |
|---|---|---|---|
| | % | % | % |
| Preceding Neuronal Oscill. | 52.6315789 | 72.9166667 | 63.9534884 |
| During Neuronal Oscill. | 47.3684211 | 27.0833333 | 36.0465116 |

| TTX+Cocktail | mouse#1 P16 | mouse#2 P16 | mouse#3 P13 | TOTAL |
|---|---|---|---|---|
| | ROIs nb | ROIs nb | ROIs nb | ROIs nb |
| Preceding Neuronal Oscill. | 28 | 21 | 62 | 111 |
| During Neuronal Oscill. | 15 | 18 | 21 | 54 |

| TTX+Cocktail | mouse#1 P16 | mouse#2 P16 | mouse#3 P13 | MEAN |
|---|---|---|---|---|
| | % | % | % | % |
| Preceding Neuronal Oscill. | 65.1162791 | 53.8461538 | 74.6987952 | 64.5537427 |
| During Neuronal Oscill. | 34.8837209 | 46.1538462 | 25.3012048 | 35.4462573 |

**Fig. 3B** Data Source File

| | Age | | Vrest Astro. (mV) | Rin Astro. (Mohm) |
|---|---|---|---|---|
| mouse#1 | P11 | astro 1 | -85.1341797 | 8.53514862 |
| mouse#1 | P11 | astro 2 | -85.4622753 | 6.49588903 |
| mouse#2 | P12 | astro 3 | -82.7101561 | 19.4316016 |
| mouse#2 | P12 | astro 4 | -90.206466 | 12.0715035 |
| mouse#3 | P9 | astro 5 | -87.1432915 | 75.0576436 |
| mouse#3 | P9 | astro 6 | -83.5922447 | 19.0843565 |
| mouse#3 | P9 | astro 7 | -80.2843338 | 22.1349349 |
| mouse#3 | P9 | astro 8 | -89.0165222 | 30.1026521 |
| mouse#4 | P4 | astro 9 | -89.4133862 | 34.7171885 |
| mouse#4 | P4 | astro 10 | -96.7491119 | 3.70445954 |
| mouse#4 | P4 | astro 11 | -79.6710857 | 29.1137356 |
| mouse#5 | P13 | astro 12 | -87.2568771 | 35.0382784 |
| mouse#5 | P13 | astro 13 | -90.2398026 | 20.850366 |
| mouse#5 | P13 | astro 14 | -85.113826 | 17.1491475 |
| mouse#6 | P7 | astro 15 | -96.4 | 17 |
| mouse#6 | P7 | astro 16 | -88.4 | 19.6 |
| mouse#7 | P5 | astro 17 | -84.4 | 51.8 |
| mouse#8 | P6 | astro 18 | -78.4 | 91.6 |
| mouse#9 | P9 | astro 19 | -86.4 | 41.61 |
| mouse#10 | P7 | astro 20 | -94.2 | 17 |
| mouse#11 | P8 | astro 21 | -92.4 | 9.8 |
| mouse#12 | P6 | astro 22 | -95 | 36 |
| mouse#12 | P6 | astro 23 | -93.6 | 20 |
| mouse#12 | P6 | astro 24 | -86.1 | 26 |
| mouse#13 | P7 | astro 25 | -87.7 | 26 |
| mouse#14 | P9 | astro 26 | -92.7 | 25 |
| mouse#14 | P9 | astro 27 | -90.3 | 17 |
| mouse#15 | P12 | astro 28 | -93.3 | 25.8 |
| | | MEAN | -88.26048424 | 27.0606038 |
| | | SEM | 0.937092427 | 3.64473334 |

**Fig. 3D (a)** Data Source File

| | mouse#1 | mouse#2 | mouse#3 | mouse#4 | mouse#5 | mouse#6 |
|---|---|---|---|---|---|---|
| | P9 | P11 | P8 | P6 | P4 | P12 |
| Voltage (mV) | Ba2+-sensitive | Ba2+-sensitive | Ba2+-sensitive | Ba2+-sensitive | Ba2+-sensitive | Ba2+-sensitive current (pA) |
| -140 | -660.516632 | -2584.0221 | -1087.31079 | -374.7 | -586.2 | -675.1 |
| -130 | | -2381.84875 | | -228.3 | -435.4 | -552.9760742 |
| -120 | -374.610535 | -2098.43591 | -335.144043 | -134 | -317.3 | -553.9269409 |
| -110 | | -1555.43079 | | -9.2 | -222.2 | -370.5203247 |
| -100 | -105.416428 | -1076.5014 | -96.7102051 | -39.1 | -68.4 | -131.619812 |
| -90 | | -509.460297 | | 57 | 48.2 | 25.2389679 |
| -80 | 92.3374691 | 143.952269 | 69.6411133 | 190 | 98 | 151.6219902 |
| -70 | | 447.822296 | | 797.7 | 230.7 | 297.0698242 |
| -60 | 330.345528 | 938.597839 | 197.113037 | 749.6 | 333.2 | 433.9328613 |
| -50 | | 1089.23589 | | 916.1 | 423 | 597.4717407 |
| -40 | 519.757935 | 1720.56976 | 306.915283 | 1013.9 | 772.2 | 884.58 |

| | mouse#7 | mouse#8 | mouse#8 | mouse#8 | mouse#8 | mouse#8 |
|---|---|---|---|---|---|---|
| | P5 | P7 | P7 | P7 | P7 | P7 |
| Voltage (mV) | Ba2+-sensitive | Ba2+-sensitive | Ba2+-sensitive | Ba2+-sensitive | Ba2+-sensitive | Ba2+-sensitive current (pA) |
| -140 | -1787.84559 | -502.9 | -1012 | -1224.4 | -1167 | -2420.7 |
| -130 | -1623.26 | -355.8 | -899 | -1020.5 | -892.9 | -1939.7 |
| -120 | -1421.63335 | -276.5 | -695.2 | -811.2 | -699.5 | -1463 |
| -110 | -1167.4042 | -215.5 | -524.3 | -609.7 | -263.7 | -1076.7 |
| -100 | -795.646448 | -141.6 | -340.6 | -407.7 | -255.1 | -730 |
| -90 | -418.627722 | -72 | -175.8 | -204.5 | -188.6 | -358.3 |
| -80 | -17.4483147 | 1.2 | -7.3 | 3.1 | 1.2 | 0.6 |
| -70 | 372.961456 | 76.9 | 172.7 | 170.9 | 182.5 | 352.8 |
| -60 | 826.86145 | 160.5 | 356.4 | 310.7 | 352.4 | 705.6 |
| -50 | 1287.29993 | 258.8 | 532.8 | 501.1 | 515.1 | 1018.7 |
| -40 | 1715.38806 | 348.5 | 727.5 | 683 | 665.9 | 1242.7 |

| | mouse#8 | mouse#9 | mouse#9 | mouse#9 | mouse#9 |
|---|---|---|---|---|---|
| | P7 | P8 | P8 | P8 | P8 |
| Voltage (mV) | Ba2+-sensitive | Ba2+-sensitive | Ba2+-sensitive | Ba2+-sensitive | Ba2+-sensitive current (pA) |
| -140 | -1192.6 | -1228 | -293 | -1394 | -1747.4 |
| -130 | -1008.9 | -990.6 | -208.1 | -1174.9 | -1277.5 |
| -120 | -923.5 | -761.7 | -102.5 | -956.4 | -875.9 |
| -110 | -978.4 | -571.9 | -22.6 | -843.5 | -588.4 |
| -100 | -686.6 | -364.4 | -9.6 | -514.5 | -358.3 |
| -90 | -338.7 | -179.4 | -9.2 | -251.5 | -175.2 |
| -80 | -2.4 | -2.4 | 5.5 | -6.1 | 0.6 |
| -70 | 311.9 | 177.6 | 11 | 252.7 | 154.4 |
| -60 | 643.3 | 330.2 | 0 | 513.3 | 305.2 |
| -50 | 977.2 | 468.8 | 2.4 | 756.8 | 444.3 |
| -40 | 1168.2 | 592 | 35.4 | 1005.2 | 707.4 |

**Fig. 3D (b)** Data Source File

| | Ba2+-sensitive current | |
|---|---|---|
| Voltage (mV) | MEAN | SEM |
| -140 | -1172.80559 | 161.102532 |
| -130 | -999.312322 | 161.349476 |
| -120 | -752.967693 | 127.723144 |
| -110 | -601.297021 | 115.676228 |
| -100 | -360.105547 | 74.9768477 |
| -90 | -183.389937 | 44.6852962 |
| -80 | 42.4767369 | 16.3339299 |
| -70 | 267.310238 | 48.0566473 |
| -60 | 440.426513 | 61.4514264 |
| -50 | 652.60717 | 89.4468395 |
| -40 | 829.947708 | 110.086514 |

**Fig. 3F (a)**

Data Source File

Ba2+-sensitive current (pA)

| Voltage (mV) | Control astro#1 | Control astro#2 | Control astro#3 | Control astro#4 |
|---|---|---|---|---|
| -140 | -1387.9 | -1507.6 | -640.3 | -2590.9 |
| -130 | -1139.5 | -1210.3 | -487.7 | -2142.3 |
| -120 | -893.6 | -932.6 | -357.1 | -1759 |
| -110 | -675 | -676.3 | -288.1 | -1343.4 |
| -100 | -443.1 | -423 | -192.3 | -926.5 |
| -90 | -220.9 | -206.3 | 96.4 | -466.9 |
| -80 | -2.4 | -4.3 | -0.6 | -6.1 |
| -70 | 217.3 | 184.9 | 89.1 | 476.7 |
| -60 | 431.5 | 369.9 | 185.5 | 941.8 |
| -50 | 658.6 | 539.6 | 281.4 | 1392.8 |
| -40 | 869.1 | 698.2 | 385.1 | 1832.9 |
| -30 | 1077.3 | 849.6 | 491.3 | 2300.4 |
| -20 | 1295.2 | 1013.2 | 596.3 | 2823.5 |
| -10 | 1518.6 | 1200 | 672.5 | 3219 |
| 0 | 1753.2 | 1419.7 | 745.3 | 3578.2 |

| Voltage (mV) | Control astro#5 | Control astro#6 | Control astro#7 | Control astro#8 |
|---|---|---|---|---|
| -140 | -603.6 | -977.2 | -1729.7 | -1616.2 |
| -130 | -577.4 | -738.5 | -1339.1 | -1308 |
| -120 | -485.8 | -566.4 | -1015.6 | -1079.1 |
| -110 | -407.7 | -431.5 | -741 | -850.2 |
| -100 | -263.7 | -285.6 | -479.1 | -593.9 |
| -90 | -134.9 | -148.3 | -240.5 | -274 |
| -80 | -3.1 | -3.7 | -9.2 | -1.2 |
| -70 | 88.5 | 144.7 | 247.8 | 304.6 |
| -60 | 287.5 | 266.1 | 489.5 | 667.1 |
| -50 | 448.6 | 409.5 | 747.7 | 966.2 |
| -40 | 583.5 | 568.2 | 1029.1 | 1342.8 |
| -30 | 775.1 | 693.4 | 1288.5 | 1891.5 |
| -20 | 1220.7 | 724.5 | 1577.8 | 2915 |
| -10 | 1538.9 | 954 | 1906.7 | 3309.3 |
| 0 | 1877.3 | 1081.5 | 2144.8 | 3502.2 |

**Fig. 3F (b)**

Data Source File

Ba2+-sensitive current (pA)

| Voltage (mV) | Intracellular BAPTA astro#1 | Intracellular BAPTA astro#2 | Intracellular BAPTA astro#3 | Intracellular BAPTA astro#4 |
|---|---|---|---|---|
| -140 | -497.4 | -1178.6 | -403.4 | -343.6 |
| -130 | -436.4 | -845.3 | -347.3 | -308.2 |
| -120 | -368 | -628.7 | -267.9 | 252.1 |
| -110 | -267.3 | -457.2 | -209.4 | -205.7 |
| -100 | -169.7 | -289.3 | -159.3 | -124.5 |
| -90 | -94.6 | -111.1 | -94.6 | -56.2 |
| -80 | -2.4 | 1.2 | 5.5 | 5.5 |
| -70 | 78.7 | 126.3 | 103.8 | 72 |
| -60 | 185.5 | 305.2 | 177.6 | 155 |
| -50 | 263.1 | 486.5 | 263.7 | 214.8 |
| -40 | 346.7 | 522.5 | 330.8 | 313.1 |
| -30 | 460.8 | 597.3 | 371.7 | 383.9 |
| -20 | 538.3 | 634.4 | 346.1 | 457.8 |
| -10 | 595.1 | 666.7 | 406.5 | 598.1 |
| 0 | 640.3 | 702.7 | 433.2 | 647.4 |

| Voltage (mV) | Intracellular BAPTA astro#5 | Intracellular BAPTA astro#6 | Intracellular BAPTA astro#7 |
|---|---|---|---|
| -140 | -448.6 | -865.5 | -243.3 |
| -130 | -386.4 | -743.4 | -139.2 |
| -120 | -260 | -595.7 | -135.3 |
| -110 | -191.7 | -463.3 | -108.6 |
| -100 | -147.7 | -310.7 | -82.6 |
| -90 | -69 | -164.7 | -39.2 |
| -80 | -1.8 | 5.5 | 1.3 |
| -70 | 73.9 | 148.3 | 14.2 |
| -60 | 119 | 297.9 | 20.3 |
| -50 | 162.4 | 449.2 | 36.8 |
| -40 | 219.1 | 615.8 | 70.8 |
| -30 | 203.2 | 799.6 | 90.3 |
| -20 | 191 | 971.7 | 140.3 |
| -10 | 231.2 | 1177.4 | 210.3 |
| 0 | 246.4 | 1367.2 | 222 |

**Fig. 3F (c)**

Data Source File

| | age | GFP(+) | Control<br>Ba2+-sensitive current (pA) |
|---|---|---|---|
| mouse#1 | P6 | astro#1 | -1387.9 |
| mouse#1 | P6 | astro#2 | -1507.6 |
| mouse#1 | P6 | astro#3 | -640.3 |
| mouse#2 | P7 | astro#4 | -2590.9 |
| mouse#2 | P7 | astro#5 | -603.6 |
| mouse#3 | P9 | astro#6 | -977.2 |
| mouse#3 | P9 | astro#7 | -1729.7 |
| mouse#3 | P9 | astro#8 | -1616.2 |
| | | MEAN | -1382 |
| | | SEM | 230.4 |

Intracellular BAPTA

| | age | GFP(+) | Ba2+-sensitive current (pA) |
|---|---|---|---|
| mouse#1 | P6 | astro#1 | -497.4 |
| mouse#1 | P6 | astro#2 | -1178.6 |
| mouse#1 | P6 | astro#3 | -403.4 |
| mouse#2 | P7 | astro#4 | -343.6 |
| mouse#2 | P7 | astro#5 | -448.6 |
| mouse#3 | P9 | astro#6 | -865.5 |
| mouse#4 | P12 | astro#7 | -243.3 |
| | | MEAN | -568.6 |
| | | SEM | 125.7 |

Control vs BAPTAi
Ba2+-sensitive current (pA)

| Mann Whitney test | |
|---|---|
| P value | 0.0059 |
| P value summary | ** |

**Fig. 4C-D (a)**

Data Source File

| Hb9-GFP mice | | age | Neuronal oscill. in oscill. aCSF |
|---|---|---|---|
| mouse#1 | IN1 | P9 | + |
| mouse#2 | IN2 | P10 | + |
| mouse#3 | IN3 | P5 | - |
| mouse#4 | IN4 | P5 | - |
| mouse#4 | IN5 | P5 | + |
| mouse#5 | IN6 | P5 | + |
| mouse#6 | IN7 | P6 | + |
| mouse#6 | IN8 | P6 | + |
| mouse#6 | IN9 | P6 | - |
| mouse#6 | IN10 | P6 | - |
| mouse#7 | IN11 | P7 | - |
| mouse#7 | IN12 | P7 | + |
| mouse#7 | IN13 | P7 | + |
| mouse#8 | IN14 | P12 | + |
| mouse#9 | IN15 | P5 | + |
| mouse#9 | IN16 | P5 | + |
| mouse#10 | IN17 | P6 | - |
| mouse#10 | IN18 | P6 | - |
| mouse#11 | IN19 | P9 | + |
| mouse#11 | IN20 | P9 | + |
| mouse#12 | IN21 | P8 | + |
| mouse#13 | IN22 | P13 | + |
| mouse#14 | IN23 | P7 | + |
| mouse#14 | IN24 | P7 | - |

| | |
|---|---|
| Hb9+ total | 24 |
| Hb9+ oscillatory | 16 |

Hb9 INs

K6Ca0.9

| | | |
|---|---|---|
| oscillatory | % | 66.66666667 |
| non oscillatory | % | 33.33333333 |

**Fig. 4C-D (b)**

Data Source File

| | | Oscill. aCSF Frequ. N oscill. (Hz) | Oscill. aCSF Ampl N oscill (mV) | Oscill. aCSF Duration N oscill (s) | Bath application of Ba2+ |
|---|---|---|---|---|---|
| mouse#1 | IN1 | 0.05 | 4.54666667 | 2.56 | tested, no longer bursting under Barium |
| mouse#2 | IN2 | 0.9153198 | 6.97016042 | 8.996136465 | tested, no longer bursting under Barium |
| mouse#3 | IN3 | | | | not tested |
| mouse#4 | IN4 | | | | not tested |
| mouse#4 | IN5 | 0.275 | 5 | 1.35 | not tested |
| mouse#5 | IN6 | 0.36794739 | 7.12017229 | 2.2 | tested, no longer bursting under Barium |
| mouse#6 | IN7 | 0.53085351 | 5.931331 | 0.96836496 | tested, no longer bursting under Barium |
| mouse#6 | IN8 | 0.1 | 10.6168564 | 7.079337457 | tested, no longer bursting under Barium |
| mouse#6 | IN9 | | | | not tested |
| mouse#6 | IN10 | | | | not tested |
| mouse#7 | IN11 | | | | not tested |
| mouse#7 | IN12 | 0.27328378 | 8.268 | 5.37 | tested, no longer bursting under Barium |
| mouse#7 | IN13 | 0.46851179 | 6.42694052 | 1.345 | tested, no longer bursting under Barium |
| mouse#8 | IN14 | 0.1 | 7.5 | 10.5 | not tested |
| mouse#9 | IN15 | 0.1 | 8.96333333 | 0.7866666667 | not tested |
| mouse#9 | IN16 | 0.1 | 8.36 | 1.5 | not tested |
| mouse#10 | IN17 | | | | not tested |
| mouse#10 | IN18 | | | | not tested |
| mouse#11 | IN19 | 0.40563051 | 4.58128491 | 2.151 | not tested |
| mouse#11 | IN20 | 0.44162478 | 4.65408099 | 5.416 | not tested |
| mouse#12 | IN21 | 1.5 | 0.31 | 2.08 | tested, no longer bursting under Barium |
| mouse#13 | IN22 | 1.6 | 4.15 | 0.576 | tested, no longer bursting under Barium |
| mouse#14 | IN23 | 0.3 | 6.44 | 1.2 | not tested |
| mouse#14 | IN24 | | | | not tested |
| | MEAN | 0.47051072 | 6.23992666 | 3.379906597 | n=9/9 no longer bursting under Barium |
| | SEM | 0.11875739 | 0.60363729 | 0.780608271 | |

**Fig. 4C-D (c)**

Data Source File

| | Hb9-GFP mice | age | Neuronal oscill. in TTX + Cocktail |
|---|---|---|---|
| mouse#1 | IN1 | P6 | + |
| mouse#1 | IN2 | P6 | + |
| mouse#1 | IN3 | P6 | - |
| mouse#1 | IN4 | P6 | + |
| mouse#2 | IN5 | P10 | + |
| mouse#2 | IN6 | P10 | + |
| mouse#2 | IN7 | P10 | + |
| mouse#3 | IN8 | P12 | + |
| mouse#3 | IN9 | P12 | - |
| mouse#4 | IN10 | P13 | + |
| mouse#4 | IN11 | P13 | - |
| mouse#5 | IN12 | P13 | + |
| mouse#5 | IN13 | P13 | + |
| | | Hb9+ total | 13 |
| | | Hb9+ oscillatory | 11 |

| Hb9 INs | |
|---|---|
| TTX+Cocktail oscillatory % | 84.61538462 |
| non oscillatory % | 15.38461538 |

**Fig. 4C-D (d)**

Data Source File

| | | TTX+cocktail | TTX+cocktail | TTX+ Cocktail | Bath application of Ba2+ |
|---|---|---|---|---|---|
| | | Frequ. N oscill. (Hz) | Ampl N oscill (mV) | Duration N oscill (s) | |
| mouse#1 | IN1 | 1.5 | 12.3833333 | 0.62066666 | tested, no longer bursting under Barium |
| mouse#1 | IN2 | 0.16 | 29.04375 | 6.74375 | tested, no longer bursting under Barium |
| mouse#1 | IN3 | | | | not tested |
| mouse#1 | IN4 | 1.1 | 7.23545455 | 0.643636 | tested, no longer bursting under Barium |
| mouse#2 | IN5 | 0.60606061 | 8.36 | 0.91 | not tested |
| mouse#2 | IN6 | 2 | 6.17636364 | 0.399090909 | not tested |
| mouse#2 | IN7 | 0.5 | 32.383 | 1.274 | tested, no longer bursting under Barium |
| mouse#3 | IN8 | 1 | 4.18 | 0.799 | not tested |
| mouse#3 | IN9 | | | | not tested |
| mouse#4 | IN10 | 2.4 | 5.31 | 0.43 | not tested |
| mouse#4 | IN11 | | | | not tested |
| mouse#5 | IN12 | 3 | 4.55 | 0.1 | tested, no longer bursting under Barium |
| mouse#5 | IN13 | 0.83333333 | 22 | 1.15 | tested, no longer bursting under Barium |
| | MEAN | 1.30993939 | 13.1621902 | 1.307014394 | n=6/6 no longer bursting under Barium |
| | SEM | 1.18222222 | 17.9325896 | 1.755342172 | |

**Fig. 4F** Data Source File

| | | | |
|---|---|---|---|
| | | Oscillatory aCSF | Oscillatory aCSF |
| | Age | Still oscillating under Ba2+ | No longer oscillating under Ba2+ |
| mouse#1 | P13 | n=1 | n=1 |
| mouse#2 | P13 | n=1 | n=6 |
| | Total ROIs | n=2 | n=7 |
| | % | 22.22222222 | 77.7777778 |

| | | | |
|---|---|---|---|
| | | TTX+Cocktail | TTX+Cocktail |
| | Age | Still oscillating under Ba2+ | No longer oscillating under Ba2+ |
| mouse#1 | P16 | n=1 | n=3 |
| mouse#2 | P16 | n=2 | n=12 |
| mouse#3 | P13 | n=0 | n=10 |
| | Total ROIs | n=3 | n=25 |
| | % | 10.71428571 | 89.2857143 |

**Fig. 4G (a)**

Data Source File

| | Oscill. aCSF Amplitude (df/f) | Ba2+ Amplitude (df/f) | Oscill. aCSF Duration (ms) | Ba2+ Duration (ms) | Oscill. aCSF Frequency (Events Nb/min) | Ba2+ Frequency (Events Nb/min) |
|---|---|---|---|---|---|---|
| 68 / UG / ROI: | 0.28852316 | 0.29494372 | 1305.27468 | 1335.68408 | 1.26315789 | 1.58490566 |
| 69 / UG / ROI: | 0.38152904 | 0.4148512 | 1303.30527 | 1078.62651 | 2.52631579 | 4.0754717 |
| 38 / UG / ROI: | 0.31914027 | 0.36531867 | 1028.87879 | 1289.56007 | 15.7894737 | 11.7735849 |
| 59 / UG / ROI: | 0.32748687 | 0.33360327 | 962.508007 | 1213.89701 | 8.21052632 | 7.01886792 |
| 44 / UG / ROI: | 0.51208406 | 0.52586653 | 872.202598 | 822.225867 | 27.7894737 | 30.7924528 |
| 55 / UG / ROI: | 0.31282242 | 0.33327052 | 1232.69866 | 1277.27962 | 10.1052632 | 11.3207547 |
| 49 / UG / ROI: | 0.39146181 | 0.38333446 | 1079.42828 | 1054.65941 | 15.7894737 | 17.2075472 |
| 40 / UG / ROI: | 0.30332239 | 0.3134104 | 925.321067 | 1195.34075 | 5.05263158 | 3.39622642 |
| 63 / UG / ROI: | 0.26789174 | 0.37426776 | 1918.10755 | 1057.66258 | 0.63157895 | 0.90566038 |
| 78 / UG / ROI: | 0.36034426 | 0.35071705 | 1262.36734 | 1093.80949 | 10.1052632 | 11.3207547 |
| 60 / UG / ROI: | 0.3503458 | 0.34084693 | 944.417607 | 1402.43057 | 5.68421053 | 4.30188679 |
| 64 / UG / ROI: | 0.5377941 | 0.58550402 | 834.293887 | 793.848966 | 29.0526316 | 30.3396226 |
| 22 / UG / ROI: | 0.37757319 | 0.36843816 | 980.970036 | 1057.49516 | 13.8947368 | 15.3962264 |
| 41 / UG / ROI: | 0.33557444 | 0.62892673 | 835.714635 | 2181.21136 | 1.89473684 | 2.71698113 |
| 9 / UG / ROI:{( | 0.36302551 | 0.36825191 | 1112.44503 | 999.176342 | 13.2631579 | 14.0377358 |
| 61 / UG / ROI: | 0.32003349 | 0.31662156 | 1445.3652 | 1242.79806 | 5.68421053 | 6.11320755 |
| 16 / UG / ROI: | 0.34301748 | 0.32991305 | 1067.386 | 1203.88009 | 9.33333333 | 6.44444444 |
| 47 / UG / ROI: | 0.31947652 | 0.33979181 | 994.743902 | 1012.56812 | 10 | 6.88888889 |
| 64 / UG / ROI: | 0.34533786 | 0.3488755 | 1058.64512 | 1215.1369 | 14.6666667 | 10.4444444 |
| 55 / UG / ROI: | 0.26370793 | 0.34518201 | 1887.85024 | 1151.70876 | 2.66666667 | 3.55555556 |
| 57 / UG / ROI: | 0.34676445 | 0.37634016 | 1037.56471 | 1188.36782 | 15.3333333 | 14.8888889 |
| 19 / UG / ROI: | 0.38658918 | 0.35699086 | 1123.80082 | 1126.13803 | 17.3333333 | 14 |
| 58 / UG / ROI: | 0.32579633 | 0.34817442 | 874.94134 | 1186.88069 | 8 | 7.11111111 |
| 30 / UG / ROI: | 0.29608325 | 0.32869825 | 957.623846 | 1410.18662 | 4 | 4.66666667 |
| MEAN | 0.349 | 0.378 | 1127 | 1191 | 10.34 | 10.01 |
| | 0.01313 | 0.01703 | 58.58 | 53.16 | 1.545 | 1.611 |

| Amplitude (df/f) | |
|---|---|
| Wilcoxon matched-pairs signed rank test | |
| P value | 0.0079 |
| P value summary | ** |

| Duration (ms) | |
|---|---|
| Wilcoxon matched-pairs signed rank test | |
| P value | 0.2768 |
| P value summary | ns |

| Frequency (Events Nb/min) | |
|---|---|
| Wilcoxon matched-pairs signed rank test | |
| P value | 0.71 |
| P value summary | ns |

| | TTX+Cocktail Amplitude (df/f) | Ba2+ Amplitude (df/f) | TTX+Cocktail Duration (ms) | Ba2+ Duration (ms) | TTX+Cocktail Frequency (Events Nb/min) | Ba2+ Frequency (Events Nb/min) |
|---|---|---|---|---|---|---|
| 53 / UG / ROI: | 0.48252146 | 0.466840635 | 2604.27462 | 2577.43361 | 5.05263158 | 3.31034483 |
| 60 / UG / ROI: | 0.358202 | 0.326995915 | 1072.3678 | 1030.55223 | 15.7894737 | 16.5517241 |
| 54 / UG / ROI: | 0.34197094 | 0.3006918 | 3668.20481 | 4472.42016 | 3.15789474 | 0.82758621 |
| 67 / UG / ROI: | 0.302490155 | 0.278337485 | 1494.8044 | 988.994761 | 4.42105263 | 7.44827586 |
| 64 / UG / ROI: | 0.37004571 | 0.319300565 | 1000.64168 | 1226.94874 | 6.94736842 | 12.4137931 |
| 43 / UG / ROI: | 0.279385095 | 0.279384715 | 1064.41076 | 1286.75329 | 4.42105263 | 4.13793103 |
| 51 / UG / ROI: | 0.270246405 | 0.280248575 | 1380.74744 | 1120.51198 | 4.42105263 | 5.79310345 |
| 72 / UG / ROI: | 0.29800045 | 0.27310868 | 1352.91595 | 1598.88933 | 3.15789474 | 4.55172414 |
| 61 / UG / ROI: | 0.359377485 | 0.338562335 | 1488.46679 | 1822.5168 | 1.26315789 | 3.31034483 |
| 52 / UG / ROI: | 0.349533385 | 0.279416275 | 1335.96536 | 1678.67621 | 1.89473684 | 1.24137931 |
| 49 / UG / ROI: | 0.314239015 | 0.31707855 | 904.984554 | 1160.29279 | 10.1052632 | 13.6551724 |

Data Source File

**Fig. 4G (b)**

| | TTX+Cocktail | Ba2+ | TTX+Cocktail | Ba2+ | TTX+Cocktail | Ba2+ |
|---|---|---|---|---|---|---|
| | Amplitude (df/f) | Amplitude (df/f) | Duration (ms) | Duration (ms) | Frequency (Events Nb/min) | Frequency (Events Nb/min) |
| 50 / UG / ROI: | 0.37242607 | 0.32740179 | 1449.55112 | 1066.42973 | 11.3684211 | 13.2413793 |
| 44 / UG / ROI: | 0.303914655 | 0.27869746 | 1453.19652 | 960.206457 | 3.78947368 | 3.31034483 |
| 46 / UG / ROI: | 0.343317795 | 0.326587155 | 948.647893 | 1392.43017 | 7.57894737 | 6.20689655 |
| 45 / UG / ROI: | 0.51363784 | 0.29542479 | 1197.21001 | 1544.84984 | 4.42105263 | 5.79310345 |
| 47 / UG / ROI: | 0.30354382 | 0.31936215 | 1494.76082 | 1107.48797 | 6.94736842 | 14.0689655 |
| 69 / UG / ROI: | 0.360671155 | 0.292321035 | 1336.92508 | 1003.05429 | 1.26315789 | 3.31034483 |
| 65 / UG / ROI: | 0.382487155 | 0.30306208 | 987.522462 | 1288.35426 | 4.42105263 | 4.96551724 |
| 77 / UG / ROI: | 0.375494775 | 0.344432335 | 954.886977 | 1109.05099 | 13.8947368 | 15.7241379 |
| 78 / UG / ROI: | 0.269425415 | 0.38005879 | 850.616736 | 1298.54871 | 5.05263158 | 7.44827586 |
| 80 / UG / ROI: | 0.37397423 | 0.39776399 | 832.860362 | 921.553088 | 25.8947368 | 24.8275862 |
| 37 / UG / ROI: | 0.286922235 | 0.319465225 | 1057.92718 | 1133.39943 | 9.47368421 | 11.5862069 |
| 38 / UG / ROI: | 0.4277196 | 0.41105047 | 848.112581 | 842.902646 | 23.3684211 | 27.3103448 |
| 39 / UG / ROI: | 0.38673794 | 0.35935123 | 1062.43598 | 1101.94131 | 18.3157895 | 20.2758621 |
| 73 / UG / ROI: | 0.460460735 | 0.28387903 | 2456.49776 | 2321.34036 | 1.89473684 | 1.65517241 |
| 55 / UG / ROI: | 0.282462655 | 0.3460207 | 1533.52369 | 1320.15825 | 3.78947368 | 4.96551724 |
| 58 / UG / ROI: | 0.2708096 | 0.31360422 | 1409.81986 | 1448.7953 | 3.78947368 | 3.72413793 |
| 62 / UG / ROI: | 0.34979322 | 0.34961499 | 1872.11344 | 1276.27837 | 1.89473684 | 4.13793103 |
| 74 / UG / ROI: | 0.494474725 | 0.51444915 | 1105.95238 | 1085.09941 | 5.05263158 | 5.37931034 |
| 56 / UG / ROI: | 0.27594316 | 0.24480685 | 1130.49345 | 2993.63389 | 2.52631579 | 1.24137931 |
| 68 / UG / ROI: | 0.34279158 | 0.30893926 | 1682.73011 | 1936.79109 | 1.26315789 | 3.31034483 |
| 71 / UG / ROI: | 0.3090143 | 0.29466011 | 1905.69472 | 1522.10229 | 1.89473684 | 2.06896552 |
| 63 / UG / ROI: | 0.255024935 | 0.29695296 | 1254.25066 | 1725.4693 | 1.26315789 | 2.48275862 |
| 76 / UG / ROI: | 0.29730816 | 0.30091662 | 1080.73703 | 1304.39094 | 4.42105263 | 4.13793103 |
| 37 / UG / ROI: | 0.286922235 | 0.31946523 | 1057.92718 | 1133.39943 | 9.47368421 | 11.5862069 |
| 38 / UG / ROI: | 0.4277196 | 0.41105047 | 848.112581 | 842.902646 | 23.3684211 | 27.3103448 |
| 39 / UG / ROI: | 0.38673794 | 0.35935123 | 1062.43598 | 1101.94131 | 18.3157895 | 20.2758621 |
| 10 / UG / ROI: | 0.496394585 | 0.3661041 | 1467.80659 | 2401.77354 | 2.96296296 | 2.15053763 |
| 80 / UG / ROI: | 0.328630345 | 0.3979327 | 1235.52151 | 1277.05706 | 6.66666667 | 9.46236559 |
| 32 / UG / ROI: | 0.363620665 | 0.43223885 | 1019.0093 | 1814.44813 | 0.74074074 | 0.43010753 |
| 48 / UG / ROI: | 0.283284835 | 0.331345265 | 1491.0014 | 1219.38998 | 4.44444444 | 5.59139785 |
| 101 / UG / RO | 0.3311333 | 0.366832895 | 982.927328 | 1077.85024 | 10.3703704 | 13.1182796 |
| 63 / UG / ROI: | 0.295055995 | 0.299958025 | 1573.58029 | 1225.72285 | 1.48148148 | 2.3655914 |
| 73 / UG / ROI: | 0.262567575 | 0.315608145 | 2401.43604 | 4071.68835 | 0.74074074 | 1.93548387 |
| 50 / UG / ROI: | 0.26843738 | 0.300158745 | 1640.79811 | 1696.4302 | 0.74074074 | 1.50537634 |
| 42 / UG / ROI: | 0.351954 | 0.38248169 | 1173.97518 | 1138.37233 | 4.44444444 | 10.7526882 |
| 81 / UG / ROI: | 0.447371895 | 0.40724462 | 970.808532 | 1095.48707 | 13.3333333 | 13.3333333 |
| 86 / UG / ROI: | 0.34286495 | 0.38276523 | 1231.60296 | 1144.17545 | 5.18518519 | 8.17204301 |
| 12 / UG / ROI: | 0.29535657 | 0.30366741 | 1891.69156 | 1239.05689 | 1.48148148 | 1.93548387 |
| 56 / UG / ROI: | 0.257521185 | 0.323916395 | 3883.93827 | 1507.73252 | 0.74074074 | 1.50537634 |
| 49 / UG / ROI: | 0.356522515 | 0.383684825 | 1153.60788 | 1203.49797 | 5.92592593 | 6.88172043 |
| 57 / UG / ROI: | 0.47182195 | 0.6954864 | 2371.32688 | 4742.65375 | 0.32258065 | 0.64516129 |
| 4 / UG / ROI:{ | 0.307315825 | 0.32817444 | 1213.46516 | 1424.47472 | 3.7037037 | 5.80645161 |
| 69 / UG / ROI: | 0.361306995 | 0.325778945 | 1119.76358 | 1197.45594 | 2.96296296 | 4.08602151 |
| 72 / UG / ROI: | 0.390426735 | 0.35527333 | 1157.13828 | 1118.54574 | 8.88888889 | 11.1827957 |
| 21 / UG / ROI: | 0.361035445 | 0.530157305 | 1209.92447 | 875.322507 | 8.88888889 | 8.17204301 |
| 71 / UG / ROI: | 0.32714711 | 0.390501055 | 1192.32091 | 1082.71911 | 5.92592593 | 8.8172043 |
| 19 / UG / ROI: | 0.32252415 | 0.49355566 | 6570.6022 | 13141.2044 | 0.10752688 | 0.21505376 |
| 44 / UG / ROI: | 0.57610541 | 0.51318206 | 775.290089 | 776.999023 | 32.5925926 | 32.2580645 |
| 52 / UG / ROI: | 0.25126203 | 0.459977235 | 3021.13038 | 4688.9249 | 0.74074074 | 0.21505376 |
| 46 / UG / ROI: | 0.441813595 | 0.492277875 | 1165.27512 | 1122.06512 | 3.7037037 | 6.66666667 |

Data Source File

**Fig. 4G (c)**

| | TTX+Cocktail Amplitude | Ba2+ Amplitude | TTX+Cocktail Duration | Ba2+ Duration | TTX+Cocktail Frequency (Events Nb/min) | Ba2+ Frequency (Events Nb/min) |
|---|---|---|---|---|---|---|
| 64 / UG / ROI: | 0.35441916 | 0.433911475 | 1017.52663 | 961.727611 | 17.037037 | 20.2150538 |
| 15 / UG / ROI: | 0.35982307 | 0.38753904 | 1064.34882 | 1129.34096 | 0.74074074 | 2.79569892 |
| 95 / UG / ROI: | 0.297627515 | 0.454570495 | 1536.67099 | 1406.7263 | 2.22222222 | 4.30107527 |
| 87 / UG / ROI: | 0.35594263 | 0.368002845 | 1410.04842 | 1078.62505 | 5.92592593 | 6.02150538 |
| 83 / UG / ROI: | 0.42795364 | 0.34571678 | 831.55545 | 1334.53698 | 1.48148148 | 3.01075269 |
| 30 / UG / ROI: | 0.32018535 | 0.33640819 | 895.844882 | 1132.11178 | 3.7037037 | 3.87096774 |
| 18 / UG / ROI: | 0.297786475 | 0.357753075 | 1914.49707 | 1297.04494 | 2.22222222 | 2.79569892 |
| 23 / UG / ROI: | 0.36296928 | 0.359181495 | 1027.35421 | 1101.68846 | 1.48148148 | 5.37634409 |
| 53 / UG / ROI: | 0.356155855 | 0.363266315 | 1311.7283 | 1231.31784 | 10.3703704 | 14.6236559 |
| 54 / UG / ROI: | 0.2215548 | 0.47988526 | 1068.45478 | 2136.90956 | 0.43010753 | 0.86021505 |
| 60 / UG / ROI: | 0.47444941 | 0.485222835 | 948.895236 | 924.422289 | 20 | 20.2150538 |
| 84 / UG / ROI: | 0.350230045 | 0.32203502 | 1036.23339 | 1196.16162 | 2.96296296 | 7.95698925 |
| 28 / UG / ROI: | 0.281270215 | 0.32501932 | 1222.45132 | 1199.61585 | 4.44444444 | 3.65591398 |
| 9 / UG / ROI:{( | 0.29248796 | 0.328765935 | 1252.57223 | 2054.83829 | 2.22222222 | 1.07526882 |
| 3 / UG / ROI:{( | 0.1235689 | 0.21568978 | 770.244445 | 1540.48889 | 0.7526882 | 1.5053764 |
| 25 / UG / ROI: | 0.25314779 | 0.36763797 | 2037.49008 | 1560.44179 | 0.74074074 | 0.64516129 |
| 14 / UG / ROI: | 0.28848378 | 0.33798067 | 1438.16841 | 1177.40773 | 1.48148148 | 1.72043011 |
| 29 / UG / ROI: | 0.34287541 | 0.38299934 | 1292.63897 | 1007.72505 | 3.7037037 | 4.7311828 |
| 24 / UG / ROI: | 0.19672395 | 0.39344789 | 1484.0471 | 2968.0942 | 0.43010753 | 0.86021505 |
| 43 / UG / ROI: | 0.50407179 | 0.38039922 | 1721.35741 | 1072.8672 | 8.14814815 | 8.8172043 |
| 67 / UG / ROI: | 0.33031468 | 0.352824385 | 1210.86243 | 1248.07709 | 6.66666667 | 7.09677419 |
| 51 / UG / ROI: | 0.30584321 | 0.3173152 | 4228.08008 | 1215.59865 | 1.48148148 | 1.50537634 |
| 55 / UG / ROI: | 0.372569355 | 0.51440076 | 932.820025 | 929.508315 | 17.037037 | 24.7311828 |
| 5 / UG / ROI:{! | 0.357342435 | 0.411632105 | 1405.75332 | 1718.06005 | 0.74074074 | 3.01075269 |
| 88 / UG / ROI: | 0.39225729 | 0.334141165 | 1086.0595 | 1097.69341 | 4.44444444 | 9.89247312 |
| 74 / UG / ROI: | 0.411317115 | 0.465590325 | 1006.50971 | 939.570345 | 18.5185185 | 20.8602151 |
| 85 / UG / ROI: | 0.30734387 | 0.32977438 | 2039.27124 | 1233.70711 | 2.96296296 | 5.37634409 |
| 13 / UG / ROI: | 0.317058225 | 0.34431522 | 1355.37369 | 1168.32383 | 2.22222222 | 4.08602151 |
| 100 / UG / RO | 0.37215685 | 0.36673452 | 1248.75439 | 1111.36837 | 14.0740741 | 14.6236559 |
| 92 / UG / ROI: | 0.397464245 | 0.400364565 | 851.598554 | 1028.21542 | 16.2962963 | 18.0645161 |
| 31 / UG / ROI: | 0.25275036 | 0.34743317 | 3359.20308 | 1084.91036 | 0.74074074 | 1.29032258 |
| 59 / UG / ROI: | 0.28809308 | 0.31848159 | 1618.542 | 1305.37041 | 2.96296296 | 5.59139785 |
| 62 / UG / ROI: | 0.26905092 | 0.351781555 | 1185.80846 | 1460.52945 | 0.74074074 | 1.72043011 |
| 33 / UG / ROI: | 0.32952572 | 0.368078955 | 1323.27327 | 1039.00978 | 8.88888889 | 9.67741935 |
| 6 / UG / ROI:{( | 0.31326474 | 0.344810915 | 2255.98582 | 1778.18033 | 0.74074074 | 1.72043011 |
| 97 / UG / ROI: | 0.36451209 | 0.34892101 | 1320.99914 | 1242.66242 | 9.62962963 | 11.6129032 |
| 16 / UG / ROI: | 0.34217449 | 0.357738555 | 1669.10607 | 1253.14588 | 3.7037037 | 6.02150538 |
| 17 / UG / ROI: | 0.345380525 | 0.41776078 | 994.533442 | 1272.21613 | 13.3333333 | 13.3333333 |
| 61 / UG / ROI: | 0.20888039 | 0.41776078 | 2762.3061 | 5524.6122 | 0.10752688 | 0.21505376 |
| 37 / UG / ROI: | 0.18572594 | 0.37145187 | 763.477587 | 1526.95517 | 0.96774194 | 1.93548387 |
| 79 / UG / ROI: | 0.404084335 | 0.42655774 | 1011.79858 | 917.814599 | 17.7777778 | 20 |
| 105 / UG / RO | 0.282819015 | 0.36976738 | 1163.92992 | 1395.5437 | 5.18518519 | 3.87096774 |
| 106 / UG / RO | 0.5041891 | 0.50078816 | 883.96743 | 879.454875 | 27.4074074 | 26.6666667 |
| 36 / UG / ROI: | 0.54404304 | 0.33881083 | 1194.65373 | 1318.18518 | 4.44444444 | 4.94623656 |
| 68 / UG / ROI: | 0.346419795 | 0.38805229 | 907.762608 | 1056.36458 | 11.8518519 | 15.6989247 |
| 45 / UG / ROI: | 0.366337245 | 0.38419493 | 1179.87991 | 1108.44938 | 16.2962963 | 15.0537634 |
| 78 / UG / ROI: | 0.29833253 | 0.36427177 | 1592.52643 | 1290.92126 | 6.66666667 | 8.8172043 |
| 70 / UG / ROI: | 0.279298215 | 0.3386783 | 2165.88676 | 2228.61435 | 1.48148148 | 0.43010753 |
| 40 / UG / ROI: | 0.25005349 | 0.50010699 | 1225.08868 | 2450.17737 | 0.21505376 | 0.43010753 |
| 41 / UG / ROI: | 0.3052892 | 0.31936427 | 1515.20272 | 1308.52397 | 2.96296296 | 6.02150538 |

Data Source File

**Fig. 4G (d)**

| | TTX+Cocktail | Ba2+ | TTX+Cocktail | Ba2+ | TTX+Cocktail | Ba2+ |
|---|---|---|---|---|---|---|
| | Amplitude | Amplitude | Duration | Duration | Frequency (Events Nb/min) | Frequency (Events Nb/min) |
| 77 / UG / ROI: | 0.294906935 | 0.37401755 | 2300.84704 | 1265.4092 | 1.48148148 | 4.94623656 |
| 65 / UG / ROI: | 0.18196016 | 0.36392033 | 655.890814 | 1311.78163 | 0.64516129 | 1.29032258 |
| 39 / UG / ROI: | 0.299640435 | 0.35805313 | 1545.72796 | 1515.73549 | 3.7037037 | 4.51612903 |
| 75 / UG / ROI: | 0.282769225 | 0.36273374 | 2443.17146 | 2238.73236 | 0.74074074 | 0.86021505 |
| 76 / UG / ROI: | 0.326698465 | 0.40471022 | 1260.64157 | 1181.6505 | 7.40740741 | 8.60215054 |
| 35 / UG / ROI: | 0.31550708 | 0.35028797 | 2314.49472 | 1502.14559 | 1.48148148 | 3.22580645 |
| 94 / UG / ROI: | 0.32662796 | 0.33003282 | 1684.21918 | 1313.3042 | 7.40740741 | 7.95698925 |
| 58 / UG / ROI: | 0.397921855 | 0.43674536 | 846.788495 | 961.212822 | 21.4814815 | 20.2150538 |
| 66 / UG / ROI: | 0.536648445 | 0.45805758 | 1576.08979 | 1350.04052 | 7.40740741 | 7.74193548 |
| 52 / UG / ROI: | 0.350598978 | 0.333893529 | 1148.590067 | 1076.201743 | 11.1764706 | 11.4953271 |
| 55 / UG / ROI: | 0.328428346 | 0.360500715 | 1493.504149 | 1021.013244 | 7.05882353 | 8.13084112 |
| 56 / UG / ROI: | 0.41798337 | 0.354077744 | 1061.369706 | 1111.684529 | 10 | 9.25233645 |
| 57 / UG / ROI: | 0.384940678 | 0.378359278 | 974.5438689 | 1028.015693 | 2.35294118 | 2.80373832 |
| 59 / UG / ROI: | 0.390955696 | 0.398980047 | 934.5174131 | 1199.791666 | 18.2352941 | 17.3831776 |
| 62 / UG / ROI: | *0.49171893* | 0.493649833 | *875.06788* | 918.1210462 | 22.9411765 | 21.8691589 |
| 63 / UG / ROI: | 0.518932751 | 0.389884024 | 1297.738837 | 1223.456217 | 5.29411765 | 4.20560748 |
| 64 / UG / ROI: | 0.457053251 | 0.425455239 | 969.7597087 | 953.9940122 | 16.4705882 | 14.8598131 |
| 68 / UG / ROI: | 0.74519547 | 0.726656165 | *444.217007* | 479.4130727 | 38.3529412 | 35.7196262 |
| 69 / UG / ROI: | 0.46827982 | 0.459655305 | 992.2566818 | 924.913341 | 20 | 17.3831776 |
| 71 / UG / ROI: | 0.550641729 | 0.596474525 | 680.9279003 | 782.1910045 | 32.3529412 | 32.2429907 |
| 76 / UG / ROI: | 0.373203446 | 0.372569274 | 1215.460014 | 1103.843888 | 11.7647059 | 9.53271028 |
| 81 / UG / ROI: | 0.446088817 | 0.475431442 | 868.7864635 | 847.5110446 | 22.9411765 | 22.7102804 |
| 89 / UG / ROI: | 0.668814021 | 0.566144561 | 785.2774076 | 743.1110654 | 28.8235294 | 31.9626168 |
| 90 / UG / ROI: | 0.525813852 | *0.44857416* | 955.9022366 | *855.589591* | 20 | 19.3457944 |
| 93 / UG / ROI: | 0.391219272 | 0.411094976 | 1065.076524 | 1108.643749 | 12.9411765 | 15.4205607 |
| 31 / UG / ROI: | 0.72742345 | 0.71110386 | 488.863863 | 549.2118872 | 37.6470588 | 37.7570093 |
| 60 / UG / ROI: | 0.537879215 | 0.541562655 | 838.56981 | 772.037032 | 32.3529412 | 27.4766355 |
| 65 / UG / ROI: | 0.468381855 | 0.35668294 | 946.766857 | 1136.49038 | 14.7058824 | 10.0934579 |
| 66 / UG / ROI: | 0.54326482 | 0.539893095 | 841.294086 | 833.704019 | 22.3529412 | 24.1121495 |
| 72 / UG / ROI: | 0.730078595 | 0.362592585 | 852.465626 | 1187.40624 | 1.76470588 | 2.24299065 |
| 73 / UG / ROI: | 0.38333707 | 0.357472065 | 1082.04569 | 1143.88914 | 4.70588235 | 6.72897196 |
| 74 / UG / ROI: | 0.3447834 | 0.387694725 | 921.810524 | 1021.83259 | 16.4705882 | 15.4205607 |
| 75 / UG / ROI: | 0.28924283 | 0.278469245 | 2320.5697 | 1811.24804 | 1.76470588 | 1.96261682 |
| 77 / UG / ROI: | 0.748670015 | 0.72095165 | 495.865023 | 518.487179 | 25.2941176 | 55.5140187 |
| 78 / UG / ROI: | 0.27857467 | 0.293996445 | 1790.41366 | 1059.81147 | 1.17647059 | 1.68224299 |
| 79 / UG / ROI: | 0.12227389 | 0.24454777 | 419.074074 | 838.148148 | 0.14018692 | 0.28037383 |
| 80 / UG / ROI: | 0.386690925 | 0.39119472 | 1064.93103 | 1080.99585 | 10.5882353 | 11.7757009 |
| 82 / UG / ROI: | 0.28757048 | 0.33688019 | 1492.68182 | 1629.76175 | 2.94117647 | 3.08411215 |
| 83 / UG / ROI: | 0.5949646 | 0.62702387 | 727.924352 | 766.339101 | 36.4705882 | 33.364486 |
| 84 / UG / ROI: | 0.37916707 | 0.419214625 | 877.54043 | 889.960678 | 18.8235294 | 19.6261682 |
| 85 / UG / ROI: | 0.35148428 | 0.3730932 | 986.046976 | 1117.87071 | 11.1764706 | 10.6542056 |
| 94 / UG / ROI: | 0.319749795 | 0.353816505 | 1108.21958 | 1232.12997 | 3.52941176 | 3.92523364 |
| 95 / UG / ROI: | 0.19082525 | 0.3816505 | 1288.91455 | 2577.82911 | 0.28037383 | 0.56074766 |
| 21 / UG / ROI: | 0.319146685 | 0.31889718 | 1484.0714 | 1184.22314 | 4.11764706 | 5.04672897 |
| 23 / UG / ROI: | 0.392383475 | 0.37342958 | 1180.16813 | 1014.17006 | 3.52941176 | 5.04672897 |
| 24 / UG / ROI: | 0.251965845 | 0.291809275 | 1997.51723 | 2696.46909 | 0.58823529 | 0.28037383 |
| MEAN | 0.363 | 0.3813 | 1379 | 1463 | 8.257 | 9.366 |
| SEM | 0.008633 | 0.007073 | 59 | 95.65 | 0.7006 | 0.7423 |

| Amplitude (df/f) | |
|---|---|
| Wilcoxon matched-pairs signed rank test | |
| P value | 0.0004 |
| P value summary | *** |

| Duration (ms) | |
|---|---|
| Wilcoxon matched-pairs signed rank test | |
| P value | 0.4342 |
| P value summary | ns |

| Frequency (Events Nb/min) | |
|---|---|
| Wilcoxon matched-pairs signed | |
| P value | <0.0001 |
| P value summary | **** |

**Fig. 5D (a)**

Data Source File

| RVL2 | age | Oscill. aCSF<br>burst amplitu | +Barium<br>burst amplitude (mV) |
|---|---|---|---|
| mouse#1 | P3 | 9.68724444 | 9.44801587 |
| mouse#2 | P4 | 2.04364122 | 3.32192308 |
| mouse#3 | P3 | 2.98541667 | 11.8294048 |
| mouse#4 | P4 | 12.5625 | 11.1975 |
| mouse#5 | P3 | 7.22470588 | 7.63422619 |
| mouse#6 | P4 | 9.33333333 | 15.8912281 |
| | MEAN | 7.306 | 9.887 |
| | SEM | 1.671 | 1.731 |

Amplitude
Wilcoxon matched-pairs signed rank test

| | |
|---|---|
| P value | 0.3125 |
| P value summ | ns |

| RVL2 | age | Oscill. aCSF<br>burst duratior | +Barium<br>burst duration (ms) |
|---|---|---|---|
| mouse#1 | P3 | 993.402667 | 1691.60079 |
| mouse#2 | P4 | 895.352467 | 1264.08659 |
| mouse#3 | P3 | 385.083333 | 1219.40476 |
| mouse#4 | P4 | 400.8125 | 1023.7875 |
| mouse#5 | P3 | 837.287798 | 1167.55595 |
| mouse#6 | P4 | 393.5 | 845.777193 |
| | MEAN | 650.9 | 1202 |
| | SEM | 117.1 | 116 |

Duration
Wilcoxon matched-pairs signed rank test

| | |
|---|---|
| P value | 0.0313 |
| P value summ | * |

| RVL2 | age | Oscill. aCSF<br>burst frequen | +Barium<br>burst frequency (Hz) |
|---|---|---|---|
| mouse#1 | P3 | 0.96702006 | 0.84797937 |
| mouse#2 | P4 | 1.13795709 | 0.64034882 |
| mouse#3 | P3 | 2.08461067 | 0.67534975 |
| mouse#4 | P4 | 2.11267606 | 0.59259259 |
| mouse#5 | P3 | 1.01437262 | 0.77362244 |
| mouse#6 | P4 | 1.5 | 0.95833333 |
| | MEAN | 1.469 | 0.748 |
| | SEM | 0.2131 | 0.05654 |

Frequency
Wilcoxon matched-pairs signed rank test

| | |
|---|---|
| P value | 0.0313 |
| P value summ | * |

**Fig. 5D (b)**

Data Source File

| | | Cocktail | +Barium |
|---|---|---|---|
| RVL2 | | burst amplitu | burst amplitude (mV) |
| mouse#1 | P3 | 9.68153768 | 8.63862608 |
| mouse#2 | P3 | 3.21763467 | 5.15845209 |
| mouse#3 | P3 | 5.50809779 | 3.45863188 |
| mouse#4 | P4 | 3.81435353 | 3.09385522 |
| mouse#5 | P3 | 9.64040573 | 8.98513735 |
| mouse#6 | P4 | 9.4318811 | 18.5644366 |
| mouse#7 | P3 | 2.78572452 | 3.01211802 |
| | MEAN | 6.297 | 7.273 |
| | SEM | 1.206 | 2.11 |

Amplitude
Wilcoxon matched-pairs signed rank test

| | |
|---|---|
| P value | 0.9375 |
| P value summ | ns |

| | | Cocktail | +Barium |
|---|---|---|---|
| RVL2 | | burst duratior | burst duration (ms) |
| mouse#1 | P3 | 918.57672 | 1124.54697 |
| mouse#2 | P3 | 789.323055 | 1064.27221 |
| mouse#3 | P3 | 1020.04401 | 995.2616 |
| mouse#4 | P4 | 1251.2512 | 1300.39965 |
| mouse#5 | P3 | 898.883333 | 1293.44526 |
| mouse#6 | P4 | 652.527778 | 833.083279 |
| mouse#7 | P3 | 1202.5004 | 1353.12616 |
| | MEAN | 961.9 | 1138 |
| | SEM | 81.11 | 71.68 |

Duration
Wilcoxon matched-pairs signed rank test

| | |
|---|---|
| P value | 0.0313 |
| P value summ | * |

| | | Cocktail | +Barium |
|---|---|---|---|
| RVL2 | | | |
| mouse#1 | P3 | burst frequen | burst frequency (Hz) |
| mouse#2 | P3 | 0.73618936 | 0.6727479 |
| mouse#3 | P3 | 0.82401229 | 0.67752055 |
| mouse#4 | P4 | 0.65755733 | 0.70548081 |
| mouse#5 | P3 | 0.53359404 | 0.47861621 |
| mouse#6 | P4 | 0.66645228 | 0.56636262 |
| mouse#7 | P3 | 1.03682599 | 0.83707864 |
| | | 0.61002673 | 0.54174776 |
| | MEAN | 0.7235 | 0.6257 |
| | SEM | 0.06267 | 0.04597 |

Frequency
Wilcoxon matched-pairs signed rank test

| | |
|---|---|
| P value | 0.0313 |
| P value summary | * |

**Fig. 5F** Data Source File

| | Cross-correlogram coefficient factor | | |
|---|---|---|---|
| | RVL2l-RVL2r | Oscill. aCSF | +Barium |
| mouse#1 | #190319 | -0.68799543 | -0.42537308 |
| mouse#2 | #200319 | -0.32078913 | -0.12526347 |
| mouse#3 | #210319 | -0.8402155 | -0.62014 |
| mouse#4 | #250319 | -0.7596255 | -0.57956 |
| mouse#5 | #260319 | -0.6034956 | -0.35937051 |
| mouse#6 | #171221 | -0.795644 | -0.659563 |
| | MEAN | -0.668 | -0.4615 |
| | SEM | 0.07734 | 0.08228 |

| | Cross-correlogram coefficient factor | | |
|---|---|---|---|
| | RVL2l-RVL2r | Cocktail | +Barium |
| mouse#1 | #060219 | -0.771666667 | -0.656666667 |
| mouse#2 | #180319 | -0.80224 | -0.613833333 |
| mouse#3 | #190319 | -0.823002 | -0.600666667 |
| mouse#4 | #200319 | -0.808666667 | -0.6758 |
| mouse#5 | #210319 | -0.7655584 | -0.638445 |
| mouse#6 | #250319 | -0.61045 | -0.601266667 |
| mouse#7 | #260319 | -0.7642 | -0.728166667 |
| | MEAN | -0.7637 | -0.645 |
| | SEM | 0.02698 | 0.01752 |

**Fig. 6C**

Data Source File

| | Ctrl-ShRNA | | Kir4.1-ShRNA |
|---|---|---|---|
| mouse#1 | 0.889176169 | mouse#1 | 0.720688399 |
| mouse#2 | 1.110823831 | mouse#2 | 0.633681985 |
| mouse#3 | 1 | mouse#3 | 0.552888474 |
| mouse#4 | 0.969968611 | mouse#4 | 0.591369071 |
| mouse#5 | 1.030031389 | mouse#5 | 0.642108793 |
| MEAN | 1 | MEAN | 0.6281 |
| SEM | 0.03631 | SEM | 0.02811 |

| | |
|---|---|
| Mann Whitney test | |
| P value | 0.0079 |
| P value summary | ** |

Fig. 6F (a)

### Data Source File

| | Ctrl-ShRNA | age | Vrest (mV) | Rin (Mohm) | Coktail-induced depol. (mV) |
|---|---|---|---|---|---|
| mouse#1 | IN1 | P10 | -72.11 | 142 | 20.69 |
| mouse#1 | IN2 | P10 | -64.87 | 567 | 4.73 |
| mouse#1 | IN3 | P10 | -68.35 | 793 | 18.59 |
| mouse#2 | IN4 | P12 | -60.39 | 986 | 5 |
| mouse#2 | IN5 | P12 | -68.14 | 328 | 16.82 |
| mouse#2 | IN6 | P12 | -51.28 | 249 | 13.15 |
| mouse#2 | IN7 | P12 | -57.97 | 575 | 12.97 |
| mouse#2 | IN8 | P12 | -60.79 | 515 | 12 |
| mouse#3 | IN9 | P12 | -60.48 | 380 | |
| mouse#3 | IN10 | P12 | -64.54 | 371 | |
| mouse#3 | IN11 | P12 | -61.6 | 467 | |
| mouse#3 | IN12 | P12 | -70.3 | 469 | |
| mouse#3 | IN13 | P12 | -60.77 | 656 | |
| mouse#3 | IN14 | P12 | -60.47 | 294 | |
| mouse#4 | IN15 | P13 | -63.15 | 272 | |
| mouse#4 | IN16 | P13 | -74.44 | 296 | |
| mouse#4 | IN17 | P13 | -60.07 | 520 | |
| mouse#4 | IN18 | P13 | -59.5 | 362 | |
| | | MEAN | -63.29 | 458 | 12.99375 |
| | | SEM | 1.29 | 47.71 | 1.33 |

**Fig. 6F (b)**

| | Kir4.1-ShRNA | age | Vrest (mV) | Rin (Mohm) | Coktail-induced depol. (mV) |
|---|---|---|---|---|---|
| mouse#1 | IN1 | P11 | -51.98 | 829 | 23.22 |
| mouse#1 | IN2 | P11 | -60.78 | 950.97083 | 22.8 |
| mouse#1 | IN3 | P11 | -52.48 | 663.519841 | 10.45 |
| mouse#1 | IN4 | P11 | -54.2 | 882.696457 | 16.11 |
| mouse#1 | IN5 | P11 | -67.17 | 421 | 23.2 |
| mouse#2 | IN6 | P13 | -50.44 | 319.706145 | 15.63 |
| mouse#2 | IN7 | P13 | -55.21 | 210.77535 | 6.74 |
| mouse#2 | IN8 | P13 | -50.53 | 982.566631 | 16.94 |
| mouse#3 | IN9 | P13 | -62 | 824.1 | |
| mouse#3 | IN10 | P13 | -63 | 300.2 | |
| mouse#3 | IN11 | P13 | -57 | 688.4 | |
| mouse#3 | IN12 | P13 | -57.1 | 609.2 | |
| mouse#3 | IN13 | P13 | -58.4 | 761.2 | |
| mouse#3 | IN14 | P13 | -59.2 | 684 | |
| mouse#4 | IN15 | P14 | -64.7 | 320 | |
| mouse#4 | IN16 | P14 | -59.6 | 802 | |
| mouse#4 | IN17 | P14 | -56 | 109.75 | |
| mouse#4 | IN18 | P14 | -63.9 | 506.25 | |
| mouse#4 | IN19 | P14 | -57.2 | 289.4 | |
| mouse#4 | IN20 | P14 | -59.3 | 806.8 | |
| mouse#4 | IN21 | P14 | -52.7 | 550.6 | |
| mouse#5 | IN22 | P12 | -55.9 | 653 | |
| mouse#5 | IN23 | P12 | -54.3 | 349.8 | |
| mouse#5 | IN24 | P12 | -56.1 | 512.6 | |
| mouse#5 | IN25 | P12 | -60.2 | 728.3 | |
| mouse#5 | IN26 | P12 | -57.9 | 427.3 | |
| mouse#5 | IN27 | P12 | -64.8 | 341.2 | |
| mouse#5 | IN28 | P12 | -68.9 | 608 | |
| mouse#5 | IN29 | P12 | -58.2 | 352.8 | |
| mouse#5 | IN30 | P12 | -57.1 | 277 | |
| | | MEAN | -58.2096667 | 558.737842 | 16.88625 |
| | | SEM | 0.86 | 43.68 | 1.11 |

Vrest

| | |
|---|---|
| Mann Whitney test | |
| P value | 0.001 |
| P value summary | ** |

Rin

| | |
|---|---|
| Mann Whitney test | |
| P value | 0.1582 |
| P value summary | ns |

Cocktail-induced depol (mV)

| | |
|---|---|
| Mann Whitney test | |
| P value | 0.1304 |
| P value summary | ns |

**Fig. 6I (a)** Data Source File

| | Ctrl-ShRNA | age | Oscill. induced by K+6mM Ca2+0.9mM |
|---|---|---|---|
| mouse#1 | IN1 | P12 | + |
| mouse#1 | IN2 | P12 | + |
| mouse#1 | IN3 | P12 | + |
| mouse#1 | IN4 | P12 | - |
| mouse#1 | IN5 | P12 | - |
| mouse#1 | IN6 | P12 | + |
| mouse#2 | IN7 | P13 | + |
| mouse#2 | IN8 | P13 | + |
| mouse#2 | IN9 | P13 | + |
| mouse#2 | IN10 | P13 | + |

| Ctrl-ShRNA | | |
|---|---|---|
| | Oscillating INs (nb) | 8 |
| | Not Oscillating INs (nb) | 2 |
| | Oscillating INs (%) | 80 |
| | Not Oscillating INs (%) | 20 |

| | Ctrl-ShRNA | Ampl Oscill. (mV) | Duration Oscill. (s) | Frequ. Oscill. (Hz) |
|---|---|---|---|---|
| mouse#1 | IN1 | 5.2 | 0.15 | 1.05 |
| mouse#1 | IN2 | 10.5 | 1.36333333 | 0.36666667 |
| mouse#1 | IN3 | 7 | 0.12 | 1.1 |
| mouse#1 | IN6 | 5.722857143 | 0.44714286 | 0.48 |
| mouse#2 | IN7 | 7.621666667 | 0.55083333 | 0.8 |
| mouse#2 | IN8 | 5 | 1.14 | 0.1 |
| mouse#2 | IN9 | 11 | 0.58 | 0.75 |
| mouse#2 | IN10 | 15.5 | 3 | 0.26666667 |
| | MEAN | 8.443065476 | 0.91891369 | 0.61416667 |
| | SEM | 1.289366259 | 0.33486972 | 0.12979035 |

**Fig. 6I (b)**

Data Source File

| | Kir4.1-ShRNA | age | Oscill. induced by K+6mM Ca2+0.9mM |
|---|---|---|---|
| mouse#1 | IN1 | P12 | - |
| mouse#1 | IN2 | P12 | - |
| mouse#1 | IN3 | P12 | - |
| mouse#1 | IN4 | P12 | - |
| mouse#1 | IN5 | P12 | - |
| mouse#1 | IN6 | P12 | + |
| mouse#1 | IN7 | P12 | - |
| mouse#1 | IN8 | P12 | + |
| mouse#1 | IN9 | P12 | - |
| mouse#2 | IN10 | P13 | - |
| mouse#2 | IN11 | P13 | - |
| mouse#2 | IN12 | P13 | - |
| mouse#2 | IN13 | P13 | + |
| mouse#3 | IN14 | P14 | + |
| mouse#3 | IN15 | P14 | - |
| mouse#3 | IN16 | P14 | - |
| mouse#3 | IN17 | P14 | - |

| Kir4.1-ShRNA | | |
|---|---|---|
| | Oscillating INs (nb) Not | 4 |
| | Oscillating INs (nb) | 13 |
| | Oscillating INs (%) | 23.5294118 |
| | Not Oscillating INs (%) | 76.4705882 |

| | Kir4.1-ShRNA | Ampl Oscill. (mV) | Duration Oscill. (s) | Frequ. Oscill. (Hz) |
|---|---|---|---|---|
| mouse#1 | IN6 | 5.065 | 0.51 | 0.8 |
| mouse#1 | IN8 | 9.47875 | 1.760375 | 0.12 |
| mouse#2 | IN13 | 15.412 | 1.966 | 0.125 |
| mouse#3 | IN14 | 7.903333333 | 1.29 | 0.5 |
| | MEAN | 9.464770833 | 1.38159375 | 0.38625 |
| | SEM | 2.182616943 | 0.32314483 | 0.16413124 |

| | |
|---|---|
| Test | Fisher's exact test |
| P value | <0.0001 |
| P value summary | ***** |

**Fig. 6J (a)**

Data Source File

| | Ctrl-ShRNA | age | Oscillations induced by TTX+Cocktail |
|---|---|---|---|
| mouse#3 | IN1 | P10 | + |
| mouse#3 | IN2 | P10 | + |
| mouse#3 | IN3 | P10 | + |
| mouse#3 | IN4 | P12 | - |
| mouse#3 | IN5 | P12 | - |
| mouse#4 | IN6 | P12 | + |
| mouse#4 | IN7 | P12 | + |
| mouse#4 | IN8 | P12 | + |

| Ctrl-ShRNA | | |
|---|---|---|
| | Oscillating INs (nb) | 6 |
| | Not Oscillating INs (nb) | 8 |
| | Oscillating INs (%) | 75 |
| | Not Oscillating (%) | 25 |

| | Ctrl-ShRNA | Ampl (mV) | Duration (s) | Frequency (Hz) |
|---|---|---|---|---|
| mouse#3 | IN1 | 6.68153846 | 0.917692308 | 0.61904762 |
| mouse#3 | IN2 | 10.3547619 | 2.484285714 | 0.21 |
| mouse#3 | IN3 | 10.2031111 | 0.626888889 | 1.5 |
| mouse#4 | IN6 | 4.67333333 | 0.931111111 | 0.9 |
| mouse#4 | IN7 | 2.408 | 1.05 | 0.33333 |
| mouse#4 | IN8 | 5.30833333 | 0.616333333 | 1.5 |
| | MEAN | 6.60484636 | 1.104385226 | 0.8437296 |
| | SEM | 1.29151675 | 0.285082367 | 0.22928519 |

| | Kir4.1-ShRNA | age | Oscillations induced by TTX+Cocktail |
|---|---|---|---|
| mouse#4 | IN1 | P11 | + |
| mouse#4 | IN2 | P11 | + |
| mouse#4 | IN3 | P11 | - |
| mouse#4 | IN4 | P11 | - |
| mouse#4 | IN5 | P11 | - |
| mouse#5 | IN6 | P13 | - |
| mouse#5 | IN7 | P13 | - |
| mouse#5 | IN8 | P13 | - |
| mouse#6 | IN9 | P13 | - |
| mouse#6 | IN10 | P13 | - |
| mouse#7 | IN11 | P14 | - |
| mouse#7 | IN12 | P14 | - |
| mouse#7 | IN13 | P14 | + |

| Kir4.1-ShRNA | | |
|---|---|---|
| | Oscillating INs (nb) | 3 |
| | Not Oscillating INs (nb) | 13 |
| | Oscillating INs (%) | 76.92307692 |
| | Not Oscillating (%) | 23.07692308 |

**Fig. 6J (b)**

Data Source File

| | Kir4.1-ShRNA | Ampl (mV) | Duration (s) | Frequency (Hz) |
|---|---|---|---|---|
| mouse#4 | IN1 | 22.8720833 | 2.082916667 | 0.4 |
| mouse#4 | IN2 | 4.89363636 | 1.483636364 | 0.275 |
| mouse#7 | IN13 | 6.7 | 1.18604651 | 0.2934 |
| | MEAN | 11.4885732 | 1.584199847 | 0.3228 |
| | SEM | 5.71559166 | 0.263741528 | 0.03896374 |

| | |
|---|---|
| Test | Fisher's exact test |
| P value | <0.0001 |
| P value summary | ***** |
| One- or two-sided | Two-sided |
| Statistically signiicant (P < 0.05)? | Yes |

**Fig. 7D (a)**

Data Source File

| | Ctrl-ShRNA | | Kir4.1-ShRNA |
|---|---|---|---|
| | Stand (s) | | Stand (s) |
| mouse#1 P14 | 0.13183605 | mouse#1 P14 | 0.13753184 |
| mouse#2 P14 | 0.12255223 | mouse#2 P14 | 0.1324181 |
| mouse#3 P14 | 0.11113208 | mouse#3 P14 | 0.10555308 |
| mouse#4 P14 | 0.10609027 | mouse#4 P14 | 0.212185 |
| mouse#5 P14 | 0.13111559 | mouse#5 P14 | 0.22234056 |
| mouse#6 P14 | 0.17072504 | mouse#6 P14 | 0.1254767 |
| mouse#7 P14 | 0.11339491 | mouse#7 P14 | 0.26274614 |
| mouse#8 P14 | 0.16089835 | mouse#8 P14 | 0.13915135 |
| mouse#9 P14 | 0.08315397 | mouse#9 P14 | 0.16973554 |
| mouse#10 P18 | 0.13874314 | mouse#10 P18 | 0.14976349 |
| mouse#11 P18 | 0.1662625 | mouse#11 P18 | 0.17652985 |
| mouse#12 P18 | 0.10471217 | mouse#12 P18 | 0.11026923 |
| mouse#13 P18 | 0.12872248 | mouse#13 P18 | 0.13393126 |
| mouse#14 P18 | 0.11784834 | mouse#14 P18 | 0.13854294 |
| MEAN | 0.1277 | MEAN | 0.1583 |
| SEM | 0.006702 | SEM | 0.0122 |

| | |
|---|---|
| Stand (s) | |
| Mann Whitney test | |
| P value | 0.0395 |
| P value summary | * |

| | Ctrl-ShRNA | | Kir4.1-ShRNA |
|---|---|---|---|
| | Stand (%) | | Stand (%) |
| mouse#1 P14 | 58.6698596 | mouse#1 P14 | 61.2046088 |
| mouse#2 P14 | 54.5383634 | mouse#2 P14 | 58.9288863 |
| mouse#3 P14 | 49.4561495 | mouse#3 P14 | 46.9733766 |
| mouse#4 P14 | 47.2124378 | mouse#4 P14 | 94.4268617 |
| mouse#5 P14 | 58.3492409 | mouse#5 P14 | 98.9463036 |
| mouse#6 P14 | 75.976294 | mouse#6 P14 | 55.8398159 |
| mouse#7 P14 | 50.4631594 | mouse#7 P14 | 116.927647 |
| mouse#8 P14 | 71.6032033 | mouse#8 P14 | 61.9253257 |
| mouse#9 P14 | 37.0052934 | mouse#9 P14 | 75.5359452 |
| mouse#10 P18 | 61.7436618 | mouse#10 P18 | 66.6479548 |
| mouse#11 P18 | 73.9903645 | mouse#11 P18 | 78.5595574 |
| mouse#12 P18 | 46.5991551 | mouse#12 P18 | 49.0721661 |
| mouse#13 P18 | 57.2842536 | mouse#13 P18 | 59.6022758 |
| mouse#14 P18 | 52.4450306 | mouse#14 P18 | 61.6545691 |
| MEAN | 56.8097476 | MEAN | 70.4460925 |
| SEM | 2.98247688 | SEM | 5.42720316 |

| | |
|---|---|
| Stand (%) | |
| Mann Whitney test | |
| P value | 0.0395 |
| P value summary | * |

**Fig. 7D (b)**

Data Source File

| | Ctrl-ShRNA | | Kir4.1-ShRNA |
|---|---|---|---|
| | Swing (s) | | Swing (s) |
| mouse#1 P14 | 0.09159867 | mouse#1 P14 | 0.10406992 |
| mouse#2 P14 | 0.07269519 | mouse#2 P14 | 0.11514976 |
| mouse#3 P14 | 0.06705247 | mouse#3 P14 | 0.32321963 |
| mouse#4 P14 | 0.15065118 | mouse#4 P14 | 0.10309257 |
| mouse#5 P14 | 0.08588921 | mouse#5 P14 | 0.10084047 |
| mouse#6 P14 | 0.10670315 | mouse#6 P14 | 0.102487 |
| mouse#7 P14 | 0.08478785 | mouse#7 P14 | 0.12164805 |
| mouse#8 P14 | 0.14046662 | mouse#8 P14 | 0.11239732 |
| mouse#9 P14 | 0.0976225 | mouse#9 P14 | 0.12730166 |
| mouse#10 P18 | 0.11174753 | mouse#10 P18 | 0.11881237 |
| mouse#11 P18 | 0.11195052 | mouse#11 P18 | 0.12175196 |
| mouse#12 P18 | 0.10260458 | mouse#12 P18 | 0.1683842 |
| mouse#13 P18 | 0.0690561 | mouse#13 P18 | 0.10687886 |
| mouse#14 P18 | 0.06590361 | mouse#14 P18 | 0.12356532 |
| MEAN | 0.09705 | MEAN | 0.1321 |
| SEM | 0.006982 | SEM | 0.01539 |

| | |
|---|---|
| Swing (s) | |
| Mann Whitney test | |
| P value | 0.0067 |
| P value summary | ** |

| | Ctrl-ShRNA | | Kir4.1-ShRNA |
|---|---|---|---|
| | Swing (%) | | Swing (%) |
| mouse#1 P14 | 40.7633662 | mouse#1 P14 | 46.3133391 |
| mouse#2 P14 | 32.3509135 | mouse#2 P14 | 51.2441047 |
| mouse#3 P14 | 29.8397825 | mouse#3 P14 | 143.839645 |
| mouse#4 P14 | 67.0429955 | mouse#4 P14 | 45.8783974 |
| mouse#5 P14 | 38.2225345 | mouse#5 P14 | 44.8761648 |
| mouse#6 P14 | 47.4851827 | mouse#6 P14 | 45.6089058 |
| mouse#7 P14 | 37.7324057 | mouse#7 P14 | 54.1359827 |
| mouse#8 P14 | 62.5106486 | mouse#8 P14 | 50.0192101 |
| mouse#9 P14 | 43.4440993 | mouse#9 P14 | 56.65196 |
| mouse#10 P18 | 49.7300396 | mouse#10 P18 | 52.8740445 |
| mouse#11 P18 | 49.8203745 | mouse#11 P18 | 54.1822248 |
| mouse#12 P18 | 45.6612314 | mouse#12 P18 | 74.9345685 |
| mouse#13 P18 | 30.7314417 | mouse#13 P18 | 47.5633774 |
| mouse#14 P18 | 29.3285171 | mouse#14 P18 | 54.9892088 |
| MEAN | 43.1902524 | MEAN | 58.7936524 |
| SEM | 3.10708785 | SEM | 6.85028801 |

| | |
|---|---|
| Swing (%) | |
| Mann Whitney test | |
| P value | 0.0067 |
| P value summary | ** |

**Fig. 7D (c)**

Data Source File

| | Ctrl-ShRNA Regularity Index (%) | | Kir4.1-ShRNA Regularity Index (%) |
|---|---|---|---|
| mouse#1 P14 | 100 | mouse#1 P14 | 85.3294853 |
| mouse#2 P14 | 100 | mouse#2 P14 | 89.1812865 |
| mouse#3 P14 | 100 | mouse#3 P14 | 64.7826087 |
| mouse#4 P14 | 100 | mouse#4 P14 | 78.5714286 |
| mouse#5 P14 | 100 | mouse#5 P14 | 86.9565217 |
| mouse#6 P14 | 100 | mouse#6 P14 | 79.9422799 |
| mouse#7 P14 | 95 | mouse#7 P14 | 80 |
| mouse#8 P14 | 91.5384615 | mouse#8 P14 | 95.2380952 |
| mouse#9 P14 | 100 | mouse#9 P14 | 100 |
| mouse#10 P18 | 95.2380952 | mouse#10 P18 | 100 |
| mouse#11 P18 | 100 | mouse#11 P18 | 92.6739927 |
| mouse#12 P18 | 94.4444444 | mouse#12 P18 | 80 |
| mouse#13 P18 | 100 | mouse#13 P18 | 84.1750842 |
| mouse#14 P18 | 100 | mouse#14 P18 | 86.6666667 |
| MEAN | 98.3 | MEAN | 85.97 |
| SEM | 0.7765 | SEM | 2.515 |

| Regularity Index (%) | |
|---|---|
| Mann Whitney test | |
| P value | 0.0001 |
| P value summary | *** |

| | Ctrl-ShRNA Run Avg Speed (cm/s) | | Kir4.1-ShRNA Run Avg Speed (cm/s) |
|---|---|---|---|
| mouse#1 P14 | 27.905 | mouse#1 P14 | 19.34 |
| mouse#2 P14 | 22.545 | mouse#2 P14 | 19.0766667 |
| mouse#3 P14 | 21.145 | mouse#3 P14 | 8.135 |
| mouse#4 P14 | 17.5 | mouse#4 P14 | 9.37 |
| mouse#5 P14 | 17.325 | mouse#5 P14 | 4.23 |
| mouse#6 P14 | 24.14 | mouse#6 P14 | 14.8933333 |
| mouse#7 P14 | 15.45 | mouse#7 P14 | 6.84 |
| mouse#8 P14 | 18.05 | mouse#8 P14 | 14.97 |
| mouse#9 P14 | 45.17 | mouse#9 P14 | 21.29 |
| mouse#10 P18 | 28.69 | mouse#10 P18 | 27.18 |
| mouse#11 P18 | 23.5366667 | mouse#11 P18 | 23.5233333 |
| mouse#12 P18 | 29.155 | mouse#12 P18 | 22.3933333 |
| mouse#13 P18 | 22.4 | mouse#13 P18 | 23.4533333 |
| mouse#14 P18 | 40.22 | mouse#14 P18 | 21.96 |
| MEAN | 25.23 | MEAN | 16.9 |
| SEM | 2.299 | SEM | 1.935 |

| Run Avg Speed (cm/s) | |
|---|---|
| Mann Whitney test | |
| P value | 0.0162 |
| P value summary | * |

**Fig. 7D (d)**

Data Source File

| | Ctrl-ShRNA Base of Support (cm) | | Kir4.1-ShRNA Base of Support (cm) |
|---|---|---|---|
| mouse#1 P14 | 1.78082437 | mouse#1 P14 | 1.70445469 |
| mouse#2 P14 | 1.72634409 | mouse#2 P14 | 1.97646697 |
| mouse#3 P14 | 1.80148148 | mouse#3 P14 | 1.84474142 |
| mouse#4 P14 | 2.04285714 | mouse#4 P14 | 2.02988662 |
| mouse#5 P14 | 2.03049645 | mouse#5 P14 | 2.08888889 |
| mouse#6 P14 | 2.00380952 | mouse#6 P14 | 1.91451751 |
| mouse#7 P14 | 1.6 | mouse#7 P14 | 1.61579254 |
| mouse#8 P14 | 1.4533517 | mouse#8 P14 | 1.9489899 |
| mouse#9 P14 | 3.01742424 | mouse#9 P14 | 2.41287879 |
| mouse#10 P18 | 2.0007732 | mouse#10 P18 | 2.60290404 |
| mouse#11 P18 | 1.64187285 | mouse#11 P18 | 1.89693585 |
| mouse#12 P18 | 1.7830756 | mouse#12 P18 | 2.2102606 |
| mouse#13 P18 | 1.90000006 | mouse#13 P18 | 1.86426638 |
| mouse#14 P18 | 1.75589226 | mouse#14 P18 | 1.90721649 |
| MEAN | 1.896 | MEAN | 2.001 |
| SEM | 0.0981 | SEM | 0.07009 |

| Base of Support (cm) | |
|---|---|
| Mann Whitney test | |
| P value | 0.1781 |
| P value summary | ns |

**Fig. 7E (a)** Data Source File

| Strain | Manip | Trial 1 (s) | Trial 2 (s) | Trial 3 (s) | Mean 3 Trials (s) |
|---|---|---|---|---|---|
| mouse#1 | Ctrl-ShRNA | 108 | 97 | 212 | 139 |
| mouse#2 | Ctrl-ShRNA | 290 | 280 | 270 | 280 |
| mouse#1 | Kir4.1-ShRNA | 207 | 238 | 147 | 197.3333333 |
| mouse#2 | Kir4.1-ShRNA | 147 | 184 | 216 | 182.3333333 |
| mouse#3 | Ctrl-ShRNA | 151 | 125 | 233 | 169.6666667 |
| mouse#4 | Ctrl-ShRNA | 112 | 145 | 202 | 153 |
| mouse#3 | Kir4.1-ShRNA | 112 | 50 | 140 | 100.6666667 |
| mouse#4 | Kir4.1-ShRNA | 138 | 117 | 138 | 131 |
| mouse#5 | Ctrl-ShRNA | 120 | 135 | 154 | 136.3333333 |
| mouse#6 | Ctrl-ShRNA | 90 | 100 | 137 | 109 |
| mouse#5 | Kir4.1-ShRNA | 81 | 90 | 62 | 77.66666667 |
| mouse#6 | Kir4.1-ShRNA | 30 | 74 | 54 | 52.66666667 |
| mouse#7 | Ctrl-ShRNA | 93 | 110 | 133 | 112 |
| mouse#8 | Ctrl-ShRNA | 89 | 98 | 116 | 101 |
| mouse#7 | Kir4.1-ShRNA | 52 | 78 | 65 | 65 |
| mouse#8 | Kir4.1-ShRNA | 28 | 53 | 63 | 48 |
| mouse#9 | Ctrl-ShRNA | 48 | 51 | 60 | 53 |
| mouse#10 | Ctrl-ShRNA | 66 | 100 | 126 | 97.33333333 |
| mouse#9 | Kir4.1-ShRNA | 51 | 61 | 88 | 66.66666667 |
| mouse#10 | Kir4.1-ShRNA | 12 | 15 | 71 | 32.66666667 |
| mouse#11 | Ctrl-ShRNA | 128 | 143 | 147 | 139.3333333 |
| mouse#12 | Ctrl-ShRNA | 135 | 147 | 161 | 147.6666667 |
| mouse#11 | Kir4.1-ShRNA | 30 | 125 | 133 | 96 |
| mouse#12 | Kir4.1-ShRNA | 102 | 91 | 100 | 97.66666667 |
| mouse#13 | Ctrl-ShRNA | 148 | 130 | 152 | 143.3333333 |
| mouse#14 | Ctrl-ShRNA | 137 | 161 | 162 | 153.3333333 |
| mouse#13 | Kir4.1-ShRNA | 116 | 83 | 118 | 105.6666667 |
| mouse#14 | Kir4.1-ShRNA | 56 | 38 | 81 | 58.33333333 |

| | | Trial 1 (s) | Trial 2 (s) | Trial 3 (s) | Mean 3 Trials (s) |
|---|---|---|---|---|---|
| mouse#1 | Ctrl-ShRNA | 108 | 97 | 212 | 139 |
| mouse#2 | Ctrl-ShRNA | 290 | 280 | 270 | 280 |
| mouse#3 | Ctrl-ShRNA | 151 | 125 | 233 | 169.6666667 |
| mouse#4 | Ctrl-ShRNA | 112 | 145 | 202 | 153 |
| mouse#5 | Ctrl-ShRNA | 120 | 135 | 154 | 136.3333333 |
| mouse#6 | Ctrl-ShRNA | 90 | 100 | 137 | 109 |
| mouse#7 | Ctrl-ShRNA | 93 | 110 | 133 | 112 |
| mouse#8 | Ctrl-ShRNA | 89 | 98 | 116 | 101 |
| mouse#9 | Ctrl-ShRNA | 48 | 51 | 60 | 53 |
| mouse#10 | Ctrl-ShRNA | 66 | 100 | 126 | 97.33333333 |
| mouse#11 | Ctrl-ShRNA | 128 | 143 | 147 | 139.3333333 |
| mouse#12 | Ctrl-ShRNA | 135 | 147 | 161 | 147.6666667 |
| mouse#13 | Ctrl-ShRNA | 148 | 130 | 152 | 143.3333333 |
| mouse#14 | Ctrl-ShRNA | 137 | 161 | 162 | 153.3333333 |
| | | | | MEAN | 138.1428571 |
| | | | | SEM | 13.5709016 |

**Fig. 7E (b)** Data Source File

| | | Trial 1 (s) | Trial 2 (s) | Trial 3 (s) | Mean 3 Trials (s) |
|---|---|---|---|---|---|
| mouse#1 | Kir4.1-ShRNA | 207 | 238 | 147 | 197.3333333 |
| mouse#2 | Kir4.1-ShRNA | 147 | 184 | 216 | 182.3333333 |
| mouse#3 | Kir4.1-ShRNA | 112 | 50 | 140 | 100.6666667 |
| mouse#4 | Kir4.1-ShRNA | 138 | 117 | 138 | 131 |
| mouse#5 | Kir4.1-ShRNA | 81 | 90 | 62 | 77.66666667 |
| mouse#6 | Kir4.1-ShRNA | 30 | 74 | 54 | 52.66666667 |
| mouse#7 | Kir4.1-ShRNA | 52 | 78 | 65 | 65 |
| mouse#8 | Kir4.1-ShRNA | 28 | 53 | 63 | 48 |
| mouse#9 | Kir4.1-ShRNA | 51 | 61 | 88 | 66.66666667 |
| mouse#10 | Kir4.1-ShRNA | 12 | 15 | 71 | 32.66666667 |
| mouse#11 | Kir4.1-ShRNA | 30 | 125 | 133 | 96 |
| mouse#12 | Kir4.1-ShRNA | 102 | 91 | 100 | 97.66666667 |
| mouse#13 | Kir4.1-ShRNA | 116 | 83 | 118 | 105.6666667 |
| mouse#14 | Kir4.1-ShRNA | 56 | 38 | 81 | 58.33333333 |
| | | | | MEAN | 93.69047619 |
| | | | | SEM | 13.00030357 |

| **Šídák's multi** | Summary | Adjusted P Value |
|---|---|---|
| ShLuci vs ShKir | | |
| Trial1 | ns | 0.1594 |
| Trial2 | ns | 0.1946 |
| Trial3 | * | 0.0212 |

Mean latency
to fall (s)

| **Mann Whitney test** | |
|---|---|
| P value | 0.0106 |
| P value summ | * |

**Fig. 7F** Data Source File

| | Ctrl-ShRNA | Distance moved center-point Total cm | Velocity center-point Mean cm/s | Movement Moving / center-point Cumulative Duration % | Movement Not Moving / center-point Cumulative Duration % |
|---|---|---|---|---|---|
| mouse#1 | P14 | 716 | 7.962621 | 87.31852 | 12.6814815 |
| mouse#2 | P14 | 466.6197 | 5.195379 | 73.82082 | 26.1791753 |
| mouse#3 | P14 | 667.5029 | 7.77132 | 78.24473 | 21.7552718 |
| mouse#4 | P14 | 642.088333 | 7.44402667 | 89.8289554 | 10.1710446 |
| mouse#5 | P14 | 619.640667 | 7.08528333 | 88.697231 | 11.302769 |
| mouse#6 | P14 | 554.7058 | 6.1689 | 80.87407 | 19.12593 |
| mouse#7 | P14 | 505.958 | 5.626755 | 77.82222 | 22.17778 |
| mouse#8 | P14 | 679.5493 | 7.565613 | 84.02267 | 15.97733 |
| mouse#9 | P14 | 526.232 | 5.852227 | 77.82222 | 22.17778 |
| | MEAN | 597.6 | 6.741 | 82.05 | 17.9498402 |
| | SEM | 29.01 | 0.3455 | 1.884 | 1.88442822 |

| | Kir4.1-ShRNA | Distance moved center-point Total cm | Velocity center-point Mean cm/s | Movement Moving / center-point Cumulative Duration % | Movement Not Moving / center-point Cumulative Duration % |
|---|---|---|---|---|---|
| mouse#1 | P14 | 566.5483 | 6.311365 | 80.54319 | 19.4568121 |
| mouse#2 | P14 | 285.5235 | 3.181282 | 45.21726 | 54.7827377 |
| mouse#3 | P14 | 301.9565 | 3.372984 | 62.65437 | 37.3456331 |
| mouse#4 | P14 | 542.989667 | 6.13686 | 82.9719964 | 17.0280036 |
| mouse#5 | P14 | 440.208333 | 5.03252 | 81.0712112 | 18.9287888 |
| mouse#6 | P14 | 469.4305 | 5.220647 | 72.71111 | 27.28889 |
| mouse#7 | P14 | 517.828 | 5.75876 | 79.15556 | 20.84444 |
| mouse#8 | P14 | 504.8126 | 5.624644 | 84.78984 | 15.21016 |
| mouse#9 | P14 | 549.778 | 6.129445 | 85.50336 | 14.49664 |
| | MEAN | 464.3 | 5.197 | 74.96 | 25.0424561 |
| SEM | | 34.84 | 0.3893 | 4.408 | 4.40835443 |

| | |
|---|---|
| Distance moved (cm) | |
| Mann Whitney test P value | 0.0188 |
| P value summary | * |
| Velocity (cm/s) | |
| Mann Whitney test P value | 0.0188 |
| P value summary | * |
| Mobility (%) | |
| Mann Whitney test P value | 0.3736 |
| P value summary | ns |

**Fig. S1B-C** Data Source File

| Astrocytes | Rest.Mb.Pot (mV) | Rest.Mb.Pot (mV) | Rest.Mb.Pot (mV) | Rest.Mb.Pot (mV) |
|---|---|---|---|---|
| $[K^+]_e$ mM | 3 | 6 | 9 | 12 |
| Astro1 | -84.4 | -76.4 | -67.4 | -59.4 |
| Astro2 | -86.9 | -74.9 | -66.9 | -61.9 |
| Astro3 | -87.4 | -77.4 | -75.4 | -66.4 |
| Astro4 | -87.4 | -84.4 | -73.9 | -65.4 |
| Astro5 | -87.9 | -76.4 | -72.9 | -66.4 |
| Astro6 | -88.4 | -74.4 | -68.4 | -58.4 |
| Astro7 | -85.6 | -70.1 | -60.6 | -58.6 |
| Astro8 | -80.4 | -70.4 | -65.4 | -58.4 |

| | | Delta Rest.Mb.Pot (mV) | Delta Rest.Mb.Pot (mV) | Delta Rest.Mb.Pot (mV) |
|---|---|---|---|---|
| Astro1 | 0 | 8 | 17 | 25 |
| Astro2 | 0 | 12 | 20 | 25 |
| Astro3 | 0 | 10 | 12 | 21 |
| Astro4 | 0 | 3 | 13.5 | 22 |
| Astro5 | 0 | 11.5 | 15 | 21.5 |
| Astro6 | 0 | 14 | 20 | 30 |
| Astro7 | 0 | 15.5 | 25 | 27 |
| Astro8 | 0 | 10 | 15 | 22 |

| | age | Vrest TTX | Vrest TTX+cocktail | delta Vm |
|---|---|---|---|---|
| mouse#1 | Astro1 | -84.92 | -76.42 | 8.5 |
| mouse#1 | Astro2 | -96.16 | -89.32 | 6.84 |
| mouse#1 | Astro3 | -86.46 | -81.86 | 4.6 |
| mouse#2 | Astro4 | -91.17 | -85.62 | 5.55 |
| mouse#3 | Astro5 | -91.32 | -67.21 | 24.11 |
| mouse#3 | Astro6 | -83.26 | -73.95 | 9.31 |
| mouse#4 | Astro7 | -89.47 | -82.2 | 7.27 |
| mouse#4 | Astro8 | -86.73 | -68 | 18.73 |
| | MEAN | -88.68625 | -78.0725 | 10.61375 |
| | SEM | 1.476 | 2.847 | 2.469161901 |

| Kruskal-Wallis test Multiple comparisons | Dunn's multiple comparisons test | Significant? | Summary | Adjusted P Value |
|---|---|---|---|---|
| | 3 vs. 6 | No | ns | 0.1974 |
| | 3 vs. 9 | Yes | ** | 0.0026 |
| | 3 vs. 12 | Yes | **** | <0.0001 |

**Fig. S2D (a)**

Astrocytes
$Ca^{2+}$transients

Data Source File

| aCSF Amplitude (df/f) | aCSF Duration (ms) | aCSF Frequency (Events nb/min) | TTX Amplitude (df/f) | TTX Duration (ms) | TTX Frequency (Events nb/min) |
|---|---|---|---|---|---|
| 0.18921709 | 1210.81452 | 3.33333333 | 0.21115006 | 1102.58108 | 3 |
| 0.23523119 | 1956.39492 | 3.33333333 | 0.22318505 | 1408.62573 | 1.33333333 |
| 0.27496242 | 1634.46739 | 2 | 0.31638458 | 1772.69696 | 2 |
| 0.17644326 | 1177.67581 | 3.66666667 | 0.2349851 | 1261.96118 | 2 |
| 0.26710048 | 1920.6321 | 1.66666667 | 0.25168255 | 1465.5141 | 3 |
| 0.20703131 | 826.668013 | 0.66666667 | 0.17412678 | 874.093755 | 0.33333333 |
| 0.20996703 | 3040.92943 | 2.33333333 | 0.38449924 | 1579.33497 | 3.33333333 |
| 0.29837739 | 1619.42009 | 3.33333333 | 0.45051927 | 1797.36832 | 1.66666667 |
| 0.20312656 | 960.873348 | 2.33333333 | 0.17777594 | 1369.35391 | 1.66666667 |
| 0.22206242 | 1425.48991 | 3.33333333 | 0.302056 | 1024.51625 | 4.33333333 |
| 0.29319243 | 871.152255 | 3.33333333 | 0.2554744 | 914.833901 | 2 |
| 0.19990877 | 1262.43441 | 1 | 0.2425289 | 1075.61263 | 2 |
| 0.39613683 | 904.040296 | 5 | 0.31885599 | 828.977308 | 3.66666667 |
| 0.19059307 | 1190.11122 | 1 | 0.16990323 | 798.2744 | 2 |
| 0.2296116 | 871.308021 | 8.33333333 | 0.20818247 | 1025.88354 | 6.33333333 |
| 0.24309587 | 2186.65573 | 0.33333333 | 0.31631746 | 1805.88568 | 0.66666667 |
| 0.23995208 | 980.636728 | 6.33333333 | 0.29930966 | 950.913543 | 7.33333333 |
| 0.34051397 | 2117.69741 | 3.33333333 | 0.28442313 | 2286.62803 | 3.66666667 |
| 0.1956287 | 1299.62504 | 2.33333333 | 0.26931223 | 983.952025 | 2.66666667 |
| 0.32524809 | 941.698253 | 3 | 0.28086276 | 874.575025 | 2.33333333 |
| 0.35789628 | 2675.02045 | 1.33333333 | 0.18482914 | 2516.92395 | 1 |
| 0.27803958 | 893.431537 | 4 | 0.21494077 | 1128.01585 | 3.33333333 |
| 0.32355634 | 1233.63703 | 5.66666667 | 0.35640611 | 1232.27165 | 4.66666667 |
| 0.18962074 | 1258.78686 | 1 | 0.17050432 | 1551.61631 | 1.66666667 |
| 0.22085465 | 1518.65328 | 1.66666667 | 0.21122109 | 1765.2881 | 2.66666667 |
| 0.26098795 | 953.366686 | 12.3333333 | 0.24997793 | 738.34471 | 9.33333333 |
| 0.21385838 | 1107.66708 | 4.66666667 | 0.25397867 | 1229.75066 | 3.33333333 |
| 0.3924914 | 915.94859 | 3.66666667 | 0.35641136 | 1024.80347 | 3.66666667 |
| 0.47582687 | 1648.9144 | 2.66666667 | 0.48109642 | 1625.05935 | 2 |
| 0.26363522 | 1247.48633 | 5.33333333 | 0.21756322 | 1203.30842 | 3.66666667 |
| 0.25991639 | 1393.2374 | 1.66666667 | 0.30536743 | 1265.98396 | 3.33333333 |
| 0.21596972 | 1350.46573 | 3 | 0.23373337 | 1253.0879 | 2.33333333 |
| 0.2541328 | 1093.42469 | 3 | 0.22862194 | 886.432119 | 3.33333333 |
| 0.29494669 | 1050.36068 | 7.33333333 | 0.23740622 | 975.492966 | 4.66666667 |
| 0.19837006 | 1182.88948 | 4 | 0.18712301 | 1078.55371 | 2.66666667 |
| 0.26997188 | 1010.06568 | 3.66666667 | 0.29475921 | 1182.26384 | 3.33333333 |
| 0.22994448 | 982.072011 | 4 | 0.33283379 | 1133.46349 | 5.66666667 |
| 0.35491144 | 1072.26853 | 3.33333333 | 0.3744445 | 1116.24964 | 3.66666667 |
| 0.32017189 | 1631.44763 | 3 | 0.22120974 | 1917.67964 | 3 |
| 0.253612 | 1127.26421 | 5.66666667 | 0.20889798 | 1131.60053 | 6.33333333 |
| 0.18190062 | 1073.68931 | 2 | 0.20748466 | 1728.11528 | 2 |
| 0.2649996 | 1456.13483 | 2.66666667 | 0.29519069 | 798.545828 | 2.33333333 |
| 0.22490271 | 2047.79295 | 2.33333333 | 0.21751846 | 1466.5653 | 2 |
| 0.48736332 | 1336.57343 | 4.33333333 | 0.2051127 | 1024.4032 | 2 |
| 0.21032039 | 1237.08986 | 2.66666667 | 0.229691 | 1639.37361 | 3 |

**Fig. S2D (b)** Data Source File

| | | | | | |
|---|---|---|---|---|---|
| 0.35236011 | 1685.92177 | 4.33333333 | 0.32266569 | 1775.15794 | 1.33333333 |
| 0.25299525 | 1261.5568 | 3.33333333 | 0.22719559 | 979.767231 | 3 |
| 0.2107656 | 1177.77459 | 3.33333333 | 0.23122627 | 1168.36737 | 3 |
| 0.25298496 | 1144.91762 | 6.33333333 | 0.29151406 | 1133.55411 | 4.66666667 |
| 0.27684503 | 1228.1734 | 2.33333333 | 0.2943782 | 1763.88858 | 2.33333333 |
| 0.28130669 | 1343.93498 | 6.33333333 | 0.41036353 | 1082.93153 | 4.66666667 |
| 0.26432641 | 1228.00876 | 1 | 0.63151889 | 1983.5219 | 2.66666667 |
| 0.25555896 | 990.855474 | 3.33333333 | 0.301811 | 828.379362 | 4.66666667 |
| 0.26596874 | 807.754193 | 4 | 0.52210397 | 843.424898 | 3.66666667 |
| 0.38153531 | 1631.29337 | 2.66666667 | 0.22113685 | 2124.98622 | 2.33333333 |
| 0.654651 | 8489.65636 | 0.33333333 | 0.40812796 | 2246.18043 | 3.33333333 |
| 0.61950143 | 1600.83854 | 1.66666667 | 0.31083646 | 1558.6531 | 3 |
| 0.19638254 | 1111.80203 | 0.33333333 | 0.43558871 | 1466.77285 | 0.66666667 |
| 0.28926586 | 1113.36992 | 1.33333333 | 0.25202322 | 1145.71797 | 2 |
| 0.24152964 | 1492.92119 | 4.66666667 | 0.27778979 | 1315.85337 | 3 |
| 0.33317559 | 810.795559 | 6 | 0.3623096 | 786.426027 | 4.66666667 |
| 0.39571068 | 1293.71802 | 5 | 0.21088214 | 1364.05185 | 2.66666667 |
| 0.4575267 | 1566.44313 | 1.33333333 | 0.22493717 | 989.277806 | 2 |
| 0.83270763 | 1005.68183 | 14.0487805 | 0.54520528 | 1081.62769 | 15.2 |
| 0.49860584 | 1343.83044 | 7.6097561 | 0.43389401 | 1471.81749 | 5.2 |
| 0.60723293 | 2116.74052 | 3.51219512 | 0.40258115 | 1597.82464 | 3.6 |
| 0.66420259 | 1012.87099 | 21.6585366 | 0.63135467 | 943.772892 | 22 |
| 0.4210124 | 1210.02456 | 16.3902439 | 0.40586797 | 1193.316 | 12.8 |
| 0.48195693 | 1159.20814 | 11.1219512 | 0.42115243 | 1161.84272 | 10.4 |
| 0.39714063 | 1050.13482 | 16.9756098 | 0.50033415 | 1068.85123 | 17.6 |
| 0.48116645 | 1035.63639 | 5.85365854 | 0.39318703 | 1158.28067 | 5.2 |
| 0.41147336 | 4593.88269 | 1.17073171 | 0.41304991 | 4946.70261 | 1.2 |
| 0.2529437 | 1685.92745 | 0.58536585 | 0.36147249 | 1026.70508 | 0.4 |
| 0.6459058 | 820.252284 | 14.6341463 | 0.50551369 | 780.96963 | 19.2 |
| 0.65668959 | 797.748278 | 17.5609756 | 0.50667538 | 784.486815 | 20 |
| 0.63667707 | 1067.86674 | 15.2195122 | 0.54183462 | 1251.56555 | 14.4 |
| 0.97117243 | 1462.58432 | 4.68292683 | 0.96434275 | 1654.07429 | 4.4 |
| 0.4916197 | 1037.53643 | 11.1219512 | 0.4770957 | 1036.42322 | 10 |
| 0.41738321 | 995.304522 | 14.0487805 | 0.45077403 | 959.730005 | 17.6 |
| 0.27717872 | 1670.98195 | 1.75609756 | 0.26097405 | 1815.1716 | 0.8 |
| 0.40981191 | 1010.22229 | 7.6097561 | 0.38970789 | 1170.0561 | 6 |
| 0.84295839 | 691.522513 | 22.2439024 | 0.6848252 | 818.537324 | 23.6 |
| 0.38203333 | 1433.51288 | 8.19512195 | 0.36595449 | 1434.91195 | 8 |
| 0.47733062 | 930.823873 | 6.43902439 | 0.54900056 | 983.617982 | 6.4 |
| 0.56776997 | 854.274751 | 22.8292683 | 0.45757428 | 1008.44681 | 18.8 |
| 0.97807023 | 1156.46541 | 2.92682927 | 0.83968098 | 1075.98559 | 2.8 |
| 0.44022262 | 1098.82202 | 14.0487805 | 0.61760619 | 974.726124 | 16.8 |
| 0.80742987 | 2057.86161 | 1.75609756 | 0.55116843 | 1812.66459 | 2.8 |
| 0.30223638 | 1399.07222 | 1.17073171 | 0.26732326 | 1835.31471 | 2.4 |
| 0.38159506 | 2020.02234 | 2.92682927 | 0.38956129 | 1673.98903 | 2.8 |
| 0.4200787 | 1294.14917 | 2.34146341 | 0.35456781 | 1601.54188 | 3.2 |
| 0.56713957 | 1124.14089 | 7.02439024 | 0.3830477 | 1045.54704 | 8 |
| 0.31021112 | 1317.30068 | 2.34146341 | 0.26772994 | 1258.06884 | 1.6 |
| 0.51886869 | 889.207118 | 14.6341463 | 0.35824475 | 1132.22419 | 11.6 |
| 0.44266685 | 1434.93717 | 3.51219512 | 0.31491122 | 1102.78329 | 2.8 |

**Fig. S2D (c)** Data Source File

| | | | | | |
|---|---|---|---|---|---|
| 0.83234142 | 2065.02098 | 4.09756098 | 0.68031281 | 2103.78983 | 4.8 |
| 0.2895792 | 1218.32498 | 1.17073171 | 0.3097487 | 1711.33498 | 2 |
| 0.2809676 | 1206.3543 | 4.09756098 | 0.66496736 | 1318.89503 | 2.8 |
| 0.27038804 | 1480.61375 | 1.17073171 | 0.32048463 | 1807.25185 | 2 |
| 0.4067383 | 1002.26714 | 18.7317073 | 0.40111541 | 1036.42375 | 16 |
| 0.61651651 | 937.889003 | 28.097561 | 0.70414203 | 1010.58209 | 24.8 |
| 0.31852218 | 1114.22526 | 2.22222222 | 0.33138153 | 1065.68676 | 0.75471698 |
| 0.45586252 | 879.828813 | 0.74074074 | 0.38153086 | 9078.68676 | 0.75471698 |
| 0.38513081 | 1140.61166 | 14.0740741 | 0.34266946 | 925.983684 | 11.6981132 |
| 0.31375954 | 995.82596 | 8.14814815 | 0.35633439 | 1217.21234 | 7.54716981 |
| 0.33250857 | 807.047736 | 2.22222222 | 0.27679075 | 1059.11962 | 2.64150943 |
| 0.47312641 | 1954.15849 | 2.22222222 | 0.60098501 | 1344.68686 | 1.50943396 |
| 0.40108934 | 1030.21643 | 14.8148148 | 0.36807599 | 1024.89539 | 14.3396226 |
| 0.34525507 | 1273.93668 | 11.8518519 | 0.3642665 | 1294.03051 | 8.67924528 |
| 0.32462916 | 1487.05061 | 4.44444444 | 0.36254171 | 1280.60445 | 3.39622642 |
| 0.28505142 | 943.973308 | 1.48148148 | 0.64369922 | 2111.90594 | 2.26415094 |
| 0.33706847 | 1259.59308 | 7.40740741 | 0.38355504 | 1146.36843 | 8.30188679 |
| 0.50505461 | 763.173095 | 29.6296296 | 0.6069559 | 896.209577 | 29.4339623 |
| 0.3679685 | 1064.21807 | 14.0740741 | 0.36335786 | 1039.39991 | 14.3396226 |
| 0.46442288 | 1017.18288 | 2.96296296 | 0.33373037 | 1703.52441 | 2.26415094 |
| 0.46442288 | 1017.18288 | 2.96296296 | 0.32276199 | 1968.48568 | 1.88679245 |
| 0.48170073 | 1010.86169 | 9.62962963 | 0.36517734 | 1090.57132 | 12.0754717 |
| 0.37507521 | 1901.08007 | 1.48148148 | 0.33784838 | 1306.51251 | 1.13207547 |
| 0.25418757 | 1931.03138 | 1.48148148 | 0.30832013 | 1934.50487 | 1.88679245 |
| 0.28849468 | 1450.7606 | 2.96296296 | 0.31489919 | 1188.16695 | 1.88679245 |
| 0.29346518 | 1196.47522 | 2.22222222 | 0.28331044 | 1105.93819 | 1.50943396 |
| 0.31957724 | 880.468127 | 2.22222222 | 0.30809247 | 1433.94591 | 1.88679245 |
| 0.34759847 | 972.341465 | 1.48148148 | 0.33793353 | 1161.67942 | 1.88679245 |
| 0.37311404 | 2665.93615 | 2.22222222 | 0.32133943 | 2492.82371 | 0.75471698 |
| 0.39707446 | 878.470719 | 6.66666667 | 0.39859634 | 973.163981 | 9.05660377 |
| 0.31698101 | 2160.37962 | 1.48148148 | 0.3807373 | 1081.79701 | 3.01886792 |
| 0.38476945 | 928.122873 | 5.92592593 | 0.39829208 | 867.422794 | 7.16981132 |
| 0.35255117 | 1085.12021 | 5.92592593 | 0.31308554 | 1085.54459 | 3.01886792 |
| 0.26783448 | 2404.15146 | 2.22222222 | 0.27719732 | 1415.28453 | 1.50943396 |
| 0.53380326 | 1121.93959 | 13.3333333 | 0.36163593 | 1169.77253 | 7.16981132 |
| 0.37999611 | 828.701623 | 2.22222222 | 0.58237932 | 914.217287 | 5.28301887 |
| 0.3940035 | 993.119922 | 2.22222222 | 0.36242998 | 1153.19171 | 1.50943396 |
| 0.36884226 | 3303.86122 | 0.74074074 | 0.46235466 | 1672.96102 | 0.37735849 |
| 0.2741027 | 1789.24417 | 2.96296296 | 0.28519524 | 1274.55001 | 3.01886792 |
| 0.25718271 | 2185.08318 | 0.74074074 | 0.31255863 | 1864.54921 | 0.37735849 |
| 0.32307639 | 1169.60507 | 4.44444444 | 0.28200858 | 1038.65801 | 2.64150943 |
| 0.41738383 | 880.945344 | 2.22222222 | 0.50544093 | 848.300218 | 4.1509434 |
| 0.17180975 | 1080.94534 | 2.22222222 | 0.29491134 | 1635.79151 | 1.88679245 |
| 0.32874887 | 1195.88434 | 5.92592593 | 0.30905443 | 1357.90585 | 3.77358491 |
| 0.33885021 | 2183.51684 | 1.48148148 | 0.26488257 | 1753.52722 | 0.75471698 |
| 0.35461023 | 1047.54179 | 7.40740741 | 0.45757585 | 1274.3864 | 11.3207547 |
| 0.2974306 | 1036.5061 | 5.92592593 | 0.32329089 | 863.953702 | 7.54716981 |
| 0.32674902 | 1254.58598 | 1.48148148 | 0.29340076 | 1045.09925 | 1.13207547 |
| 0.31624149 | 831.137459 | 2.22222222 | 0.29613518 | 1307.11813 | 4.52830189 |
| 0.42474247 | 925.66679 | 8.88888889 | 0.45609899 | 999.092157 | 10.5660377 |

**Fig. S2D (d)** Data Source File

| | | | | | |
|---|---|---|---|---|---|
| 0.37989444 | 999.4967 | 2.22222222 | 0.35103402 | 1063.98515 | 2.64150943 |
| 0.38408306 | 945.101387 | 0.74074074 | 0.38883102 | 1280.34639 | 2.64150943 |
| 0.29350632 | 1094.01885 | 4.44444444 | 0.2773692 | 1130.45619 | 3.77358491 |
| 0.33176199 | 1447.87364 | 7.40740741 | 0.35364889 | 1176.79424 | 4.1509434 |
| 0.56784597 | 1341.72503 | 2.96296296 | 0.29816491 | 1164.64463 | 2.26415094 |
| 0.32140811 | 1503.23941 | 3.7037037 | 0.31824799 | 876.004502 | 3.39622642 |
| 0.3235274 | 1142.46868 | 8.88888889 | 0.3278835 | 1070.80454 | 9.05660377 |
| 0.28445782 | 1028.8927 | 0.74074074 | 0.28171075 | 1100.74251 | 0.75471698 |
| 0.30783208 | 928.892698 | 0.74074074 | 0.27311587 | 994.568828 | 0.75471698 |
| 0.51242896 | 772.852819 | 0.74074074 | 0.31433976 | 1119.83276 | 1.88679245 |
| 0.33165333 | 1300.73823 | 3.7037037 | 0.37576342 | 1383.34402 | 5.66037736 |
| 0.33530224 | 1034.99956 | 7.40740741 | 0.30814624 | 1129.75179 | 6.03773585 |
| 0.30490832 | 951.225203 | 0.74074074 | 0.40206927 | 1189.69602 | 1.50943396 |
| 0.37339458 | 2665.15974 | 1.48148148 | 0.44267514 | 2179.69602 | 1.50943396 |
| 0.42122095 | 957.96253 | 15.5555556 | 0.41801146 | 995.051124 | 13.5849057 |
| 0.50780418 | 1129.34769 | 1.48148148 | 0.43047179 | 1317.77781 | 5.28301887 |
| 0.37742668 | 1223.56301 | 9.62962963 | 0.4140086 | 1079.80176 | 10.1886792 |
| 0.31230175 | 930.54327 | 6.66666667 | 0.31676636 | 871.148742 | 6.41509434 |
| 0.27064884 | 1571.85159 | 2.96296296 | 0.29347778 | 2530.71838 | 1.88679245 |
| 0.3117067 | 1969.58243 | 2.96296296 | 0.33634557 | 1269.43829 | 3.77358491 |
| 0.28460516 | 1503.15909 | 3.7037037 | 0.34113817 | 1391.43021 | 5.66037736 |
| 0.42306753 | 3186.04541 | 2.96296296 | 0.28037074 | 1046.64682 | 1.50943396 |
| 0.32182142 | 1182.6004 | 5.18518519 | 0.39812034 | 1186.4626 | 4.52830189 |
| 0.65132005 | 817.819962 | 5.92592593 | 0.42435388 | 967.389085 | 4.90566038 |
| 0.44453113 | 1748.18096 | 2.96296296 | 0.40253826 | 1419.35888 | 2.26415094 |
| 0.67056853 | 7925.90412 | 0.74074074 | 0.44611865 | 2622.77958 | 1.88679245 |
| 0.36307535 | 1159.26323 | 11.1111111 | 0.35771435 | 1083.69228 | 10.9433962 |
| 0.32341255 | 1074.679 | 10.3703704 | 0.36844127 | 1137.69873 | 7.54716981 |
| 0.3799067 | 1442.74817 | 8.14814815 | 0.32410529 | 1212.30753 | 7.16981132 |
| 0.29136687 | 1676.93967 | 3.7037037 | 0.33002591 | 1393.39504 | 1.88679245 |
| 0.37067588 | 1159.4375 | 12.5925926 | 0.34147093 | 1006.6638 | 12.4528302 |
| 0.33688784 | 1461.49814 | 7.40740741 | 0.33190153 | 1028.81186 | 6.79245283 |
| 0.35604087 | 1063.64915 | 5.92592593 | 0.63868371 | 939.610427 | 6.79245283 |
| 0.26658548 | 1347.98567 | 0.74074074 | 0.44789044 | 1089.51199 | 1.88679245 |
| 0.352061 | 1331.69905 | 7.40740741 | 0.34744625 | 1049.34293 | 9.05660377 |
| 0.34022131 | 1028.04348 | 8.14814815 | 0.3598359 | 1173.1984 | 8.30188679 |
| 0.36511897 | 1268.01934 | 5.92592593 | 0.38841481 | 1596.66431 | 4.52830189 |
| 0.36888058 | 1400.68178 | 5.41627207 | 0.36019079 | 1345.92496 | 5.23035455 |
| 0.01082292 | 66.133883 | 0.39168844 | 0.00933625 | 55.2126949 | 0.38251684 |

**Wilcoxon matched-pairs signed rank test**

| Astro. transients | Amplitude (df/f) | Duration (ms) | Frequency (Events Nb/min) |
|---|---|---|---|
| P value | P=0.2947 | P=0.4754 | P=0.1076 |

**Fig. S2D (e)**

Data Source File

| Neuronal $Ca^{2+}$ transients | aCSF Amplitude (df/f) | aCSF Duration (ms) | aCSF Frequency (Events nb/min) | TTX Amplitude (df/f) | TTX Duration (ms) | TTX Frequency (Events nb/min) |
|---|---|---|---|---|---|---|
| | 0.335408445 | 1930.89736 | 5 | 0 | 0 | 0 |
| | 0.245817635 | 2868.7807 | 9 | 0 | 0 | 0 |
| | 0.227210015 | 3609.08324 | 0.66666667 | 0 | 0 | 0 |
| | 0.329818545 | 8626.89764 | 2.66666667 | 0 | 0 | 0 |
| | 0.19749239 | 2664.3586 | 5 | 0 | 0 | 0 |
| | 0.19214739 | 3639.82725 | 0.33333333 | 0 | 0 | 0 |
| | 0.18767138 | 1812.12635 | 0.33333333 | 0 | 0 | 0 |
| | 0.170833255 | 1813.05903 | 1 | 0 | 0 | 0 |
| | 0.247472315 | 2592.4742 | 3.66666667 | 0 | 0 | 0 |
| | 0.107565135 | 2226.84773 | 0.5 | 0 | 0 | 0 |
| | 0.119361705 | 2437.8292 | 0.5 | 0 | 0 | 0 |
| | 0.108665435 | 2851.92536 | 1 | 0 | 0 | 0 |
| | 0.110066195 | 1633.02033 | 0.5 | 0 | 0 | 0 |
| | 0.128269125 | 2253.10284 | 3.5 | 0 | 0 | 0 |
| | 0.17070808 | 1367.51401 | 21 | 0.009627649 | 0 | 0.66666667 |
| | 0.13764955 | 2078.26517 | 7.5 | 0 | 0 | 0 |
| | 0.186376745 | 1413.74221 | 17 | 0.043851077 | 82.21059 | 1.33333333 |
| | 0.142998015 | 1884.44132 | 10 | 0 | 0 | 0 |
| | 0.1176712 | 1600.74495 | 3 | 0 | 0 | 0 |
| | 0.11770529 | 1458.692 | 2 | 0.012731303 | 15.68741 | 0.66666667 |
| | 0.146704575 | 1736.48868 | 10 | 0 | 0 | 0 |
| | 0.13202443 | 2031.41413 | 7 | 0 | 0 | 0 |
| | 0.117517805 | 3358.3074 | 2 | 0 | 0 | 0 |
| | 0.157807155 | 1519.68238 | 9 | 0 | 0 | 0 |
| | 0.290888395 | 3410.54357 | 7 | 0 | 0 | 0 |
| | 0.12538251 | 2656.59576 | 4 | 0 | 0 | 0 |
| | 0.160552665 | 2856.23798 | 1 | 0.01217451 | 108.16457 | 0.66666667 |
| | 0.117160315 | 1833.72123 | 3 | 0 | 0 | 0 |
| | 0.17195915 | 1378.4864 | 16 | 0.014933946 | 73.77043 | 0.33333333 |
| | 0.135986015 | 2255.3912 | 3 | 0 | 0 | 0 |
| | 0.15086303 | 1600.23952 | 9 | 0 | 0 | 0 |
| | 0.141434045 | 1772.0306 | 9 | 0.033645093 | 157.25539 | 0.66666667 |
| | 0.1831579 | 1697.81131 | 13.5 | 0.055 | 137.18232 | 0.66666667 |
| | 0.121149465 | 3325.18864 | 0.5 | 0 | 0 | 0 |
| | 0.10147503 | 1253.24347 | 0.5 | 0 | 0 | 0 |
| | 0.116162305 | 3738.63147 | 1 | 0.05 | 0 | 0.33333333 |
| | 0.1790699 | 1598.5478 | 13.5 | 0 | 0 | 0 |
| | 0.168491385 | 1614.84375 | 10 | 0 | 0 | 0 |
| | 0.12414994 | 6002.56483 | 0.66666667 | 0 | 0 | 0 |
| | 0.1144247 | 2461.14149 | 1.66666667 | 0 | 0 | 0 |
| | 0.123693815 | 1957.0522 | 3.66666667 | 0 | 0 | 0 |
| | 0.121729485 | 2187.3186 | 1 | 0 | 0 | 0 |
| | 0.115976435 | 1128.81104 | 1.66666667 | 0 | 0 | 0 |
| | 0.10461741 | 2457.62812 | 0.33333333 | 0 | 0 | 0 |
| | 0.12271023 | 2645.56871 | 5 | 0 | 0 | 0 |
| | 0.13733603 | 1664.98606 | 9.33333333 | 0 | 0 | 0 |
| | 0.143209435 | 1916.72534 | 7.66666667 | 0 | 0 | 0 |
| | 0.13448399 | 2512.18235 | 4 | 0 | 0 | 0 |
| | 0.10728787 | 3551.85831 | 0.66666667 | 0 | 0 | 0 |
| | 0.109087625 | 2060.66021 | 1.33333333 | 0 | 0 | 0 |

**Fig. S2D (f)**

Data Source File

| | | | | | |
|---|---|---|---|---|---|
| 0.121260105 | 1880.98875 | 0.66666667 | 0 | 0 | 0 |
| 0.10633275 | 2022.17781 | 0.66666667 | 0 | 0 | 0 |
| 0.16079358 | 2112.22187 | 5 | 0 | 0 | 0 |
| 0.13715881 | 2293.53429 | 2.33333333 | 0 | 0 | 0 |
| 0.112169955 | 2268.024 | 2.66666667 | 0 | 0 | 0 |
| 0.118251455 | 1993.52232 | 0.66666667 | 0 | 0 | 0 |
| 0.121685815 | 2020.7673 | 5.33333333 | 0 | 0 | 0 |
| 0.113624835 | 3230.55659 | 0.33333333 | 0 | 0 | 0 |
| 0.105456855 | 3433.59038 | 1 | 0 | 0 | 0 |
| 0.102716925 | 1226.89753 | 0.33333333 | 0 | 0 | 0 |
| 0.139645655 | 3226.55254 | 2.66666667 | 0 | 0 | 0 |
| 0.235488535 | 1465.65095 | 17 | 0 | 0 | 0 |
| 0.125832735 | 2030.86354 | 4.66666667 | 0 | 0 | 0 |
| 0.133136605 | 1769.2715 | 4.66666667 | 0 | 0 | 0 |
| 0.11267223 | 2933.32857 | 2.33333333 | 0 | 0 | 0 |
| 0.14319922 | 1296.71982 | 1.66666667 | 0 | 0 | 0 |
| 0.10039375 | 2249.33069 | 0.33333333 | 0 | 0 | 0 |
| 0.23736454 | 2932.38818 | 3.66666667 | 0 | 0 | 0 |
| 0.108162885 | 2589.91415 | 1.33333333 | 0 | 0 | 0 |
| 0.11178563 | 1584.47102 | 2.33333333 | 0 | 0 | 0 |
| 0.225203505 | 9313.8087 | 0.33333333 | 0.01 | 90.2119 | 0.5 |
| 0.123366895 | 1547.79658 | 3 | 0 | 0 | 0 |
| 0.291630925 | 1861.4108 | 3 | 0 | 0 | 0 |
| 0.125264165 | 2183.7637 | 0.66666667 | 0 | 0 | 0 |
| 0.14860688 | 1374.15175 | 2.66666667 | 0 | 0 | 0 |
| 0.1144247 | 2461.14149 | 1.66666667 | 0 | 0 | 0 |
| 0.121729485 | 2187.3186 | 1 | 0 | 0 | 0 |
| 0.115976435 | 1128.81104 | 1.66666667 | 0 | 0 | 0 |
| 0.12271023 | 2645.56871 | 5 | 0.05 | 170.46323 | 1.5 |
| 0.13733603 | 1664.98606 | 9.33333333 | 0 | 0 | 0 |
| 0.143209435 | 1916.72534 | 7.66666667 | 0 | 0 | 0 |
| 0.13448399 | 2512.18235 | 4 | 0 | 0 | 0 |
| 0.109087625 | 2060.66021 | 1.33333333 | 0.037664804 | 54.0685 | 0.5 |
| 0.118251455 | 1993.52232 | 0.66666667 | 0 | 0 | 0 |
| 0.113624835 | 3230.55659 | 0.33333333 | 0 | 0 | 0 |
| 0.105456855 | 3433.59038 | 1 | 0.032236993 | 149.44535 | 0.25 |
| 0.235488535 | 1465.65095 | 17 | 0.015 | 76.83934 | 0.75 |
| 0.11961637 | 2184.94806 | 2.5 | 0 | 0 | 0 |
| 0.292271035 | 3035.14514 | 3 | 0 | 0 | 0 |
| 0.13223857 | 2260.73366 | 2 | 0 | 0 | 0 |
| 0.146451705 | 1501.5218 | 6.5 | 0.033033746 | 18.89296 | 1 |
| 0.28146111 | 1644.26124 | 9.5 | 0.040242237 | 165.23959 | 0.5 |
| 0.11842502 | 2617.50839 | 1 | 0 | 0 | 0 |
| 0.1721653 | 1763.47162 | 6.5 | 0 | 0 | 0 |
| 0.142223505 | 2183.06557 | 4.5 | 0.027977238 | 117.06439 | 0.5 |
| 0.127848165 | 2182.62534 | 1.5 | 0 | 0 | 0 |
| 0.20017787 | 1604.56543 | 6 | 0 | 0 | 0 |
| 0.164379585 | 6803.87784 | 1.5 | 0 | 0 | 0 |
| 0.17682014 | 1295.24839 | 7.5 | 0 | 0 | 0 |
| 0.12207267 | 2065.30078 | 2 | 0 | 0 | 0 |

**Fig. S2D (g)**

Data Source File

| | | | | | |
|---|---|---|---|---|---|
| 0.22045019 | 1353.09101 | 3.5 | 0 | 0 | 0 |
| 0.12585381 | 1302.66175 | 0.5 | 0 | 0 | 0 |
| 0.225124895 | 1480.93947 | 7.5 | 0 | 0 | 0 |
| 0.14500619 | 2125.13188 | 1 | 0 | 0 | 0 |
| 0.242788435 | 1493.33193 | 10 | 0 | 0 | 0 |
| 0.124730325 | 1409.44891 | 1 | 0 | 0 | 0 |
| 0.12139266 | 1644.3694 | 3.5 | 0.036298646 | 133.14855 | 1.5 |
| 0.12218666 | 1459.28721 | 2 | 0 | 0 | 0 |
| 0.13178627 | 2001.13284 | 1.5 | 0 | 0 | 0 |
| 0.23678 | 1784.00128 | 4.5 | 0 | 0 | 0 |
| 0.109827795 | 1902.13109 | 2 | 0 | 0 | 0 |
| 0.35478723 | 1192.50206 | 12.5 | 0 | 0 | 0 |
| 0.109449915 | 1060.88236 | 0.5 | 0 | 0 | 0 |
| 0.12434136 | 2310.82543 | 0.5 | 0 | 0 | 0 |
| 0.23946392 | 1238.33317 | 11 | 0 | 0 | 0 |
| 0.128823615 | 2369.87596 | 4 | 0 | 0 | 0 |
| 0.262349935 | 1675.75509 | 5 | 0 | 0 | 0 |
| 0.15905835 | 2529.34602 | 1 | 0 | 0 | 0 |
| 0.208799315 | 3454.66864 | 0.5 | 0 | 0 | 0 |
| 0.45658253 | 4508.44383 | 2 | 0 | 0 | 0 |
| 0.229485745 | 6464.45711 | 3 | 0 | 0 | 0 |
| 0.32411537 | 2080.93252 | 0.5 | 0 | 0 | 0 |
| 0.29460432 | 2836.5806 | 2.5 | 0 | 0 | 0 |
| 0.294891825 | 1331.66243 | 7.5 | 0 | 0 | 0 |
| 0.26626353 | 2830.97869 | 7.5 | 0 | 0 | 0 |
| 0.270901535 | 2553.3946 | 7.5 | 0 | 0 | 0 |
| 0.242385615 | 1308.12344 | 0.5 | 0 | 0 | 0 |
| 0.27301582 | 1507.88162 | 5.5 | 0 | 0 | 0 |
| 0.24563925 | 4527.69618 | 1 | 0 | 0 | 0 |
| 0.291580315 | 2442.20099 | 3 | 0 | 0 | 0 |
| 0.287268825 | 2351.58137 | 9.5 | 0 | 0 | 0 |
| 0.28341919 | 1912.06344 | 7.5 | 0 | 0 | 0 |
| 0.235078575 | 3624.66987 | 1 | 0 | 0 | 0 |
| 0.30444449 | 1084.8863 | 2.5 | 0 | 0 | 0 |
| 0.223750505 | 1577.7546 | 1 | 0 | 0 | 0 |
| 0.27819888 | 1911.13101 | 4.5 | 0 | 0 | 0 |
| 0.30055587 | 1652.43978 | 11 | 0 | 0 | 0 |
| 0.21612565 | 1102.68223 | 0.5 | 0 | 0 | 0 |
| 0.27683663 | 1604.4382 | 12.5 | 0 | 0 | 0 |
| 0.21457306 | 2281.47376 | 0.5 | 0 | 0 | 0 |
| 0.20872986 | 903.442287 | 0.5 | 0 | 0 | 0 |
| 0.1737 | 2311 | 4.168 | 0.003648 | 10.99 | 0.08747 |
| 0.005908 | 104.4 | 0.3565 | 0.0009395 | 3.004 | 0.02281 |

**Wilcoxon matched-pairs signed rank test**

| Neuronal transients | Amplitude (df/f) | Duration (ms) | Frequency (Events Nb/min) |
|---|---|---|---|
| P value | P<0.0001 | P<0.0001 | P<0.0001 |

**Fig. S4C**

Data Source File

| | Oscill. aCSF | +Barium | Oscill. aCSF | +Barium | Oscill. aCSF | +Barium |
|---|---|---|---|---|---|---|
| | Ampl oscill. (mV) | Ampl oscill. (mV) | Duration oscill. (s) | Duration oscill. (s) | Frequ. Oscill. (Hz) | Frequ. oscill. (Hz) |
| mouse#1 | 4.54666667 | 5.83 | 2.56 | 5.8875 | 0.05 | 0.03333333 |
| mouse#2 | 6.97016042 | 9.34989738 | 8.99613646 | 29.97 | 0.9153198 | 0.69519226 |
| mouse#3 | 7.12017229 | 11.8529858 | 2.2 | 4.87 | 0.36794739 | 0.2061254 |
| mouse#4 | 5.931331 | 7.20546128 | 0.96836496 | 3.54458898 | 0.53085351 | 0.32105185 |
| mouse#4 | 10.6168564 | 13 | 7.07933746 | 8.3 | 0.1 | 0.04444444 |
| mouse#5 | 8.268 | 10.27325 | 5.37 | 6.98413158 | 0.27328378 | 0.22 |
| mouse#5 | 6.42694052 | 7.94292428 | 1.345 | 2.629 | 0.46851179 | 0.33152585 |
| mouse#6 | 0.31 | 0.68 | 2.08 | 4.46 | 1.5 | 0.8 |
| mouse#7 | 4.15 | 5.28 | 0.576 | 2.04 | 1.6 | 0.5 |
| MEAN | 6.038 | 7.935 | 3.464 | 7.632 | 0.6451 | 0.3502 |
| SEM | 0.9628 | 1.253 | 0.9909 | 2.871 | 0.191 | 0.08949 |

| Ampl N oscill. (mV) | | Duration N oscill (s) | | Frequ. N oscill. (Hz) | |
|---|---|---|---|---|---|
| Wilcoxon matched-pairs signed rank tests | | | | | |
| P value | 0.0039 | P value | 0.0039 | P value | 0.0039 |
| P value summary | ** | P value summary | ** | P value summ ary | ** |

| | TTX+Cocktail | +Barium | TTX+Cocktail | +Barium | TTX+Cocktail | +Barium |
|---|---|---|---|---|---|---|
| | Ampl oscill. (mV) | Ampl oscill. (mV) | Duration oscill. (s) | Duration oscill. (s) | Frequ. Oscill. (Hz) | Frequ. oscill. (Hz) |
| mouse#1 | 12.3833333 | 17.40875 | 0.62066667 | 7.04625 | 1.5 | 0.4 |
| mouse#1 | 29.04375 | 38.805 | 6.74375 | 11.4975 | 0.16 | 0.08 |
| mouse#1 | 7.23545455 | 11.8457143 | 0.64363636 | 4.13428571 | 1.1 | 0.11666667 |
| mouse#2 | 39.383 | 40.815 | 1.274 | 12.4583333 | 0.5 | 0.06 |
| mouse#3 | 4.55 | 8.88 | 0.1 | 0.234 | 3 | 1.28 |
| mouse#3 | 22 | 25 | 1.15 | 37 | 0.83333333 | 0.05 |
| MEAN | 19.1 | 23.79 | 1.755 | 12.06 | 1.182 | 0.3311 |
| SEM | 5.527 | 5.545 | 1.012 | 5.324 | 0.41 | 0.1972 |

| Ampl N oscill. (mV) | | Duration N oscill (s) | | Frequ. N oscill. | |
|---|---|---|---|---|---|
| Wilcoxon matched-pairs signed rank tests | | | | | |
| P value | 0.0313 | P value | 0.0313 | P value | 0.0313 |
| P value summary | * | P value summary | * | P value summ ary | * |

**Fig. S6C (a)**

Data Source File

| | | Ctrl-ShRNA | |
|---|---|---|---|
| | | age | Vrest (mV) |
| mouse#1 | astro1 | P13 | -87.2568771 |
| mouse#1 | astro2 | P13 | -90.2398026 |
| mouse#1 | astro3 | P13 | -85.113826 |
| mouse#1 | astro4 | P13 | -87.1813965 |
| mouse#1 | astro5 | P13 | -91.6263016 |
| mouse#1 | astro6 | P13 | -89.2073839 |
| mouse#1 | astro7 | P13 | -89.7961327 |
| mouse#2 | astro8 | P13 | -90.4 |
| mouse#2 | astro9 | P13 | -83.4 |
| mouse#2 | astro10 | P13 | -97.7 |
| mouse#2 | astro11 | P13 | -89.4 |
| mouse#3 | astro12 | P13 | -88.4 |
| mouse#3 | astro13 | P13 | -83.54 |
| | | MEAN | -88.71 |
| | | SEM | 1.04 |

| | | Kir4.1-ShRNA | |
|---|---|---|---|
| | | age | Vrest (mV) |
| mouse#1 | astro1 | P8 | -73.1946663 |
| mouse#1 | astro2 | P8 | -90.022448 |
| mouse#1 | astro3 | P8 | -82.6924973 |
| mouse#1 | astro4 | P8 | -82.5735153 |
| mouse#1 | astro5 | P8 | -81.9515114 |
| mouse#2 | astro6 | P11 | -78.29 |
| mouse#2 | astro7 | P11 | -75.59 |
| mouse#2 | astro8 | P11 | -82.88 |
| mouse#2 | astro9 | P11 | -80.88 |
| mouse#3 | astro10 | P12 | -85.4 |
| mouse#3 | astro11 | P12 | -81.9 |
| mouse#3 | astro12 | P12 | -81.89 |
| mouse#3 | astro13 | P12 | -65.05 |
| mouse#4 | astro14 | P13 | -70.2 |
| | | MEAN | -79.47 |
| | | SEM | 1.735 |

| | |
|---|---|
| Vrest (mV) | |
| Mann Whitney test | |
| P value | <0.0001 |
| P value summary | **** |

**Fig. S6C (b)** Data Source File

| | | | Ctrl-ShRNA |
|---|---|---|---|
| | | age | Rin (Mohm) |
| mouse#1 | astro1 | P13 | 35.0382784 |
| mouse#1 | astro2 | P13 | 20.850366 |
| mouse#1 | astro3 | P13 | 17.1491475 |
| mouse#1 | astro4 | P13 | 1.54380798 |
| mouse#1 | astro5 | P13 | 48.2971308 |
| mouse#1 | astro6 | P13 | 10.8887439 |
| mouse#1 | astro7 | P13 | 10.8101619 |
| mouse#2 | astro8 | P13 | 15 |
| mouse#2 | astro9 | P13 | 20.5 |
| mouse#2 | astro10 | P13 | 23 |
| mouse#2 | astro11 | P13 | 22.5 |
| mouse#3 | astro12 | P13 | 24 |
| mouse#3 | astro13 | P13 | 20 |
| | | MEAN | 20.7367413 |
| | | SEM | 3.196 |

| | | | Kir4.1-ShRNA |
|---|---|---|---|
| | | | Rin (Mohm) |
| mouse#1 | astro1 | P8 | 39.7168901 |
| mouse#1 | astro2 | P8 | 24.1497026 |
| mouse#1 | astro3 | P8 | 32.8467602 |
| mouse#1 | astro4 | P8 | 25.835406 |
| mouse#1 | astro5 | P8 | 27.9202935 |
| mouse#2 | astro6 | P11 | 73.33 |
| mouse#2 | astro7 | P11 | 12 |
| mouse#2 | astro8 | P11 | 25 |
| mouse#2 | astro9 | P11 | 44 |
| mouse#3 | astro10 | P12 | 17 |
| mouse#3 | astro11 | P12 | 128 |
| mouse#3 | astro12 | P12 | 56 |
| mouse#3 | astro13 | P12 | 91 |
| mouse#4 | astro14 | P13 | 48 |
| | | MEAN | 46.06 |
| | | SEM | 8.601 |

| Rin (Mohm) | |
|---|---|
| Mann Whitney test | |
| P value | 0.0033 |
| P value summary | ** |

**Fig. S6E** Data Source File

Ctrl-ShRNA n=12 astrocytes from 3 mice

| | Current (pA) | | |
|---|---|---|---|
| Vm (mV) | MEAN | SEM | N |
| -150 | -2299 | 443.9 | 12 |
| -140 | -1856 | 315.8 | 12 |
| -130 | -1383 | 272.5 | 12 |
| -120 | -1049 | 210.2 | 12 |
| -110 | -753.4 | 164.2 | 12 |
| -100 | -385.3 | 84.73 | 12 |
| -90 | -38.51 | 35.99 | 12 |
| -80 | 317.1 | 85.25 | 12 |
| -70 | 668.7 | 168.2 | 12 |
| -60 | 1018 | 253.9 | 12 |
| -50 | 1370 | 348.1 | 12 |
| -40 | 1698 | 428.6 | 12 |
| -30 | 1944 | 388.3 | 12 |
| -20 | 2108 | 510.6 | 12 |
| -10 | 2444 | 617.7 | 12 |

Kir4.1-ShRNA n=14 astrocytes from 4 mice

| | Current (pA) | | |
|---|---|---|---|
| Vm (mV) | MEAN | SEM | N |
| -150 | -1372 | 178.3 | 14 |
| -140 | -1060 | 228.9 | 14 |
| -130 | -935.8 | 165 | 14 |
| -120 | -769.7 | 126.6 | 14 |
| -110 | -572.3 | 101.4 | 14 |
| -100 | -343 | 87.09 | 14 |
| -90 | -64.96 | 61.29 | 14 |
| -80 | 209.3 | 77.76 | 14 |
| -70 | 478.2 | 117.6 | 14 |
| -60 | 743.8 | 161.1 | 14 |
| -50 | 1010 | 206.7 | 14 |
| -40 | 1247 | 247.5 | 14 |
| -30 | 1484 | 289.9 | 14 |
| -20 | 1683 | 340.4 | 14 |
| -10 | 1926 | 451.1 | 14 |

Linear Regression Comp.
Are the slopes equal?
F = 168.4. DFn = 1, DFd = 26
P<0.0001

If the overall slopes were identical, there is less than a 0.01% chance of randomly choosing data points with slopes this different. You can conclude that the differences between the slopes are extremely significant.

Are the slopes equal?
F = 168.4. DFn = 1, DFd = 26
P<0.0001

If the overall slopes were identical, there is less than a 0.01% chance of randomly choosing data points with slopes this different. You can conclude that the differences between the slopes are extremely significant.

**Fig. S6F** Data Source File

| Ctrl-ShRNA | | Peak Ikir at -140mV |
|---|---|---|
| mouse#1 | astro1 | -1472.64136 |
| mouse#1 | astro2 | -888.434387 |
| mouse#1 | astro3 | -2219.03491 |
| mouse#1 | astro4 | -807.113647 |
| mouse#1 | astro5 | -1872.15381 |
| mouse#1 | astro6 | -842.399109 |
| mouse#1 | astro7 | -3081.34583 |
| mouse#2 | astro8 | -4345.09277 |
| mouse#2 | astro9 | -1186.9812 |
| mouse#2 | astro10 | -902.357483 |
| mouse#2 | astro11 | -2729.89917 |
| mouse#3 | astro12 | -1924.1842 |
| | MEAN | -1856 |
| | SEM | 315.8 |

| Kir4.1-ShRNA | | Peak Ikir at -140mV |
|---|---|---|
| mouse#1 | astro1 | -1842.6062 |
| mouse#1 | astro2 | -1191.79285 |
| mouse#1 | astro3 | -758.172913 |
| mouse#1 | astro4 | -285.20462 |
| mouse#1 | astro5 | -2542.7085 |
| mouse#2 | astro6 | -362.642731 |
| mouse#2 | astro7 | -621.13443 |
| mouse#2 | astro8 | -650.6642 |
| mouse#2 | astro9 | -2453.24536 |
| mouse#3 | astro10 | -1020.71558 |
| mouse#3 | astro11 | -549.01123 |
| mouse#3 | astro12 | -469.526587 |
| mouse#3 | astro13 | -985.20462 |
| mouse#4 | astro14 | -1112.44202 |
| | MEAN | -1060 |
| | SEM | 195.1 |

| Mann Whitney test | |
|---|---|
| P value | 0.027 |
| P value summ | * |

**Fig. S6H** Data Source File

Ctrl-ShRNA n=6 astrocytes from 2 mice

| | Ba2+-sensitive Current (pA) | | |
|---|---|---|---|
| Vm (mV) | MEAN | SEM | N |
| -140 | -1233.70969 | 219.822908 | 6 |
| -130 | -867.644023 | 177.991699 | 6 |
| -120 | -674.987762 | 146.986005 | 6 |
| -110 | -492.252345 | 104.669055 | 6 |
| -100 | -495.584152 | 189.126371 | 6 |
| -90 | -217.844957 | 77.6172805 | 6 |
| -80 | 24.7187213 | 20.4692883 | 6 |
| -70 | 240.8688 | 103.985 | 6 |
| -60 | 427.026347 | 195.362066 | 6 |
| -50 | 595.066442 | 277.968966 | 6 |
| -40 | 744.191884 | 353.399066 | 6 |
| -30 | 893.091858 | 406.752602 | 6 |
| -20 | 1094.17622 | 494.70103 | 6 |
| -10 | 1219.81226 | 626.467086 | 6 |

Kir4.1-ShRNA n=6 astrocytes from 2 mice

| | Ba2+-sensitive Current (pA) | | |
|---|---|---|---|
| Vm (mV) | MEAN | SEM | N |
| -140 | -249.588355 | 117.125807 | 7 |
| -130 | -192.928768 | 76.2377877 | 7 |
| -120 | -145.2158 | 49.106133 | 7 |
| -110 | -98.0447355 | 26.9356396 | 7 |
| -100 | -115.409827 | 59.192593 | 7 |
| -90 | -70.6714345 | 41.5672984 | 7 |
| -80 | -12.8586912 | 11.8935372 | 7 |
| -70 | 24.5110189 | 9.35520348 | 7 |
| -60 | 69.6540101 | 22.842338 | 7 |
| -50 | 109.944425 | 42.2531668 | 7 |
| -40 | 142.969312 | 48.7858707 | 7 |
| -30 | 163.741768 | 57.5884227 | 7 |
| -20 | 202.27392 | 70.2705508 | 7 |
| -10 | 205.048825 | 64.8181466 | 7 |

Are the slopes equal?
F = 86.40. DFn = 1, DFd = 175
P<0.0001

If the overall slopes were identical, there is less than a 0.01% chance of randomly choosing data points with slopes this different. You can conclude that the differences between the slopes are extremely significant.

**Fig. S6I**

Data Source File

| | | Ba2+-sensitive current at -140mV |
|---|---|---|
| | | Ctrl-ShRNA |
| mouse#1 | P13 | -1798.48511 |
| mouse#1 | P13 | -1689.24976 |
| mouse#1 | P13 | -472.106934 |
| mouse#2 | P13 | -1335.56323 |
| mouse#2 | P13 | -687.531494 |
| mouse#2 | P13 | -1419.32159 |
| | MEAN | -1234 |
| | SEM | 219.8 |

| | | Ba2+-sensitive current at -140mV |
|---|---|---|
| | | Kir4.1-ShRNA |
| mouse#1 | P11 | -42.868103 |
| mouse#1 | P11 | -109.002686 |
| mouse#1 | P11 | -914.764404 |
| mouse#1 | P11 | -331.838745 |
| mouse#2 | P12 | -76.9786377 |
| mouse#2 | P12 | -202.084366 |
| mouse#2 | P12 | -69.581543 |
| | MEAN | -249.6 |
| | SEM | 117.1 |

**Mann Whitney test**

| | |
|---|---|
| P value | 0.0047 |
| P value summary | ** |

**Fig. S7B (a)** Data Source File

| | | MNs | Label | cell body cross-sectional area (um2) |
|---|---|---|---|---|
| Ctrl-ShRNA | mouse#1 | 1 | MMP9(+) | 892.95 |
| | mouse#1 | 2 | MMP9(+) | 976.11 |
| | mouse#1 | 3 | MMP9(+) | 1001.03 |
| | mouse#1 | 4 | MMP9(+) | 657.53 |
| | mouse#1 | 5 | MMP9(+) | 713.14 |
| | mouse#1 | 1 | MMP9(+) | 788.82 |
| | mouse#1 | 2 | MMP9(+) | 1509.3 |
| | mouse#1 | 3 | MMP9(+) | 1276.99 |
| | mouse#1 | 4 | MMP9(+) | 997.99 |
| | mouse#1 | 5 | MMP9(+) | 679.7 |
| | mouse#1 | 6 | MMP9(+) | 919.41 |
| | mouse#1 | 7 | MMP9(+) | 891.08 |
| | mouse#1 | 8 | MMP9(+) | 922.27 |
| | mouse#1 | 9 | MMP9(+) | 657.52 |
| | mouse#1 | 1 | MMP9(+) | 303.08 |
| | mouse#1 | 2 | MMP9(+) | 742.89 |
| | mouse#1 | 3 | MMP9(+) | 808.53 |
| | mouse#1 | 4 | MMP9(+) | 647.69 |
| | mouse#1 | 5 | MMP9(+) | 939.4 |
| | mouse#1 | 6 | MMP9(+) | 741.03 |
| | mouse#1 | 7 | MMP9(+) | 880.22 |
| | mouse#1 | 8 | MMP9(+) | 577.72 |
| | mouse#1 | 9 | MMP9(+) | 619.45 |
| | mouse#1 | 1 | MMP9(+) | 1139.37 |
| | mouse#1 | 2 | MMP9(+) | 1029.05 |
| | mouse#1 | 3 | MMP9(+) | 464.51 |
| | mouse#1 | 4 | MMP9(+) | 642.82 |
| | mouse#1 | 5 | MMP9(+) | 705.07 |
| | mouse#1 | 6 | MMP9(+) | 776.76 |
| Ctrl-ShRNA | mouse#2 | 1 | MMP9(+) | 570.09 |
| | mouse#2 | 2 | MMP9(+) | 458.05 |
| | mouse#2 | 3 | MMP9(+) | 1045.03 |
| | mouse#2 | 4 | MMP9(+) | 604.8 |
| | mouse#2 | 5 | MMP9(+) | 729.78 |
| | mouse#2 | 6 | MMP9(+) | 1296.74 |
| | mouse#2 | 7 | MMP9(+) | 739.09 |
| | mouse#2 | 1 | MMP9(+) | 1610.07 |
| | mouse#2 | 2 | MMP9(+) | 865.19 |
| | mouse#2 | 3 | MMP9(+) | 749.47 |
| | mouse#2 | 4 | MMP9(+) | 1072.14 |
| | mouse#2 | 5 | MMP9(+) | 952.86 |
| | mouse#2 | 6 | MMP9(+) | 1124.09 |
| | mouse#2 | 7 | MMP9(+) | 941.28 |
| | mouse#2 | 8 | MMP9(+) | 1034.44 |
| | mouse#2 | 9 | MMP9(+) | 963.02 |
| | mouse#2 | 10 | MMP9(+) | 499.34 |
| | mouse#2 | 1 | MMP9(+) | 665.16 |
| | mouse#2 | 2 | MMP9(+) | 975.35 |
| | mouse#2 | 3 | MMP9(+) | 998.2 |

**Fig. S7B (b)** Data Source File

| | | MNs | Label | cell body cross-sectional area (um2) |
|---|---|---|---|---|
| | mouse#2 | 4 | MMP9(+) | 864.61 |
| | mouse#2 | 5 | MMP9(+) | 539.05 |
| | mouse#2 | 6 | MMP9(+) | 444.13 |
| | mouse#2 | 7 | MMP9(+) | 442.46 |
| | mouse#2 | 1 | MMP9(+) | 864.09 |
| | mouse#2 | 2 | MMP9(+) | 461.55 |
| | mouse#2 | 3 | MMP9(+) | 525.44 |
| | mouse#2 | 4 | MMP9(+) | 514.3 |
| | mouse#2 | 5 | MMP9(+) | 758.86 |
| | mouse#2 | 6 | MMP9(+) | 811.17 |
| | mouse#2 | 7 | MMP9(+) | 567.55 |
| | mouse#2 | 8 | MMP9(+) | 921.29 |
| | mouse#2 | 9 | MMP9(+) | 1232.88 |
| | mouse#2 | 10 | MMP9(+) | 996.83 |
| | mouse#2 | 11 | MMP9(+) | 616.91 |
| | mouse#2 | 12 | MMP9(+) | 777.04 |
| Ctrl-ShRNA | mouse#3 | 1 | MMP9(+) | 783.07 |
| | mouse#3 | 2 | MMP9(+) | 850.47 |
| | mouse#3 | 1 | MMP9(+) | 758.31 |
| | mouse#3 | 2 | MMP9(+) | 513.25 |
| | mouse#3 | 3 | MMP9(+) | 1058.99 |
| | mouse#3 | 4 | MMP9(+) | 736.14 |
| | mouse#3 | 5 | MMP9(+) | 1169.71 |
| | mouse#3 | 6 | MMP9(+) | 1094.51 |
| | mouse#3 | 7 | MMP9(+) | 807.03 |
| | mouse#3 | 1 | MMP9(+) | 1133.52 |
| | mouse#3 | 2 | MMP9(+) | 1195.83 |
| | mouse#3 | 3 | MMP9(+) | 726.99 |
| | mouse#3 | 4 | MMP9(+) | 1134.37 |
| | mouse#3 | 5 | MMP9(+) | 983.85 |
| | mouse#3 | 6 | MMP9(+) | 692.1 |
| | mouse#3 | 7 | MMP9(+) | 830.96 |
| | mouse#3 | 8 | MMP9(+) | 561.37 |
| | mouse#3 | 1 | MMP9(+) | 931.02 |
| | mouse#3 | 2 | MMP9(+) | 859.2 |
| | mouse#3 | 3 | MMP9(+) | 705.18 |
| | mouse#3 | 4 | MMP9(+) | 797.92 |
| | mouse#3 | 5 | MMP9(+) | 1091.8 |
| | mouse#3 | 6 | MMP9(+) | 740.18 |
| | | | MEAN | 832.835795 |
| | | | SEM | 25.92 |

| | | MNs | Label | cell body cross-sectional area (um2) |
|---|---|---|---|---|
| Kir4.1-ShRNA | mouse#1 | 1 | MMP9(+) | 853.51 |
| | mouse#1 | 2 | MMP9(+) | 818.53 |
| | mouse#1 | 3 | MMP9(+) | 693.55 |
| | mouse#1 | 4 | MMP9(+) | 594.28 |
| | mouse#1 | 5 | MMP9(+) | 564.2862 |

**Fig. S7B (c)** Data Source File

| | | MNs | Label | cell body cross-sectional area (um2) |
|---|---|---|---|---|
| | mouse#1 | 6 | MMP9(+) | 1059.14 |
| | mouse#1 | 7 | MMP9(+) | 956.23 |
| | mouse#1 | 8 | MMP9(+) | 887.33 |
| | mouse#1 | 9 | MMP9(+) | 531.4 |
| | mouse#1 | 10 | MMP9(+) | 474.73 |
| | mouse#1 | 1 | MMP9(+) | 1016.2 |
| | mouse#1 | 2 | MMP9(+) | 1370.75 |
| | mouse#1 | 3 | MMP9(+) | 1099.52 |
| | mouse#1 | 4 | MMP9(+) | 656.73 |
| | mouse#1 | 5 | MMP9(+) | 584.54 |
| | mouse#1 | 1 | MMP9(+) | 963.04 |
| | mouse#1 | 2 | MMP9(+) | 1028.92 |
| | mouse#1 | 3 | MMP9(+) | 832.05 |
| | mouse#1 | 4 | MMP9(+) | 1032.93 |
| | mouse#1 | 5 | MMP9(+) | 1072.53 |
| | mouse#1 | 6 | MMP9(+) | 625.78 |
| | mouse#1 | 7 | MMP9(+) | 834.98 |
| | mouse#1 | 8 | MMP9(+) | 520.73 |
| | mouse#1 | 1 | MMP9(+) | 697.8 |
| | mouse#1 | 2 | MMP9(+) | 1228.16 |
| | mouse#1 | 3 | MMP9(+) | 1066.75 |
| | mouse#1 | 4 | MMP9(+) | 978.85 |
| | mouse#1 | 5 | MMP9(+) | 1281.65 |
| | mouse#1 | 6 | MMP9(+) | 941.86 |
| Kir4.1-ShRNA | mouse#2 | 1 | MMP9(+) | 1248.37 |
| | mouse#2 | 2 | MMP9(+) | 651.17 |
| | mouse#2 | 3 | MMP9(+) | 919.15 |
| | mouse#2 | 4 | MMP9(+) | 819.97 |
| | mouse#2 | 5 | MMP9(+) | 779.51 |
| | mouse#2 | 6 | MMP9(+) | 744.94 |
| | mouse#2 | 7 | MMP9(+) | 519.66 |
| | mouse#2 | 8 | MMP9(+) | 1224.01 |
| | mouse#2 | 9 | MMP9(+) | 1301.75 |
| | mouse#2 | 10 | MMP9(+) | 639.15 |
| | mouse#2 | 11 | MMP9(+) | 1013.75 |
| | mouse#2 | 12 | MMP9(+) | 1291.75 |
| | mouse#2 | 1 | MMP9(+) | 1338.48 |
| | mouse#2 | 2 | MMP9(+) | 796.75 |
| | mouse#2 | 3 | MMP9(+) | 567.4 |
| | mouse#2 | 4 | MMP9(+) | 1037.4 |
| | mouse#2 | 5 | MMP9(+) | 580.38 |
| | mouse#2 | 6 | MMP9(+) | 616.41 |
| | mouse#2 | 7 | MMP9(+) | 1080.56 |
| | mouse#2 | 8 | MMP9(+) | 813.07 |
| | mouse#2 | 9 | MMP9(+) | 1220.59 |
| | mouse#2 | 10 | MMP9(+) | 1063.31 |
| | mouse#2 | 11 | MMP9(+) | 634.73 |
| | mouse#2 | 1 | MMP9(+) | 943.8 |
| | mouse#2 | 2 | MMP9(+) | 958.22 |
| | mouse#2 | 3 | MMP9(+) | 631.33 |

**Fig. S7B (d)** Data Source File

| | | | | |
|---|---|---|---|---|
| | mouse#2 | 4 | MMP9(+) | 830.39 |
| | mouse#2 | 5 | MMP9(+) | 1046.12 |
| | mouse#2 | 6 | MMP9(+) | 866.73 |
| | mouse#2 | 7 | MMP9(+) | 919.16 |
| | mouse#2 | 8 | MMP9(+) | 791.56 |
| | mouse#2 | 9 | MMP9(+) | 555.16 |
| | mouse#2 | 10 | MMP9(+) | 771.31 |
| | mouse#2 | 11 | MMP9(+) | 608.1 |
| | mouse#2 | 12 | MMP9(+) | 567.58 |
| | mouse#2 | 13 | MMP9(+) | 924 |
| | mouse#2 | 1 | MMP9(+) | 949.4 |
| | mouse#2 | 2 | MMP9(+) | 1154.96 |
| | mouse#2 | 3 | MMP9(+) | 941.23 |
| | mouse#2 | 4 | MMP9(+) | 966.1 |
| | mouse#2 | 5 | MMP9(+) | 801.37 |
| | mouse#2 | 6 | MMP9(+) | 1044.88 |
| | mouse#2 | 7 | MMP9(+) | 980.32 |
| | mouse#2 | 8 | MMP9(+) | 903.73 |
| | mouse#2 | 9 | MMP9(+) | 771.55 |
| | mouse#2 | 10 | MMP9(+) | 1008.96 |
| | mouse#2 | 11 | MMP9(+) | 695.18 |
| Kir4.1-ShRNA | mouse#3 | 1 | MMP9(+) | 1050.62 |
| | mouse#3 | 2 | MMP9(+) | 743.33 |
| | mouse#3 | 3 | MMP9(+) | 579.92 |
| | mouse#3 | 4 | MMP9(+) | 684.82 |
| | mouse#3 | 5 | MMP9(+) | 939.59 |
| | mouse#3 | 6 | MMP9(+) | 1022.96 |
| | mouse#3 | 7 | MMP9(+) | 934.33 |
| | mouse#3 | 1 | MMP9(+) | 1285.86 |
| | mouse#3 | 2 | MMP9(+) | 900.76 |
| | mouse#3 | 3 | MMP9(+) | 1087.76 |
| | mouse#3 | 4 | MMP9(+) | 736.68 |
| | mouse#3 | 5 | MMP9(+) | 1013.1 |
| | mouse#3 | 6 | MNP9(+) | 682.53 |
| | mouse#3 | 7 | MNP9(+) | 395.9 |
| | mouse#3 | 1 | MNP9(+) | 1064.91 |
| | mouse#3 | 2 | MNP9(+) | 886.68 |
| | mouse#3 | 3 | MNP9(+) | 1194.8 |
| | mouse#3 | 4 | MNP9(+) | 581.02 |
| | mouse#3 | 5 | MNP9(+) | 581.54 |
| | mouse#3 | 6 | MNP9(+) | 793.6 |
| | mouse#3 | 7 | MNP9(+) | 944.78 |
| | mouse#3 | 8 | MNP9(+) | 919.17 |
| | mouse#3 | 1 | MNP9(+) | 901.88 |
| | mouse#3 | 2 | MNP9(+) | 1058.71 |
| | mouse#3 | 3 | MNP9(+) | 761.41 |
| | mouse#3 | 4 | MNP9(+) | 1367.04 |
| | mouse#3 | 5 | MNP9(+) | 1073.18 |
| | mouse#3 | 6 | MNP9(+) | 1172.79 |
| | mouse#3 | 7 | MNP9(+) | 825.47 |
| | | | MEAN | 885.57425 |
| | | | SEM | 22.12 |

**Fig. S7B (e)** Data Source File

| | | |
|---|---|---|
| Test for normal distribution | | |
| Shapiro-Wilk test | | |
| W | 0.9817 | 0.9795 |
| P value | 0.2486 | 0.1039 |
| Passed normality | Yes | |
| P value summary | ns | |
| Number of values | 88 | 105 |
| Unpaired t test | | |
| P value | 0.121 | |
| P value sum | ns | |
| Significantly diff | No | |
